# Mitochondrial control of glycerolipid synthesis by a PEP shuttle

**DOI:** 10.64898/2025.12.12.693842

**Authors:** Tadashi Yamamuro, Daisuke Katoh, Guilherme Martins Silva, Hiroshi Nishida, Satoshi Oikawa, Yusuke Higuchi, Dandan Wang, Masanori Fujimoto, Naofumi Yoshida, Mark Li, Jihoon Shin, Zezhou Zhao, Jin-Seon Yook, Lijun Sun, Shingo Kajimura

## Abstract

Mitochondria provide a variety of metabolites, in addition to ATP, to meet cell-specific needs. One such metabolite is phosphoenolpyruvate (PEP), which contains a higher-energy phosphate bond than ATP and has diverse biological functions. However, how mitochondria-generated PEP is delivered to the cytosol and fulfills cell-specific requirements remains elusive. Here, we show that SLC25A35 regulates mitochondrial PEP efflux and glyceroneogenesis in lipogenic cells that utilize the pyruvate-to-PEP bypass. Reconstitution and structural studies demonstrated PEP transport by SLC25A35 in a pH gradient-dependent manner. Loss of SLC25A35 in adipocytes impaired the conversion of mitochondrial PEP into glycerol-3-phosphate, thereby reducing glycerolipid synthesis. Significantly, hepatic inhibition of SLC25A35 in obese mice alleviated steatosis and improved systemic glucose homeostasis. Together, these results suggest that mitochondria facilitate glycerolipid synthesis by providing PEP via SLC25A35, offering lipogenic mitochondria as a target to limit glycerolipid synthesis, a pivotal step in the pathogenesis of hepatic steatosis and Type 2 diabetes.

## INTRODUCTION

Mitochondria serve as metabolic hubs that provide ATP, as well as essential metabolites, to various subcellular compartments, including the nucleus, endoplasmic reticulum, and lipid droplets ^1,2^. A prime example linked to mitochondrial activity is phosphoenolpyruvate (PEP). PEP harbors one of the highest energy phosphate bonds in cells, with a standard Gibbs free energy change (ΔG°’) of approximately −61.9 kJ mol^-1^ for the hydrolysis of PEP to pyruvate and inorganic phosphate, which is substantially higher than for ATP to ADP (−30.5 kJ mol^-1^). This very large negative ΔG°’ makes PEP a versatile energy donor in a variety of metabolic processes. In glycolysis, it drives ATP production via pyruvate kinase; in gluconeogenesis, PEP is an essential substrate in the liver; and in glyceroneogenesis, PEP is converted to glycerol-3-phosphate (G3P), the glycerol backbone necessary for the esterification of fatty acids into glycerolipids, including triglycerides and phospholipids in adipose tissue and liver ^3–7^. Furthermore, PEP contributes to ATP-dependent insulin secretion in pancreatic β-cells ^7–13^ and the regulation of Ca²⁺ signaling in activated T cells ^14^. How such an energetically high molecule is generated, delivered, and meets cell-type-specific demands remains elusive.

To regenerate this high-energy-containing PEP from pyruvate, cells must overcome the energetic barrier by two-step reactions: the first step is an ATP-consuming reaction to convert pyruvate to oxaloacetate (OAA) by pyruvate carboxylase (PC), and the second step is a GTP-consuming reaction to convert OAA to PEP by phosphoenolpyruvate carboxykinase (PEPCK) ^15,16^. Of note, PEP can be synthesized from OAA either in the cytosolic compartment via PEPCK (C-PEPCK or PCK1) or within the mitochondria by PCK2 (also known as M-PEPCK) (**Figure 1A**). Historically, much attention has been paid to PCK1 in the context of gluconeogenesis because a) PCK1 is more abundant than PCK2 in the liver of rodents ^3^ (note: human PCK2 accounts for approximately 50%), b) PCK1 expression and activity are dynamically regulated by nutritional and hormonal cues, including fasting ^3^, c) overexpression of PCK1 in the liver caused hyperglycemia ^17,18^, and 4) inhibition of PCK1 attenuated hepatic gluconeogenesis from lactate/pyruvate, although liver-specific *Pck1* knockout mice maintain euglycemia due to compensatory renal gluconeogenesis ^19–24^. Thus, it is reasonable to consider that PEP is generated primarily in the cytosol.

**Figure 1.**
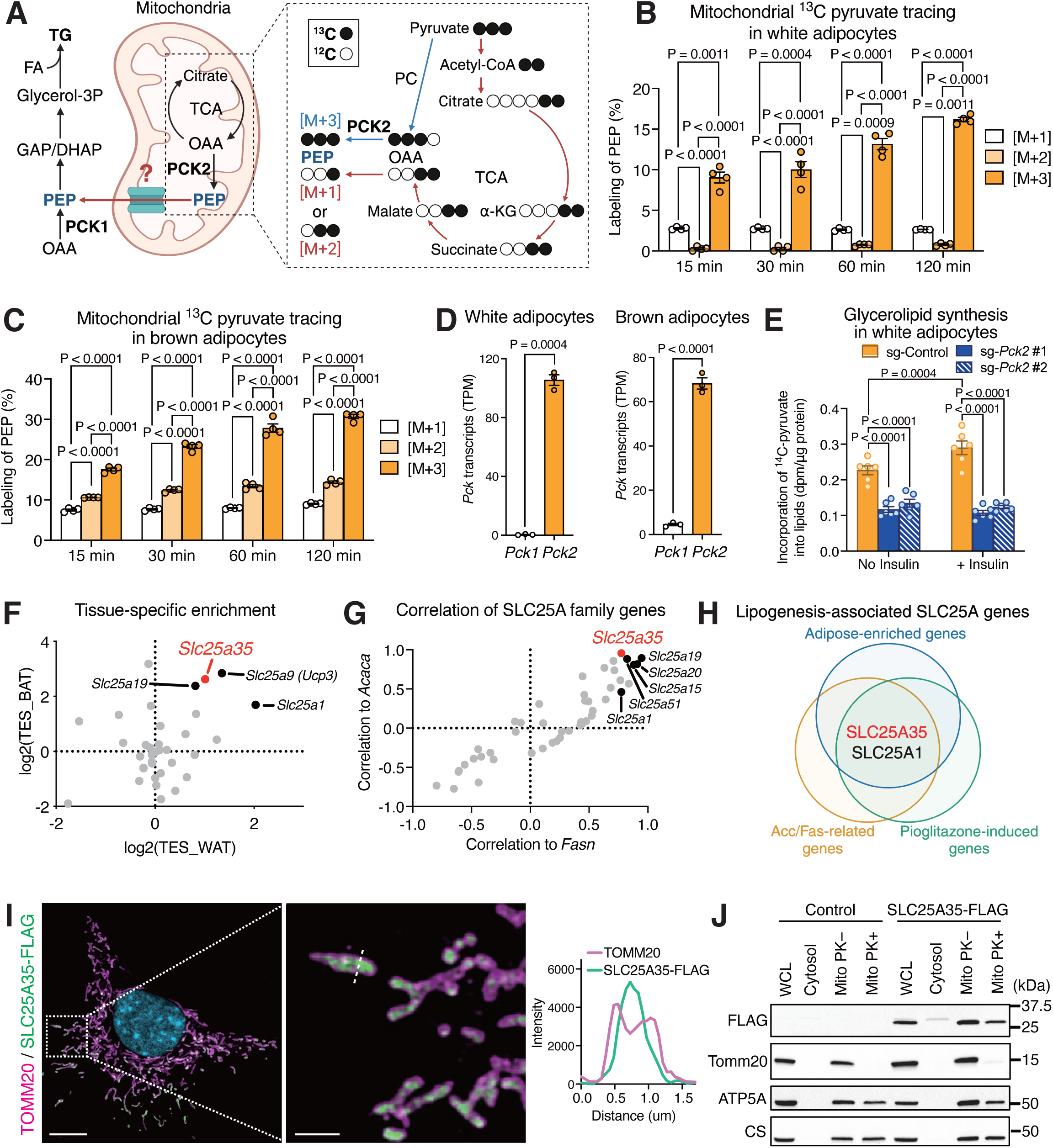
Identification of carrier proteins linking to mitochondrial PEP synthesis. (A) Mitochondrial ^13^C-pyruvate generates M+3 PEP by PC and PCK2 or M+1/M+2 PEP through TCA-cycle oxidation. (B) White adipocyte-derived mitochondria incubated with ^13^C-pyruvate (10 mM) for the indicated time for detecting labeled PEP. n = 4. (C) Brown adipocyte-derived mitochondria incubated with ^13^C-pyruvate (10 mM) for detecting labeled PEP. n = 4. (D) TPM values for *Pck1* and *Pck2* in white and brown adipocytes at day 4 (GSE173710). n = 3. (E) 1-^14^C pyruvate incorporation to glycerolipids in white adipocytes transduced with *Pck2* sgRNAs in the presence or absence of insulin. n = 6. (F) Tissue-enrichment score (TES) of SLC25A genes in mouse WAT and BAT (GSE152382). n = 3–6. (G) Correlation of SLC25A genes with Acaca and Fasn across six mouse white adipocyte subpopulations (GSE176171). (H) Venn diagram showing common SLC25A genes to adipose-enriched genes (log_2_(TES_BAT), log_2_(TES_WAT) > 0.45), *Acaca*/*Fasn*-related genes (r(*Acaca*), r(*Fasn*) > 0.45), and pioglitazone-induced genes (log_2_FC > 0.45, -log_10_(P-value) > 2, Figure S2F). (I) Representative immunofluorescence image of white preadipocytes with SLC25A35-FLAG, TOMM20, and DAPI. Scale bars, 10 μm (left) and 2 μm (right). (J) Proteinase K assay in mitochondria from control (wild-type) and SLC25A35-FLAG expressing cells.

From a bioenergetic point of view, however, PEP synthesis in the mitochondrial compartment appears more efficient than the cytosolic pathway, considering the availability of mitochondrial GTP and the need for fewer enzymatic steps and metabolite transports ^25^. In addition, mitochondria contain abundant amounts of PEP at concentrations of 10 µM and higher ^26,27^. Notably, when we performed ^13^C_3_-labeled pyruvate tracing experiments in isolated mitochondria of white adipocytes, M+3 PEP was the dominant form relative to M+1 or M+2 PEP (**Figure 1B**). Furthermore, the mitochondria from brown adipocytes generated significantly higher levels of M+3 PEP than M+1 or M+2 PEP (**Figure 1C**). Additionally, the mitochondria from white preadipocytes dominantly generated M+3 PEP from ^13^C_3_-labeled pyruvate (**Figure S1A**). The oxidative TCA cycle was active in the mitochondria of brown adipocytes, although less activities were found in white adipocytes and preadipocytes (**Figure S1B-D**). These results suggest that the conversion from pyruvate → OAA → PEP, *a.k.a.* the pyruvate-to-PEP bypass, is the primary pathway of PEP synthesis in adipocyte mitochondria regardless of their oxidative TCA cycle activities. In contrast, mitochondria from C2C12 myoblasts and HEK293T cells generated M+1 or M+2 PEP, via the oxidation of ^13^C-pyruvate in the TCA cycle, at higher levels than M+3 PEP (**Figure S1E, F**), suggesting that mitochondria regenerate PEP in a cell-type-selective manner.

These findings raise a critical question: How is mitochondrial-generated PEP transported into the cytosolic compartment? Since the inner mitochondrial membrane (IMM) is impermeable to metabolites, the export of PEP from the mitochondrial matrix to the cytosol necessitates specific mitochondrial carrier proteins. Accordingly, this study aimed to identify an as-of-yet uncharacterized mitochondrial carrier that mediates PEP transport across the IMM and further investigate its biological significance.

## RESULTS

### Mitochondrial carrier proteins that are linked to PEP synthesis

The cytosolic PCK1 has been recognized as the primary PEPCK; however, previous studies reported that the depletion of PCK2 in rat liver reduces gluconeogenesis from lactate, pyruvate, and amino acids ^25,28^. Furthermore, PCK2 is highly expressed in pancreatic islets, where mitochondria-derived PEP, generated by PCK2 using mitochondrial GTP, significantly contributes to the total PEP pool and controls glucose-stimulated insulin secretion ^7–13^. Intriguingly, PCK2 was the dominant PEPCK in inguinal white adipose tissue (WAT)-derived adipocytes, whereas PCK1 expression was low (**Figure 1D, Figure S2A**). Similarly, brown adipocytes expressed higher *Pck2* transcripts than *Pck1*. Next, we deleted *Pck1* or *Pck2* by the CRISPR-Cas9 system and examined the function in lipogenesis (**Figure S2B**). To assess glycerolipid synthesis in adipocytes, we used 1-¹⁴C pyruvate as a tracer, which bypasses direct labeling of glyceraldehyde-3-P (GA3P) and G3P from glucose via glycolysis. In addition, labeling at the first carbon ensures that the ¹⁴C is released as CO₂ during pyruvate decarboxylation, thereby minimizing incorporation of ¹⁴C into mitochondrial acetyl-CoA and downstream fatty acid synthesis (**Figure S2C**). To experimentally validate this assumption, we extracted total lipids from adipocytes incubated with 1-^14^C pyruvate, saponified the lipids, and quantified ¹⁴C in the fatty acid fraction by thin-layer chromatography (TLC). ¹⁴C signals in the fatty acid fraction were indistinguishable from background levels, indicating no detectable incorporation of ¹⁴C into the fatty acyl moiety of glycerolipids (**Figure S2D**). Hence, the assay primarily measures ^14^C incorporation into the glycerol backbone of glycerolipids. Insulin promoted 1-^14^C incorporation into the lipid fraction in adipocytes, validating the assay. We found that adipocytes lacking *Pck2,* induced by two distinct gRNAs, showed a significant decrease in glycerolipid synthesis relative to control cells (**Figure 1E**). In agreement, incorporation of ^14^C-labeled glucose into the lipid fraction was significantly reduced in adipocytes lacking *Pck2,* reflecting the contribution of PCK2 to lipogenesis, although this assay cannot differentiate whether the tracer is incorporated into the glycerol backbone or the fatty acyl moiety of glycerolipids (**Figure S2E**). On the other hand, adipocytes with sg*Pck1* retained the ability to synthesize lipids from ^14^C-labeled glucose. These results suggest that mitochondrial PEP generated via PCK2 contributes to glycerolipid synthesis in adipocytes. The results also suggest the existence of an as-yet-unidentified mitochondrial carrier that mediates PEP export in these cells.

To identify such mitochondrial carriers, we performed the following analyses: First, we searched for genes encoding mitochondrial carrier proteins that were enriched in brown and white adipocytes, which primarily generated M+3 PEP compared to other cell types. This analysis identified several mitochondrial carrier proteins, including *Slc25a1* (mitochondrial citrate carrier), *Slc25a9* (also known as UCP3), *Slc25a19* (thiamine carrier), and *Slc25a35* (**Figure 1F**). Second, we searched for genes that showed positive correlations with *Acaca* (acetyl-CoA carboxylase alpha) and *Fasn* (fatty acid synthase), the two representative genes that control lipogenesis ^29^. The analyses identified several mitochondrial carrier proteins, including *Slc25a1, Slc25a15* (ornithine carrier), *Slc25a19* (thiamine carrier), *Slc25a20* (carnitine/acylcarnitine carrier), *Slc25a35,* and *Slc25a51* (NAD carrier) (**Figure 1G**). Third, we searched for genes that were induced by lipogenic stimuli, especially synthetic agonists of peroxisome proliferator-activated receptor-gamma (PPARγ), such as troglitazone, rosiglitazone, pioglitazone, and AG035029, in rat adipose tissue (**Figure S2F**). These analyses identified SLC25A1 and SLC25A35 that met the three criteria outlined above (**Figure 1H**). While SLC25A1 is known to export citrate ^30–32^, SLC25A35 is currently an orphan carrier that is expressed abundantly in adipocyte populations of adipose tissue (**Figure S2G**). At the single-cell resolution, *Slc25a35* is co-expressed with *Acaca* and *Fasn* across all adipocyte populations, with particular enrichment in a population (mAd5) that displays the high lipogenic capacity ^33^ (**Figure S2H**). These observations motivated us to further examine the function of SLC25A35.

Although many members of the SLC25 family are expected to localize to the inner mitochondrial membrane (IMM), SLC25A17 resides in peroxisomes, while SLC25A46, SLC25A49, and SLC25A50 localize to the outer mitochondrial membrane (OMM) ^34–36^. Hence, we next examined the subcellular localization of SLC25A35 in preadipocytes expressing SLC25A35 with a Flag-tag at its C-terminus (SLC25A35-Flag). High-resolution microscopy revealed that the SLC25A35 protein was expressed selectively in the mitochondria, but the signal did not overlap with the OMM marker TOMM20 (**Figure 1I**). Rather, SLC25A35 was localized in the IMM as proteinase K treatment degraded TOMM20, whereas SLC25A35 and an IMM-localized protein, ATP5A, remained intact (**Figure 1J**). Additionally, we validated the mitochondrial localization of the SLC25A35 protein in differentiated adipocytes (**Figure S2I, J**).

### SLC25A35 is required for mitochondrial PEP efflux

To examine the biological role of SLC25A35, we generated inguinal WAT-derived preadipocytes lacking SLC25A35 (*Slc25a35* KO cells) and control cells by the *Easi*-CRISPR (Efficient additions with ssDNA inserts-CRISPR) system ^37^ (**Figure 2A**). We then rapidly isolated mitochondria from these cells using Mito-Tag ^38^, and analyzed their metabolites by liquid chromatography-mass spectrometry (LC-MS) (**Figure 2B, Figure S3A**). We found that *Slc25a35* KO cell-derived mitochondria contained higher levels of several metabolites than control cell-derived mitochondria, including PEP, NAD^+^ (detected by positive and negative ion modes), and 1,3-bisphosphoglycerate (BPG) (**Figure 2C, Table S1**). In contrast, we did not observe substantial changes in these metabolites at the whole-cell level, whereas *Slc25a35* KO cells exhibited modestly elevated levels of TCA intermediates and α-ketoisovalerate (KIV) compared to control cells (**Figure 2D, Table S2**). When mitochondrial metabolites were normalized by whole-cell metabolites to calculate mitochondrial enrichment, we found that PEP was the most enriched metabolite (9.4-fold enrichment) in the mitochondria of *Slc25a35* KO cells relative to control cells (**Figure 2E, Table S3**). There was no change in ATP and GTP levels between the two groups. In differentiated adipocytes, PEP was the only metabolite that showed significant enrichment in *Slc25a35* KO mitochondria among the metabolites detected (**Figure S3B-D**). These results suggest that elevated PEP levels in *Slc25a35* KO mitochondria reflected mitochondrial accumulation of PEP rather than an increase in total PEP amounts, while the changes in other metabolites were due to alterations occurring in whole-cell levels.

**Figure 2.**
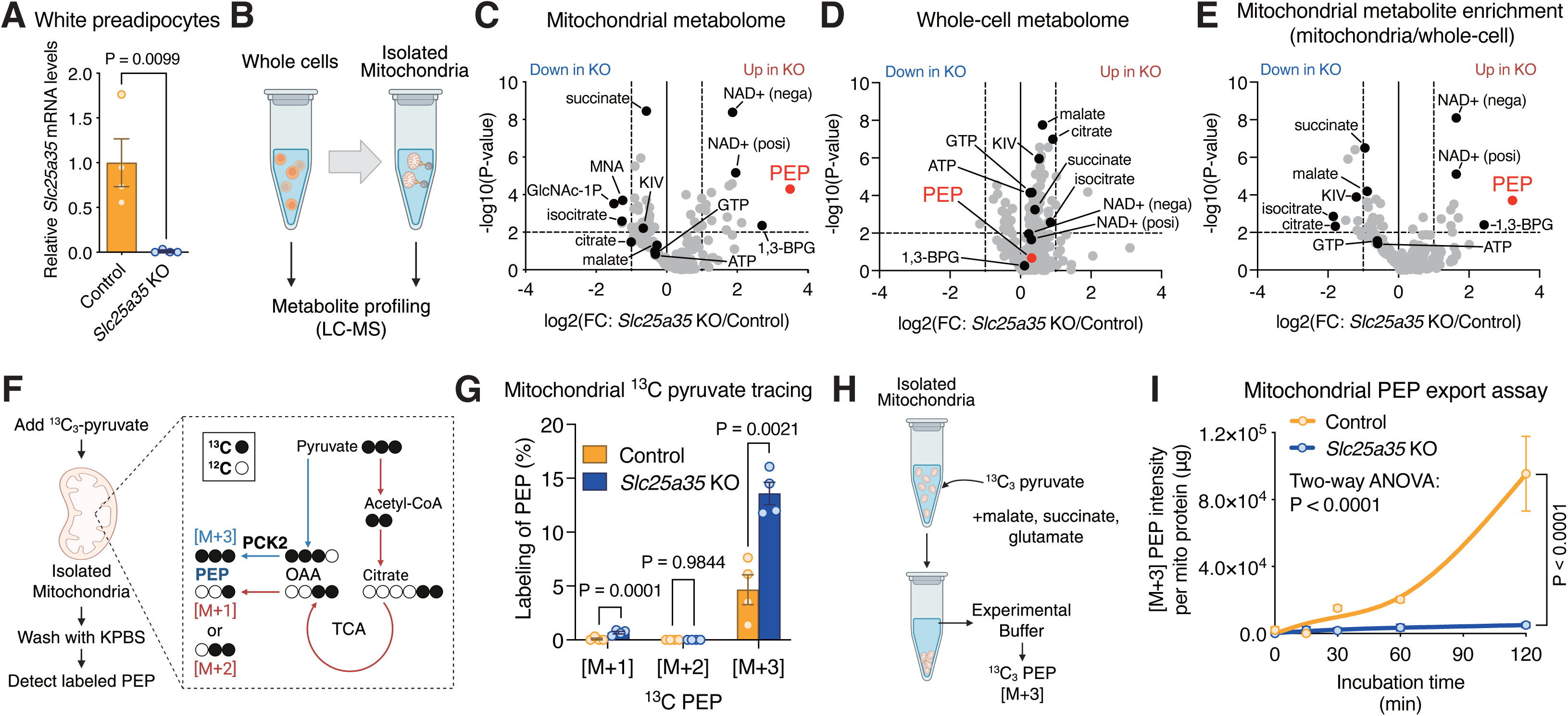
SLC25A35 is required for mitochondrial PEP export. (A) Relative mRNA level of *Slc25a35* in control and *Slc25a35* KO white preadipocytes. n = 4. (B) Schematic of mitochondrial and whole-cell metabolomics using MITO-Tag. (C) Mitochondrial metabolomics in white preadipocytes. The intensity was normalized to protein levels. n = 7. (D) Whole-cell metabolomics in white preadipocytes. n = 7. (E) Mitochondrial metabolite enrichment in (C, D). n = 7. (F) Schematic of mitochondrial ^13^C-pyruvate tracing. (G) ^13^C-pyruvate tracing (2 mM, 5 min) in mitochondria from control or *Slc25a35* KO white preadipocytes. n = 4. (H) Schematic of mitochondrial PEP export assay with ^13^C-pyruvate. (I) Mitochondrial M+3 PEP export in white preadipocytes. The values were normalized by mitochondrial protein levels. n = 4.

Next, we performed ^13^C tracing experiments in isolated mitochondria from control and *Slc25a35* KO cells, incubating mitochondria with ^13^C_3_-pyruvate for 5 min, quickly washing twice, and analyzing labeled metabolites by LC-MS (**Figure 2F**). We found that mitochondria from *Slc25a35* KO cells contained significantly higher levels of M+3 PEP than those from control cells, while there was no difference in M+2 PEP levels between the two (**Figure 2G**). While M+1 PEP was slightly elevated in *Slc25a35* KO cells, M+3 PEP was by far the dominant form in white preadipocyte-derived mitochondria. The accumulation of M+3 PEP in the mitochondria of *Slc25a35* KO cells occurred in a time-dependent manner (**Figure S3E**). These results suggest that the accumulated PEP in the mitochondria is largely derived from M+3 OAA by the action of PC and PCK2, rather than OAA derived through oxidation of pyruvate in the TCA cycle. Supporting this, we found that pyruvate oxidation in the TCA cycle was largely unchanged in *Slc25a35* KO cells, as there was no difference in ^13^C-labeled succinate and citrate levels in whole cells (**Figure S3F, G**).

Similar results were observed in isolated mitochondria, where the total oxidative TCA cycle activity remained largely unchanged between control and *Slc25a35* KO cells, despite a modest reduction in M+3 citrate levels (**Figure S3H-K**). Additionally, we found no differences in PDH phosphorylation, PDH, and PC protein expression between the two groups (**Figure S3L**). These results indicate that SLC25A35 deletion leads to mitochondrial accumulation of OAA-derived PEP due to reduced PEP efflux, rather than reduced oxidative TCA cycle activity or changes in protein expression of PC and PDH.

To test this possibility, we next developed an experimental system to directly measure mitochondrial PEP export (**Figure 2H**). In brief, isolated mitochondria from preadipocytes were incubated with ^13^C-pyruvate in media containing malate, succinate, and glutamine to energize the mitochondria. Subsequently, we harvested the incubation media at 15 min, 30 min, 60 min, and 120 min following incubation, in which we detected M+3 PEP in the media. We found that isolated mitochondria actively secreted ^13^C-labeled PEP in a time-dependent manner. However, mitochondrial PEP export was substantially reduced in *Slc25a35* KO mitochondria (**Figure 2I**). The data suggest that active PEP export from mitochondria requires SLC25A35.

### Reconstitution of PEP transport by SLC25A35

To reconstitute PEP transport by SLC25A35, we developed a cell-free liposome system with purified SLC25A35 protein (**Figure 3A**). In brief, Sf9 cells infected with baculovirus expressing either a mouse SLC25A35-TwinStrep construct or an empty vector (control) were subjected to affinity purification (**Figure 3B**). Purified SLC25A35 protein or an empty-vector control eluate was subsequently inserted into liposomes. We excluded background signals from empty liposomes without proteins, as these signals represent non-specific binding to liposome membranes. In the SLC25A35 liposomes preloaded with unlabeled PEP, we observed significantly higher transport of ^13^C-labeled PEP than in control liposomes (**Figure 3C**). To determine the transport specificity, we next tested ^13^C-PEP transport in the presence of excess unlabeled PEP as a competitor. We found that ^13^C-PEP transport into SLC25A35 liposomes was completely abolished, supporting SLC25A35’s substrate specificity for PEP. This PEP transport was consistently observed at various PEP concentrations inside and outside the liposomes (**Figure 3D**). Additionally, partial digestion of the proteo-liposomes with proteinase K significantly reduced the PEP transport activity (**Figure S4A**).

**Figure 3.**
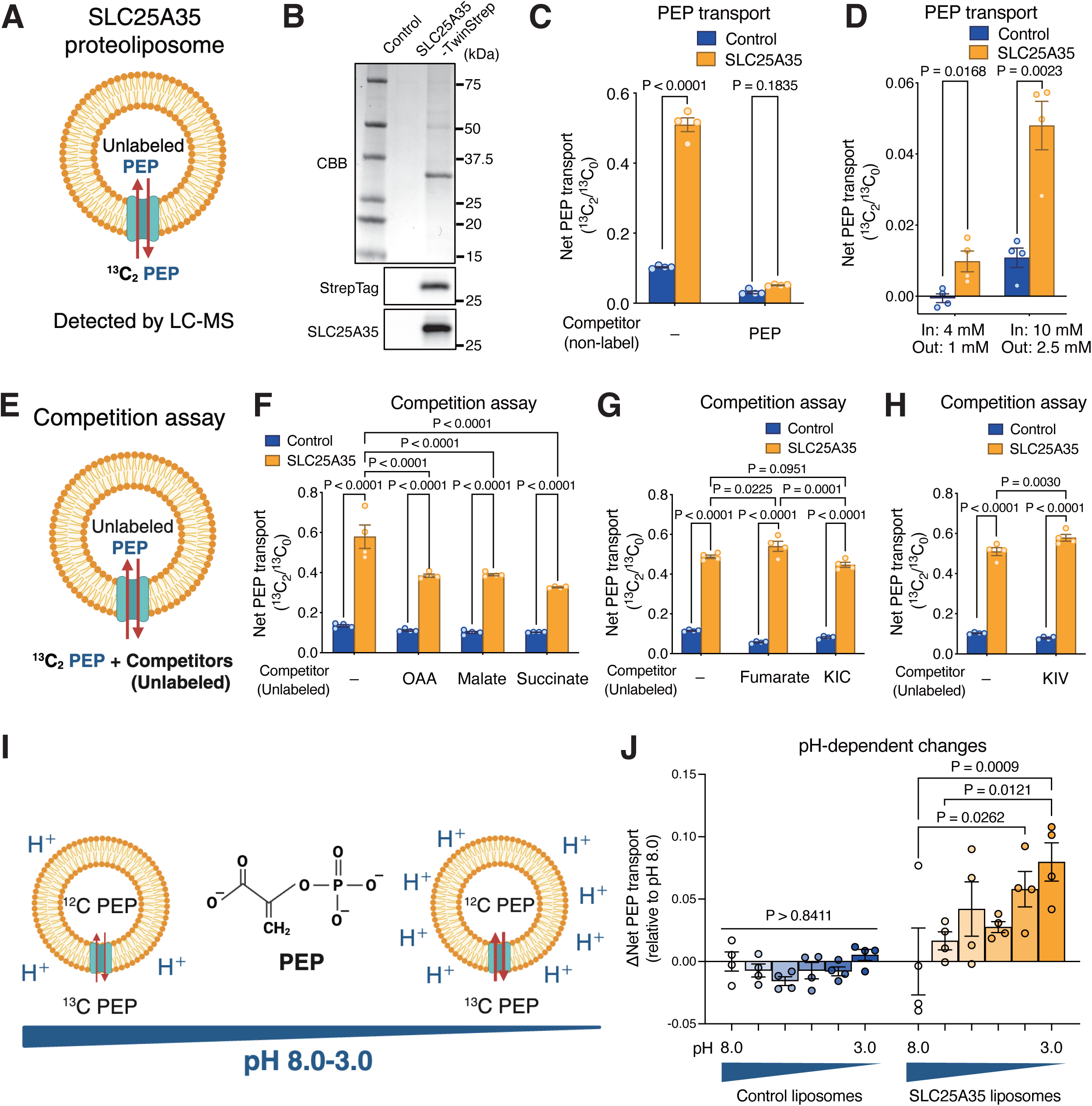
Reconstitution of PEP transport by SLC25A35. (A) Schematic of PEP transport assays in proteo-liposomes with purified SLC25A35 protein or control. (B) Purified SLC25A35 protein or an empty-vector control eluate visualized by CBB staining. (C) PEP transport assay with or without excess unlabeled PEP (50 mM) as a competitor. n = 4. (D) PEP transport assay with the indicated intra- and extra-liposomal PEP concentrations. n = 4. (E) Schematic of the competition assays with an excess of unlabeled metabolites (50 mM). (F) Competition assays using indicated metabolites as in (E). n = 4. (G) Competition assays using indicated metabolites. n = 4. (H) Competition assays using indicated metabolites. Non-competitor samples are identical to those in (C). n = 4. (I) Schematic of pH dependence in PEP transport. (J) pH-dependent PEP transport. ΔNet PEP transport was calculated by subtracting the mean value at pH 8.0 from each value at the corresponding pH. n = 4.

To further determine the substrate specificity, we next measured the transport of ^13^C-PEP in SLC25A35 liposomes in the presence of unlabeled OAA, malate, succinate, and fumarate as competitors (**Figure 3E**). While the ^13^C-PEP transport was consistently observed in proteo-liposomes containing SLC25A35, there was a modest but significant inhibition of ^13^C-PEP transport in the presence of OAA, malate, and succinate (**Figure 3F**). On the other hand, an excess amount of fumarate did not block ^13^C-PEP transport in SLC25A35 liposomes (**Figure 3G**). These results suggest that PEP is a high-affinity substrate for SLC25A35, while other TCA cycle intermediates, such as OAA, malate, and succinate, could be transported via SLC25A35, albeit with lower affinity. A previous work in *Saccharomyces cerevisiae* reported that a mitochondrial OAA carrier (Oac1p) transports α-isopropylmalate (α-IPM), which is derived from α-ketoisovalerate (KIV), and that ectopic expression of human SLC25A35 partially restored the growth defect of *Oac1p*-deficient yeast ^39^. Our metabolomics study also found reduced enrichment of KIV in *Slc25a35* KO mitochondria, indicating the possibility that KIV and α-ketoisocaproate (KIC) could be substrates for SLC25A35. Our liposome-based substrate competition assays showed that excess unlabeled KIV or KIC did not compete with ^13^C-PEP uptake in liposomes containing SLC25A35 (**Figure 3G, H**). When ^13^C-labeled KIV and KIC were used as tracers, we did not observe active transport in SLC25A35-containing liposomes (**Figure S4B**). These results suggest that reduced enrichment of KIV in *Slc25a35* KO mitochondria is unlikely to result from reduced import into mitochondria.

PEP carries a net negative charge under physiological conditions. This raises the possibility that mitochondrial PEP transport is coupled with the proton gradient across the membrane, *i.e.,* mitochondrial membrane potential. To test this, we prepared proteo-liposomes under conditions ranging from pH 3.0 to pH 8.0 to determine the extent to which PEP transport depends on the pH gradient (**Figure 3I**). We found that PEP transport was significantly enhanced at lower pH conditions in SLC25A35 liposomes, but not in control liposomes (**Figure 3J**). The data suggest that mitochondrial PEP transport can be facilitated when mitochondrial membrane potential is high.

### Structural insights into SLC25A35-mediated PEP transport

Next, we applied computational simulations to further assess the ability of SLC25A35 to transport PEP as a substrate. There are currently no reported three-dimensional (3D) protein structures for SLC25A35, and its low sequence similarity to other template structures (<30% of identity by BLASTp search) renders homology modeling a less reliable approach. Thus, we utilized the AlphaFold structure ^40^ of human SLC25A35 (AF-Q3KQZ1-F1-model_v4) to evaluate PEP as a substrate. This structure shows overall high-confidence prediction scores (**Figure S5A**) and resembles a shape that is common to solute carriers, such as a channel-like central cavity formed by six transmembrane alpha helices perpendicular to the inner mitochondrial membrane ^41^ (**Figure 4A, Figure S5B, C**). In addition, the inner walls of this cavity are lined with polar and basic amino acids, including Y72, Q73, N77, R80, Y124, K127, R175, and R276, all of which are conserved in mouse SLC25A35 (**Figure 4B**).

**Figure 4.**
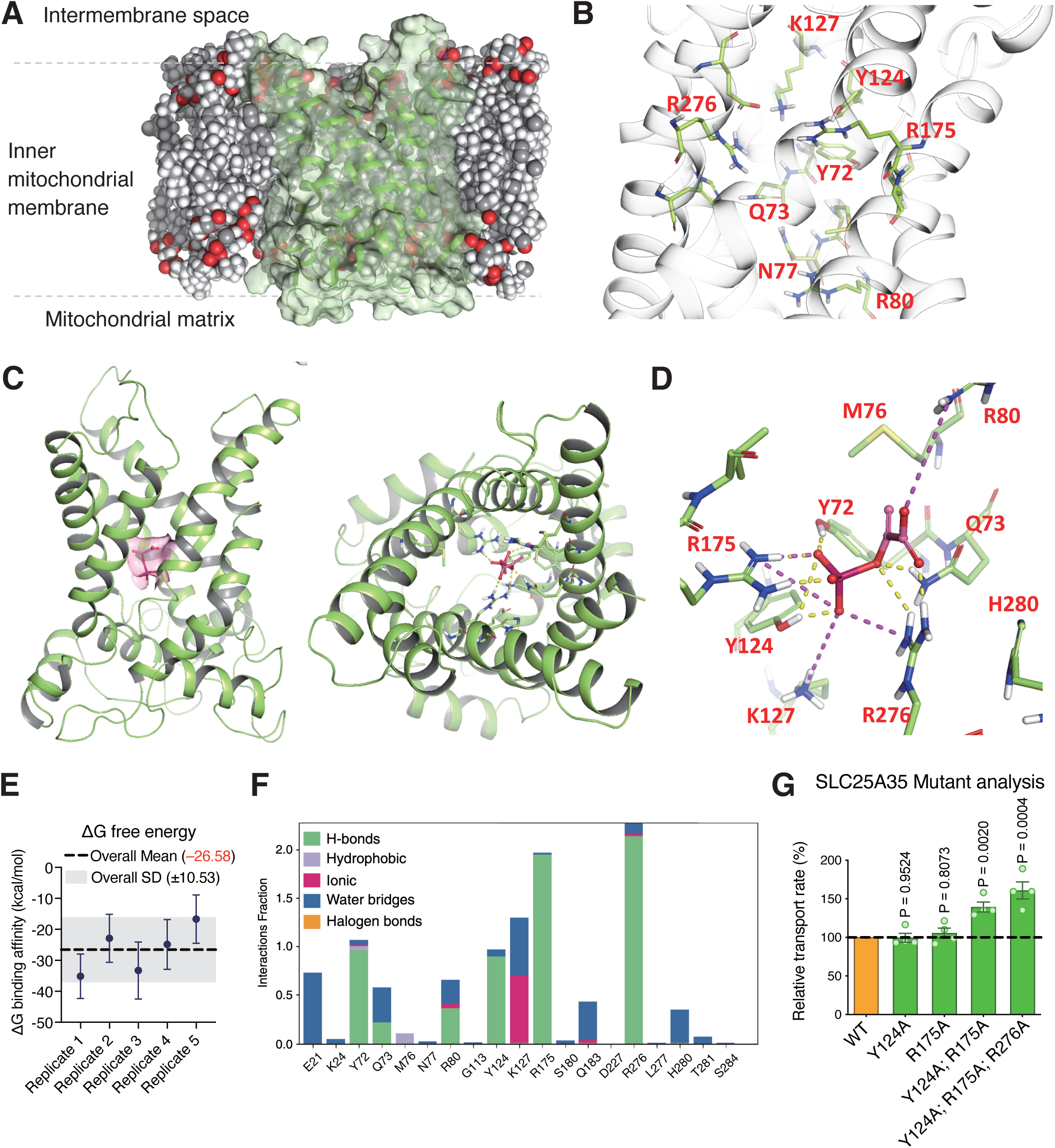
Structural insights into SLC25A35-mediated PEP transport. (A) Side view of the SLC25A35 structure at the inner mitochondrial membrane. (B) Key amino acid residues (green sticks) at the detected SLC25A35 central channel-like cavity. (C) Left: Side view of PEP (magenta carbons and surface) within the SLC25A35 binding site. Right: Top view of PEP within the SLC25A35 binding cavity. (D) Close-up 3D view of PEP (magenta carbons; ball-and-stick representation) highlighting key interactions with SLC25A35 residues observed after stabilization in one replicate of molecular dynamics (MD) simulation. Dashed yellow and magenta lines indicate hydrogen bonds and salt bridges, respectively. (E) Molecular mechanics with generalized MMGBSA ΔG binding affinity estimation from five replicate MD simulations of the PEP–SLC25A35 complex at the binding site. (F) Interaction bar chart for the PEP-SLC25A35 MD replicate simulations, showing per residue interaction fractions. (G) Relative PEP transport activity of wild-type and indicated SLC25A35 mutants in proteo-liposomes. Relative PEP transport was normalized to the corresponding wild-type activity in each assay. Original data and P values are shown in Figure S6F, G. n = 4.

We next performed induced-fit docking to generate a stable and low-energy interaction between PEP and SLC25A35 in which PEP adopted a conformation that was accommodated by the SLC25A35 central cavity (**Figure 4C, Figure S6A**). At this state, the following interactions were observed: hydrogen bonds and/or salt bridge between the phosphate oxygens of PEP and residues Q73 and R276 of SLC25A35, as well as hydrogen bonds between the carboxylate of PEP and residues R175, Y124, and Y72 of SLC25A35 (**Figure 4D, Figure S6B**). Subsequently, we performed molecular dynamics (MD) studies considering SLC25A35 orientation at the mitochondrial inner membrane, in the absence and presence of PEP. This allowed us to infer that PEP forms a stable, reproducible complex with SLC25A35. The average Root Mean Square Deviation (RMSD) value for the SLC25A35-PEP interaction was 1.73 Å, which is lower than the 2.10 Å observed in the corresponding *apo* simulation (**Figure S6C**). Importantly, the average ΔG binding affinity (MMGBSA) estimated for the PEP-SLC25A35 interaction was equal to −26.58 ± 10.53 kcal/mol (**Figure 4E**), and the key interactions were kept throughout trajectories, in particular the hydrogen bonds with R276 and R175 (**Figure 4F, Figure S6D, E**).

To determine the functional requirements of the key residues for PEP transport, we generated four mouse SLC25A35 mutants at Y124, R175, R276, and their combination, based on the above structural prediction. Subsequently, we developed proteo-liposomes containing each mutant and tested the involvement of these residues for PEP transport. We found that single SLC25A35 mutants (Y124A, R175A) exhibited PEP transport activity similar to that of the wild-type form. Unexpectedly, the double mutant (Y124A/R175A) and triple mutant (Y124A/R175A/R276A) showed higher PEP transport than the wild-type form (**Figure 4G, Figure S6F, G**). It is notable that a similar observation has been described for SLC25 carriers, such as the ADP/ATP carrier, where weakening the cytoplasmic salt-bridge network by mutating key residues lowers the conformational energy barrier between the cytoplasmic (c) and matrix (m) states, thereby altering the kinetics of substrate access, *i.e.,* leaky gate ^42^. In reconstituted liposomes under saturating substrate conditions, where assays read out Vmax, such gate-loosening substitutions could lead to higher apparent transport than wild-type. Nonetheless, these results support the *in silico* prediction that Y124, R175, and R276 are functionally important for SLC25A35-mediated PEP transport.

### PEP transport via SLC25A35 is required for glyceroneogenesis in adipose tissue

In the cytosolic compartment of adipocytes, PEP can be converted to GA3P and dihydroxyacetone phosphate (DHAP), and subsequently G3P. G3P serves as the glycerol backbone necessary for the esterification of fatty acids into glycerolipids, including triglycerides, diglycerides, and phospholipids in adipose tissue and the liver (**Figure 5A**). The synthesis of G3P from pyruvate, a process known as glyceroneogenesis, contributes to the triglyceride pool in adipose tissue and the liver ^43–45^. Accordingly, we asked the extent to which mitochondria-derived PEP via SLC25A35 contributes to the synthesis of GA3P and DHAP in adipocytes. To this end, we performed ^13^C_3_-pyruvate tracing experiments in inguinal WAT-derived adipocytes lacking *Slc25a35.* The tracing experiment found that *Slc25a35* KO cells produced significantly lower levels of ^13^C-labeled GA3P/DHAP than control cells by 77.2% (**Figure 5B**). Note that GA3P and DHAP are isomers with the same molecular weight, and our LC-MS platform does not distinguish between them; thus, the measured signal represents the combined abundance of GA3P and DHAP in these samples. Importantly, deletion of SLC25A35 led to a significant reduction in G3P levels in inguinal WAT-derived differentiated adipocytes as well as in preadipocytes, albeit at lower levels (**Figure 5C**).

**Figure 5.**
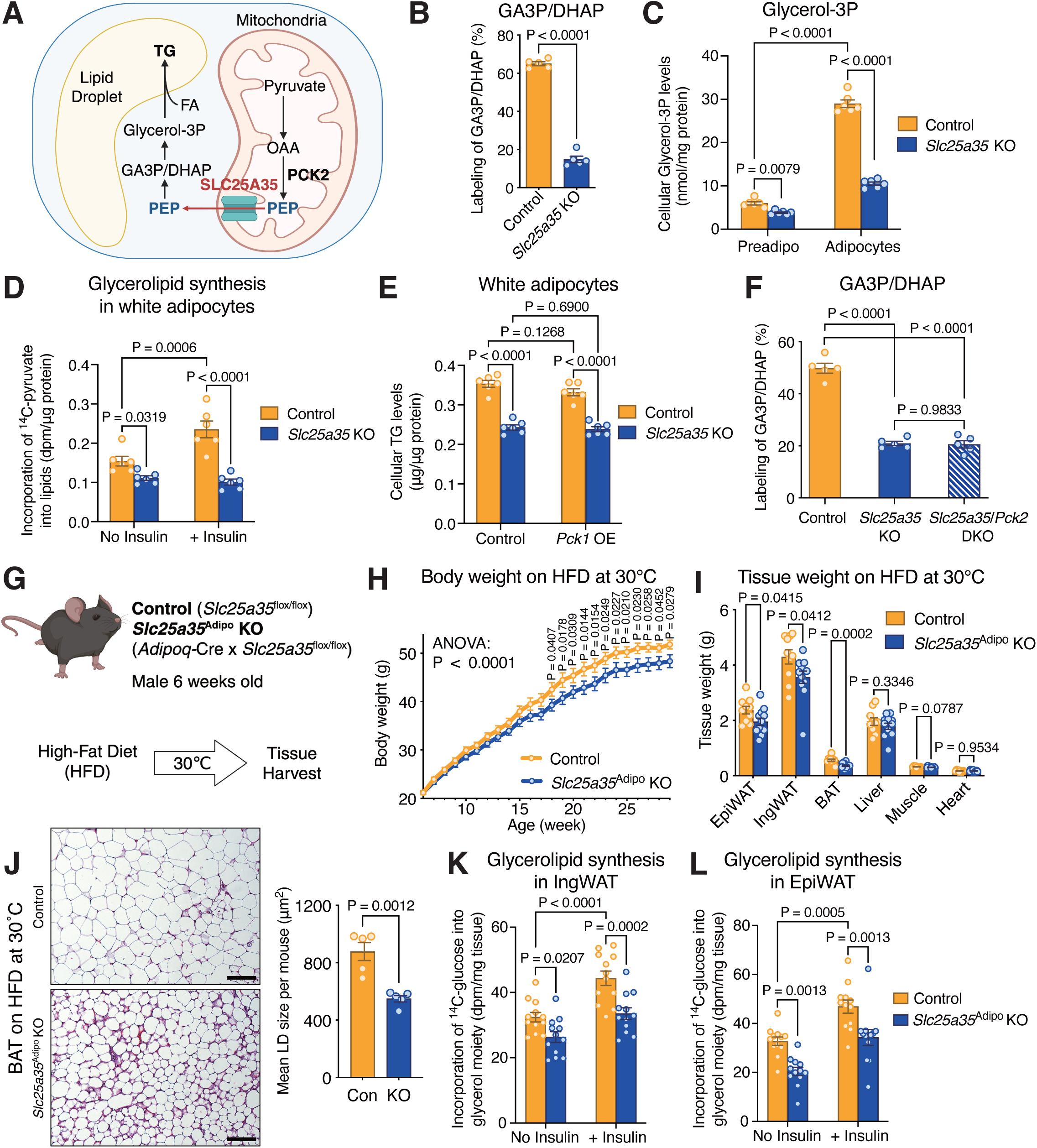
PEP transport via SLC25A35 is required for glyceroneogenesis in adipose tissue. (A) Schematic of glycerolipid synthesis in adipocytes. (B) Whole-cell ^13^C-pyruvate tracing in white preadipocytes for quantifying labeled GA3P/DHAP. n = 5. (C) Cellular glycerol-3P levels in white preadipocytes and differentiated adipocytes normalized by protein levels. n = 6. (D) 1-^14^C-pyruvate incorporation to glycerolipids in white adipocytes with or without insulin. n = 6. (E) Cellular TG levels normalized to protein content in control and *Slc25a35* KO white adipocytes expressing mouse *Pck1* or a control vector. n = 6. (F) Whole-cell ^13^C-pyruvate tracing in control or *Slc25a35* KO preadipocytes transduced with control or *Pck2* #1 sgRNA to quantify labeled GA3P/DHAP. n = 5. (G) Schematic of metabolic studies in fat-specific *Slc25a35* KO mice. (H) Body weight changes of male mice on an HFD at 30°C. n = 10 (control), 12 (KO). (I) Tissue weight of male mice in (H) at 23 weeks of HFD. n = 10 (control), 12 (KO). (J) Representative H&E staining of interscapular BAT from mice in (I). Scale bars, 100 μm. Right: Mean LD size per mouse. n = 5. (K) Glycerolipid synthesis in inguinal WAT (IngWAT) explants of male mice on an HFD at 30°C. Lipids were extracted from explants incubated with ^14^C glucose and saponified. ^14^C radioactivity (dpm) in the glycerol fraction was quantified. n = 12. (L) Glycerolipid synthesis assay in epididymal WAT (EpiWAT) as in (K). n = 12.

To test whether SLC25A35 is required for triglyceride (TG) synthesis in adipocytes, we measured the incorporation of 1-^14^C pyruvate into lipids. As discussed above, tracing studies using 1-^14^C pyruvate in *Slc25a35* KO adipocytes found no detectable ^14^C signals in the fatty acid fraction of glycerolipids (**Figure S7A**). On the other hand, *Slc25a35* KO adipocytes exhibited significantly lower incorporation of ^14^C pyruvate into lipids than control adipocytes by 27.9 % in the absence of insulin, and by 57.1% following insulin stimulation (**Figure 5D**). Consistent results were observed with ^14^C glucose as a tracer (**Figure S7B**). Importantly, total cellular TG contents in *Slc25a35* KO adipocytes were significantly lower than control adipocytes by 31.2% (**Figure 5E**). We then asked whether the total PEP level within a cell is the rate-limiting step for TG synthesis, or if the origin of PEP (*i.e.*, mitochondrial-derived PEP) is the key element. If the former is the case, overexpression of PCK1 could restore TG levels in *Slc25a35* KO adipocytes. However, we found that PCK1 overexpression was insufficient to restore cellular TG levels in *Slc25a35* KO cells (**Figure S7C**). Subsequently, we asked whether SLC25A35 and PCK2 act in the same pathway or independently for glyceroneogenesis by performing ^13^C_3_-pyruvate tracing in cells lacking *Slc25a35* and *Pck2.* In agreement, *Slc25a35* KO cells produced significantly lower amounts of ^13^C-labeled GA3P/DHAP than control cells. Similarly, *Pck2* deletion alone reduced ^13^C-labeled GA3P/DHAP levels (**Figure S7D**). However, deleting *Pck2* did not further attenuate GA3P/DHAP production in *Slc25a35* KO cells (**Figure 5F**). These results suggest that reduced glyceroneogenesis in *Slc25a35* KO cells depends on mitochondria-derived PEP synthesis by PCK2, rather than the total cellular PEP pool.

Since white adipose tissue expresses low levels of glycerol kinase that generates G3P from glycerol ^46,47^, G3P synthesis via glyceroneogenesis is considered the primary pathway for fatty acid esterification in adipose tissue. However, the contribution of mitochondria-derived PEP, as opposed to the cytosol-derived PEP, to the triglyceride pool remains unclear. In this regard, the identification of SLC25A35 as a mitochondrial PEP transporter enables us to rigorously address this question *in vivo*. Hence, we generated *Slc25a35*^flox/flox^ mice and subsequently crossed them with *Adipoq*-Cre to develop fat-specific KO mice (Adiponectin-Cre x *Slc25a35*^flox/flox^, herein *Slc25a35*^Adipo^ KO mice) (**Figure S7E**). To minimize the contribution of adaptive thermogenesis by skeletal muscle and BAT, *Slc25a35*^Adipo^ KO mice and littermate control mice (*Slc25a35*^flox/flox^) were kept under a thermoneutral condition (30°C) on a 60% high-fat diet (HFD) (**Figure 5G**). There was no difference in birth weight between the genotypes; however, male *Slc25a35*^Adipo^ KO mice gained modest but significantly less weight than controls at 18 weeks of age and thereafter (**Figure 5H**). The difference in body weight was primarily due to lower tissue mass of inguinal and epididymal WAT as well as interscapular BAT, but not in the liver or other tissues (**Figure 5I**). Consistently, histological analyses showed that adipocyte size in the interscapular BAT of *Slc25a35*^Adipo^ KO mice was significantly smaller than in control mice (**Figure 5J**). Note that mice were kept at 30°C for 23 weeks, and thus, the interscapular BAT was composed of unilocular adipocytes ^48^. This is notable because previous studies reported active glyceroneogenesis in rat BAT, which expresses high levels of PCK2 ^49–51^. Similarly, adipocyte size in the inguinal WAT of *Slc25a35*^Adipo^ KO mice was significantly smaller than in control mice (**Figure S7F**). We also found that female *Slc25a35*^Adipo^ KO mice exhibited a consistent metabolic phenotype with male mice on a high-fat diet at 30°C (**Figure S7G-J**). When mice were on a normal-chow diet (NCD) at room temperature, there was no difference in body weight (**Figure S7K, L**). It is worth noting that the pyruvate-PEP cycle is suggested to be a futile thermogenic pathway ^52^. However, our data suggest that SLC25A35 in adipose tissue is dispensable for activation of thermogenesis because *Slc25a35*^Adipo^ KO mice could elevate their whole-body energy expenditure at an equivalent level to littermate control mice in response to a β3-adrenergic receptor agonist CL316,243 at 30°C (**Figure S8A**). Furthermore, there were no differences in whole-body energy expenditure, food intake, and locomotor activities between the two groups (**Figure S8B-D**). In addition, we found no difference in adipose tissue lipolysis and glucose oxidation between the genotypes (**Figure S8E-G**).

In contrast, we found a significant difference in glycerolipid synthesis in the adipose tissue. To determine ^14^C glucose incorporation into the glycerol backbone of glycerolipids in adipose tissue, we used saponification of lipids extracted from adipose tissues, followed by the periodic acid–Schiff staining and TLC (PAS–TLC) method ^53^. We found that the inguinal WAT from *Slc25a35*^Adipo^ KO mice exhibited significantly lower ^14^C-glucose incorporation into the glycerol backbone of glycerolipids than in control mice in the absence and presence of insulin (**Figure 5K**). Similarly, the epididymal WAT of *Slc25a35*^Adipo^ KO mice showed lower ^14^C incorporation into the lipid glycerol backbone than control mice (**Figure 5L**). Since ^14^C-labeled glucose can be incorporated into the fatty acyl moiety of glycerolipids, we applied saponified lipids extracted from WAT to TLC analyses. In contrast to the functional requirement of SLC25A35 for glyceroneogenesis, we found no significant differences in ^14^C incorporation into fatty acid fraction between the genotypes (**Figure S8H, I**). These results suggest that mitochondrial PEP export into the cytosolic compartment via SLC25A35 supports glyceroneogenesis, thereby contributing to glycerolipid synthesis in adipose tissue.

### Blockade of mitochondrial PEP export ameliorates hepatic steatosis

Approximately 50% or more of the fatty acids taken up by the liver are re-esterified into triglycerides and released as very low-density lipoproteins (VLDL); a disruption of this process leads to metabolic dysfunction-associated steatotic liver disease (MASLD) ^54,55^. Previous studies have shown that glyceroneogenesis plays a key role in triglyceride synthesis in the livers of both rats and humans, particularly during prolonged fasting ^56,57^. Additionally, glyceroneogenesis seems to contribute to VLDL triglycerides in individuals with Type 2 diabetes ^58^. However, the contribution of mitochondrial-derived PEP to hepatic glyceroneogenesis and its impact on the pathogenesis of hepatic steatosis remain unknown. Notably, transcriptomic analyses of publicly available datasets revealed a modest but significant upregulation of *SLC25A35* in the human liver with MASLD compared with healthy subjects (**Figure 6A**). Similarly, higher *Slc25a35* mRNA levels were found in the liver of mice on a high-fat diet (HFD) than those on a normal-chow diet (NCD) (**Figure 6B**). Consistent with the results in adipocytes, we found that SLC25A35 loss in hepatocytes leads to the accumulation of PEP in the mitochondria, suggesting that SLC25A35 also plays a role in mitochondrial PEP export and glycerolipid synthesis in the liver (**Figure S9A**).

**Figure 6.**
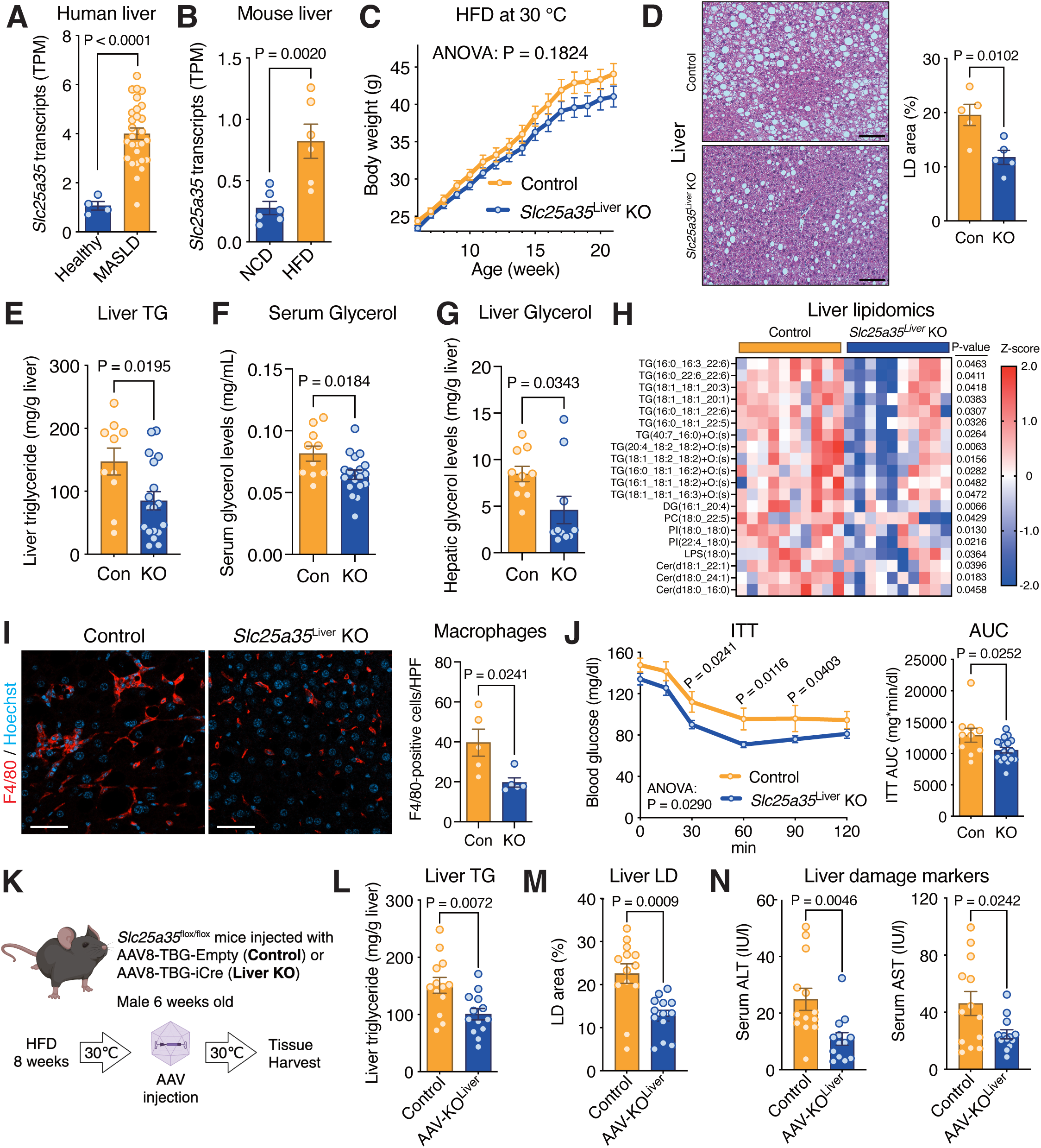
Blockade of mitochondrial PEP transport ameliorates hepatic steatosis in vivo. (A) TPM value for *SLC25A35* in human liver (GSE246221). n = 4 (healthy), n = 28 (MASLD). (B) TPM value for *Slc25a35* in the liver of mice fed a normal chow diet (NCD) or high-fat diet (HFD) (GSE199105). n = 6. (C) Body weight changes of male control mice and liver-specific *Slc25a35* KO mice on an HFD at 30°C. n = 10 (control), 18 (KO). (D) Representative H&E-staining of the liver of male mice in (C) at 19 weeks of HFD. Scale bars, 100 μm. Right: Quantification of LD area (%). n = 5. (E) Liver TG levels in male mice on an HFD, normalized to tissue weight (g). n = 10 (control), 18 (KO). (F) Serum glycerol levels in male mice on an HFD. n = 10 (control), 18 (KO). (G) Liver glycerol levels in male mice on an HFD, normalized to tissue weight (g). n = 10. (H) Liver lipidomics in male mice on an HFD. The top 20 lipid species reduced in KO mice are shown. n = 10. (I) Representative F4/80-stained liver sections of male mice on an HFD. Scale bars, 50 μm. Right: Quantification of F4/80-positive cells. n = 5. (J) Insulin tolerance test of male mice on an HFD for 18 weeks. Right: AUC. n = 10 (control), 18 (KO). (K) Schematic of inducible SLC25A35 depletion in the liver. (L) Liver TG contents in male mice at 10 weeks after AAV8 injection, normalized to tissue weight (g). n = 13. (M) Quantification of LD area (%) of the liver of male mice in (L). Representative H&E-staining is shown in Figure S10D. n = 13. (N) Serum ALT and AST levels in male mice in (L). n = 13.

To examine the extent to which liver-specific blockade of SLC25A35 prevents the pathogenesis of diet-induced hepatic steatosis, we developed mice that lacked *Slc25a35* in a liver-specific fashion by crossing *Slc25a35*^flox/flox^ mice with *Alb*-Cre (Alb-Cre x *Slc25a35*^flox/flox^, herein *Slc25a35*^Liver^ KO mice) (**Figure S9B**). When mice were housed at 30°C on a high-fat diet, *Slc25a35*^Liver^ KO mice showed a modest trend of lower body weight than littermate control mice (*Slc25a35*^flox/flox^ mice), although there was no significant difference between the genotypes (**Figure 6C**). At the tissue level, the liver weight of *Slc25a35*^Liver^ KO mice was significantly lower than that of control mice, while no difference was observed in other organs (**Figure S9C**). Hematoxylin and eosin (H&E) staining revealed that the liver of *Slc25a35*^Liver^ KO mice displayed reduced lipid droplet area compared to that of control mice (**Figure 6D**). Quantitative assays revealed significantly lower levels of triglycerides in the livers of *Slc25a35*^Liver^ KO mice compared to control mice (**Figure 6E**). Furthermore, glycerol levels in the serum and liver of SLC25A35Liver KO mice were significantly lower than those of control mice (**Figure 6F, G**). Importantly, lipidomics analyses of the liver identified several lipid species that were lower in *Slc25a35*^Liver^ KO mice than in control, including triglycerides (TG), diacylglycerol (DG) (16:1_20:4), phosphatidylcholine (PC) (18:0_22:5), phosphatidylinositol (PI), and ceramides (**Figure 6H, Table S4**). These changes in hepatic lipid content were accompanied by a reduced inflammatory profile in the liver. Histological analysis by immunostaining for macrophage marker (F4/80) found significantly less macrophage infiltration in the liver of liver-specific SLC25A35 KO mice than that in control mice (**Figure 6I**). Consistently, the liver of *Slc25a35*^Liver^ KO mice expressed significantly lower levels of genes involved in inflammation than that of control mice (**Figure S9D**).

The reduction of hepatic triglycerides and inflammation is associated with improved systemic glucose homeostasis ^59–63^. Thus, we next determined the extent to which liver-specific deletion of SLC25A35 protects mice from diet-induced insulin resistance. At 18 weeks of high-fat diet feeding, *Slc25a35*^Liver^ KO mice were more insulin sensitive than littermate controls, as shown by the insulin tolerance test (ITT) (**Figure 6J**). Similarly, pyruvate tolerance tests (PTT) at 17 weeks of high-fat diet showed that hepatic gluconeogenesis was significantly lower in *Slc25a35*^Liver^ KO mice than littermate control mice (**Figure S9E**). We then examined whether SLC25A35 is required for gluconeogenesis. To this end, we used ^13^C pyruvate as a tracer to measure ^13^C glucose production in the conditioned media of cultured primary hepatocytes. Glucagon increased the release of ^13^C-labeled glucose, validating this assay system for measuring gluconeogenesis. We found that *Slc25a35* KO hepatocytes generated equivalent levels of ^13^C-labeled glucose to control hepatocytes (**Figure S9F**). The data suggest that mitochondrial PEP via SLC25A35 plays an essential role in glycerolipid synthesis, whereas it makes a minor contribution to gluconeogenesis in the liver. The data suggest that the improved pyruvate tolerance in *Slc25a35*^Liver^ KO mice is due to reduced TG and hepatic inflammation.

Lastly, we explored the therapeutic potential of blocking SLC25A35-mediated lipogenesis by assessing the extent to which inducible SLC25A35 depletion in obese mice ameliorates hepatic steatosis. To this end, we acutely depleted SLC25A35 by delivering AAV8-TBG-iCre or AAV8-TBG-null (control) into the liver of *Slc25a35*^flox/flox^ mice fed on a high-fat diet for 8 weeks at 30°C (**Figure 6K**). At 10 weeks after AAV administration via the tail vein, delivery of AAV-Cre effectively reduced *Slc25a35* mRNA expression by over 90% (**Figure S10A**). Inducible depletion of SLC25A35 did not affect body weight (**Figure S10B**). Nonetheless, we found that inducible SLC25A35 depletion reduced hepatic triglyceride content by 33.4% (**Figure 6L**). Consistent with the Alb-Cre cohort, liver lipidomics showed that other lipid species, including diacylglycerides, phosphatidylglycerol (PG), and phosphatidic acid (PA) (16:0_18:1), were also decreased in *Slc25a35* AAV-KO^Liver^ mice (**Figure S10C, Table S5**). Histological analyses found that the liver of inducible *Slc25a35* KO mice displayed a reduced lipid droplet area than the control liver (**Figure 6M, Figure S10D**). Lastly, serum levels of ALT (Alanine Aminotransferase) and AST (Aspartate Aminotransferase), which are liver damage markers, were significantly reduced in *Slc25a35* AAV-KO^Liver^ mice compared with control mice (**Figure 6N**). We also found no difference in the hepatic expression of mitochondrial stress markers, including *Fgf21*, *Gdf15*, *Atf5*, and *Trib3,* between *Slc25a35* AAV-KO^Liver^ mice and control mice, although *Atf4* expression was modestly increased in AAV-KO^Liver^ mice (**Figure S10E**). Together, SLC25A35 blockade in the liver by two complementary approaches, Alb-Cre–mediated deletion and AAV-Cre–induced depletion, effectively ameliorated obesity-associated hepatic steatosis.

## DISCUSSION

Glyceroneogenesis is essential for generating G3P, the backbone required to esterify fatty acids into neutral triglycerides. By promoting fatty acid esterification and triglyceride storage, this pathway protects cells from lipotoxicity. When G3P accumulates in excess, Glycerol-3-Phosphate Phosphatase (G3PP) converts it to glycerol, thereby buffering the intracellular G3P pool, a process referred to as the glycerol shunt ^5,6,64^. In turn, chronic overactivation of glyceroneogenesis drives hepatic lipid overload, inflammation, and fibrosis, ultimately contributing to MASLD ^54,55^. Excess triglyceride accumulation in the liver is also a leading driver of insulin resistance and Type 2 diabetes ^65,66^. The present study identifies SLC25A35 as a critical gatekeeper of PEP transport across the IMM, providing a key substrate for the synthesis of G3P and glycerolipids, including triglycerides. In lipogenic states, as elicited by PPARγ agonists, high-fat diet feeding, or thermoneutrality, SLC25A35 expression parallels that of key lipogenic enzymes, such as acetyl-CoA carboxylase and fatty acid synthase. We found that blocking SLC25A35 effectively reduced the synthesis of pyruvate-derived G3P and glycerolipids in adipose tissue and the liver. Significantly, inducible depletion of SLC25A35 effectively alleviated hepatic steatosis without inducing liver damage. Together, this study identifies SLC25A35 as a target for selectively limiting excess fatty acid esterification and triglyceride synthesis, which are hallmarks of hepatic steatosis and insulin resistance.

Proteo-liposome-based reconstitution experiments found that PEP transport via SLC25A35 is pH gradient-dependent. The results suggest that mitochondrial PEP export into the cytosolic compartment is facilitated when mitochondrial membrane potential is high, positioning SLC25A35-mediated PEP export as a metabolic checkpoint linking energetic status to lipid synthesis. In this regard, thermoneutrality or obesogenic conditions would facilitate the export of mitochondria-derived PEP and glycerolipid synthesis. In agreement, thermoneutrality housing and a high-fat diet promote lipogenesis in the liver and MASLD, which is also accompanied by upregulation of lipogenic genes ^67,68^. Conversely, reduction of the mitochondrial membrane potential by liver-selective mitochondrial uncoupler ^69,70^ could be effective in limiting mitochondrial PEP export and glycerolipid synthesis.

Another important insight is the biological role of mitochondria-derived PEP via PCK2, even though PCK1 can catalyze the same reaction in the cytosol. We found that PCK1 overexpression was insufficient to restore TG synthesis in SLC25A35 KO adipocytes, indicating that the origin of PEP, *i.e.,* mitochondria-derived, is the crucial element, rather than its overall levels within a cell. It has been proposed that PCK2 provides an efficient pathway to generate PEP because the PCK2-mediated pathway requires fewer enzymatic reactions and metabolite transport ^25,28^. Our study proposes an alternative possibility that the mitochondria-derived PEP plays a key role in a specialized subset of mitochondria - those in close contact with lipid droplets, *a.k.a.,* peridroplet mitochondria ^71–73^, in lipogenic cells, including adipocytes and hepatocytes. Notably, proteomic analyses of BAT mitochondria-associated lipid droplet proteins detected SLC25A35 at high levels ^74,75^. In the liver, peridroplet mitochondria were observed at an early stage of hepatic steatosis ^76^. Thus, peridroplet mitochondria may be suited for providing not only ATP and citrate for fatty acid synthesis via SLC25A1 (the citrate carrier) but also PEP for G3P generation via SLC25A35, thereby supporting TG synthesis in adjacent lipid droplets. Targeting these “lipogenic mitochondria” can be effective in selectively suppressing fatty acid esterification in individuals with MASLD while preserving TCA cycle oxidation.

### Limitations of the study

The present study is limited to adipocytes and the liver. It is conceivable that mitochondria-derived PEP via SLC25A35 underpins additional physiological processes. For example, the glucose-induced PEP cycle in pancreatic beta cells, wherein mitochondria-derived PEP is consumed by pyruvate kinase (PK) to generate ATP in the cytosol, leads to the closure of the K_ATP_ channel and enhanced insulin secretion ^7–13^. Additionally, several cancer cells, such as non-small cell lung carcinoma and breast cancer, express high levels of PCK2, which controls mitochondrial PEP synthesis and tumor cell growth under low-glucose conditions ^77–81^. Future research is needed to investigate how mitochondrial PEP dynamics influence metabolic diseases, β-cell function, and cancer metabolism.

## RESOURCE AVAILABILITY

### Lead Contact

Further information and requests for resources and reagents should be directed to and will be fulfilled by the lead contact, Shingo Kajimura.

### Materials availability

All unique materials used, such as mouse strains and plasmids, are available upon request from the lead contact. Other materials are available from commercial sources as described in the text.

### Data and code availability

- The metabolomic LC-MS raw data are uploaded to the Metabolomics Workbench (https://www.metabolomicsworkbench.org), with a project ID PR002479 and a project DOI http://dx.doi.org/10.21228/M86J9P.
- The original code has been deposited at GitHub and is publicly available at: https://github.com/guimsilvaa/desmotools
- Source data are provided with this paper.
- Any additional information required to reanalyze the data reported in this paper is available from the lead contact upon request.

## ACKNOWLEDGMENTS

We are grateful to Anthony R.P. Verkerke, Ichitaro Abe, Christopher Auger, Jake Wong, Risa Waki, Hiroko Oda, and Kazusa Miyachi for their technical support. We also thank Drs. Bo Yuan and Tony Hui of the Harvard Chan Advanced Multi-Omics Platform for lipidomics, John Asara and the BIDMC mass spectrometry core facility for metabolomics, Evan Rosen at Beth Israel Deaconess Medical Center for sharing Adiponectin-Cre mice, Alexander Banks at Beth Israel Deaconess Medical Center for his support in metabolic cage studies, Suzanne L White at Beth Israel Deaconess Medical Center for her support in tissue histology study. We are also grateful for the discussion with Dr. Marc Prentki at Université de Montréal. This work was supported by grants from the National Institutes of Health (NIH) (DK125283, DK097441, and DK126160) and Howard Hughes Medical Institute to S.K. T.Y. was supported by the Nakatomi Foundation, the Yamada Science Foundation, the Takeda Science Foundation, and the American Heart Association postdoctoral fellowship (24POST1193689). D.K. was supported by the Manpei Suzuki Diabetes Foundation and the Japan Society for the Promotion of Science (JSPS). S.O. was supported by JSPS and the Uehara Memorial Foundation. Y.H. was supported by JSPS and the Sunstar Foundation. H.N. was supported by JSPS and the Uehara Memorial Foundation. M.F. was supported by the Manpei Suzuki Diabetes Foundation and the Kowa Life Science Foundation. N.Y. was supported by JSPS. J.S. was supported by the Manpei Suzuki Diabetes Foundation.

## AUTHOR CONTRIBUTIONS

Conceptualization, T.Y., S.K.; methodology, T.Y., D.K., G.M.S., S.O., Y.H., D.W., and N.Y.; investigation, T.Y., D.K., G.M.S., S.O., Y.H., D.W., H.N., N.Y., M. L., J.S., Z.Z., and J.-S. Y.; validation, T.Y., D.K., G.M.S., and Y.H.; formal analysis, T.Y., G.M.S., and M.F.; writing – original draft, T.Y., G.M.S., and S.K.; writing – review and editing, all the authors; funding acquisition, S.K.; supervision, L.S. and S.K.

## STAR METHODS

### EXPERIMENTAL MODEL AND STUDY PARTICIPANT DETAILS

#### Animals

All of the animal experiments in this study were performed in compliance with protocols approved by the Institutional Animal Care and Use Committee (IACUC, protocol# 028-2022-25) at Beth Israel Deaconess Medical Center. Unless otherwise specified, all of the mice had free access to food and water, and were housed under 12 h–12 h light–dark cycle, at 22°C, and 45% humidity on average. Heterozygous *Slc25a35*-floxed mice in the C57BL/6J background were generated by the *Easi*-CRISPR technology ^37^. Adipocyte-specific *Slc25a35* KO mice (*Slc25a35*^Adipo^ KO) and liver-specific *Slc25a35* KO mice (*Slc25a35*^Liver^ KO) were developed by crossing *Slc25a35*-floxed mice with Adiponectin-Cre mice (B6; FVB-Tg (Adipoq-Cre)1Evdr/J, 028020) and Albumin-Cre mice (B6.Cg-*Speer6-ps1^Tg(Alb-cre)21Mgn^*/J, 003574), respectively. *Slc25a35*^Adipo^ KO, *Slc25a35*^Liver^ KO, and their littermate control mice were maintained on an HFD (60% fat, D12492, Research Diets) starting at 6 weeks of age at 30°C. PTT and ITT were performed after 17 and 18 weeks of HFD feeding, respectively, in *Slc25a35*^Liver^ KO and their littermate control mice. Mice were euthanized for tissue collection after 19–23 weeks on an HFD. *Slc25a35* AAV-KO^Liver^ and control mice were developed by delivering AAV8-TBG-iCre or AAV8-TBG-null, respectively, into the liver of male *Slc25a35*^flox/flox^ mice fed an HFD for 8 weeks at 30°C. Mice were euthanized for tissue collection 10 weeks after AAV8 injection. Mouse sex is indicated in the figure legends. A list of the primer sequences used for genotyping and gRNA is provided in **Table S6**.

#### Cells

Stromal vascular fraction (SVF)-derived preadipocytes were isolated from the inguinal WAT of 7-week-old male and female *Slc25a35*-floxed mice. Cells were immortalized by expressing the SV40 large T antigen as described previously ^82^. For the generation of control and *Slc25a35*-KO cells, immortalized *Slc25a35*^flox/flox^ preadipocytes were infected with empty and Cre-expressing (34565, Addgene) retrovirus, followed by hygromycin (10687010, Thermo Fisher Scientific) selection at a dose of 200 μg ml^−1^, respectively. For the generation of *Slc25a35*-Flag–expressing cells, immortalized wild-type preadipocytes were infected with *Slc25a35*-Flag–expressing retrovirus, followed by blasticidin selection at a dose of 10 µg ml^−1^. For the generation of control, *Pck1*- and *Pck2*-depleted cells, immortalized preadipocytes were infected with an eSpCas9-LentiCRISPR v2 lentivirus with scrambled control, mouse *Pck1*, and mouse *Pck2* sgRNAs, followed by blasticidin selection at a dose of 10 µg ml^−1^, respectively. Unless otherwise specified, all of the cells were cultured in high-glucose DMEM containing 10% FBS, 1% penicillin–streptomycin, and 1% GlutaMAX™ at 37°C with 5% CO_2_. White adipocyte differentiation was induced in high-glucose DMEM containing 10% FBS, 1% penicillin–streptomycin, 1% GlutaMAX™, 5 µg ml^−1^ insulin, 1 nM T3, 1 µM rosiglitazone, 0.5 mM isobutylmethylxanthine, 125 nM indomethacin, and 2 µg ml^−1^ dexamethasone. After 48 h, cells were cultured in high-glucose DMEM containing 10% FBS, 1% penicillin–streptomycin, 1% GlutaMAX™, 5 µg ml^−1^ insulin, 1 nM T3, and 1 μM rosiglitazone for another four days ^83^. Brown adipocyte differentiation was induced in high-glucose DMEM containing 10% FBS, 1% penicillin–streptomycin, 1% GlutaMAX™, 0.5 µg ml^−1^ insulin, 1 nM T3, 0.5 mM isobutylmethylxanthine, 125 nM indomethacin, and 2 µg ml^−1^ dexamethasone. After 48 h, cells were cultured in high-glucose DMEM containing 10% FBS, 1% penicillin–streptomycin, 1% GlutaMAX™, 0.5 µg ml^−1^ insulin, and 1 nM T3 for another two days.

### METHOD DETAILS

#### DNA constructs, Antibodies, and Viruses

Mouse *Slc25a35* cDNA constructs were amplified from mouse inguinal WAT and inserted into a blasticidin-resistant pMSCV vector (75085, Addgene). Mouse *Pck1* cDNA constructs were amplified from mouse liver and inserted into a blasticidin-resistant pMSCV vector. The sgRNA sequences for scrambled control, mouse *Pck1* #1, mouse *Pck1* #2, mouse *Pck2* #1, and mouse *Pck2* #2 were inserted into a blasticidin-resistant eSpCas9-LentiCRISPR v2, which was generated by replacing the puromycin resistance cassette of the original eSpCas9-LentiCRISPR v2 (GenScript) with a blasticidin resistance cassette. An empty, hygromycin-resistant pMSCV vector was generated by removing the Cre insert from the Cre-expressing pMSCV (Addgene #34565). A list of the sgRNA sequences used in the study is provided in **Table S6**. All of the constructs were confirmed by sequencing.

The following antibodies were used in this study: anti-SLC25A35 (NBP2-85742, Novus Biologicals), anti-PEPCK1 (10004943, Cayman), anti-PEPCK2 (GTX114919, Genetex), anti-TOMM20 (11802-1-AP, Proteintech), anti-ATP5A (ab14748, Abcam), anti-Citrate Synthase (sc-390693, Santa Cruz), anti-Calreticulin (12238S, Cell Signaling Technology), anti-PEX14 (ABC142, Sigma-Aldrich), anti-EEA1 (2411S, Cell Signaling Technology), anti-GM130 (610822, BD Biosciences), anti-GAPDH (GTX100118, Genetex), anti-Histone H3 (sc-517576, Santa Cruz), anti-PDH (2784S, Cell Signaling Technology), anti-phospho-PDH α1 (Ser293) (37115S, Cell Signaling Technology), anti-Pyruvate Carboxylase (16588-1-AP, Proteintech), anti-Perilipin 1 (9349S, Cell Signaling Technology), anti-F4/80 (70076S, Cell Signaling Technology), anti-Vinculin (66305-1-IG, Proteintech), anti-Flag (8146S, Cell Signaling Technology), anti-β-actin (A3854, Sigma-Aldrich), goat anti-rabbit light chain HRP-conjugated antibody (NBP2-75935, Novus Biologicals), Strep-Tactin HRP conjugate (21502001, IBA), goat anti-mouse light chain HRP-conjugated antibody (91196S, Cell Signaling Technology), Alexa Fluor 488 (ab150117, Abcam), Alexa Fluor 555 (ab150062, Abcam), and Alexa Fluor 647 (A21245, Invitrogen). For retrovirus production, HEK293T packaging cells were transfected with 10 µg of retroviral transfer plasmids and 10 µg of packaging plasmids (gag-pol and pMD2.G) using Lipofectamine 3000. After 48 hours, the culture medium was collected and filtered using a 0.45 µm filter. Inguinal WAT-derived SVF cells were incubated in the four-times diluted viral medium supplemented with 10 µg ml^−1^ polybrene for 24 hours. For lentivirus production, HEK293T packaging cells were transfected with 10 µg of lentiviral transfer plasmids and 10 µg of packaging plasmids (psPAX2 and pMD2.G) using Lipofectamine 3000. After 48 hours, the culture medium was collected and filtered using a 0.45 µm filter. Inguinal WAT-derived SVF cells were incubated in the four-times diluted viral medium supplemented with 10 µg ml^−1^ polybrene for 24 hours.

#### Mitochondrial Isolation and Metabolomics

Following the established protocol ^84,85^, mitochondria were immunoprecipitated from SVF preadipocytes. In 150 mm cell culture dishes of SVF preadipocytes that express a MITO-Tag construct ^38^ (3xHA-EGFP-OMP25, 83356, Addgene), cells were washed twice with PBS. Cells were then scraped with KPBS (136 mM KCl, 10 mM KH2PO4, pH 7.25). Cells were pelleted by 1000 x g for 2 min at 4°C, and were resuspended in 1 mL of KPBS. One-tenth of cells was extracted directly into 80% MeOH as a whole-cell fraction. The remaining cells were homogenized with Teflon pestle and glass tube, and were centrifuged at 1000 x g for 2 min at 4°C. The supernatant was added to magnetic anti-HA beads (Pierce, 88836), and was rotated for 4 min at 4°C. The beads were collected for 1 min on a magnetic rack. The supernatant was aspirated and beads were washed three times with KPBS. After beads were washed, 80% MeOH was added to extract metabolites and the samples were stored at −80°C overnight. The samples were vortexed for 1 min and then centrifuged at 21000 x g for 10 min at 4°C. The supernatant was then transferred to a clean tube and was dried via a speed vac system. Dried metabolites were stored at −80°C for up to 1 week until resuspension in LC-MS grade H_2_O for the following LC-MS analysis. The samples were analyzed on a 5500 QTRAP hybrid triple quad quadrupole mass spectrometer (AB/SCIEX) coupled to a Prominence UFLC HPLC system (Shimadzu) with selected reaction monitoring (SRM) with positive/negative polarity switch. Peak areas were integrated using a Multi-Quant 2.1 software. The intensities were normalized to protein levels measured by a BCA assay. To minimize technical variance, values from *Slc25a35* KO cells were divided by those from the corresponding control cells within each individual sample. The data were analyzed using MetaboAnalyst. Missing values were estimated using the K-nearest neighbor (KNN) algorithm. Non-informative variables were filtered based on the interquartile range, with a 10% cutoff.

For differentiated adipocytes, cells cultured in a 150-mm dish were washed twice with ice-cold PBS and scraped into KPBS buffer (136 mM KCl, 10 mM KH₂PO₄, pH 7.25). The cell suspension was centrifuged at 1,000 × g for 2 min at 4°C, and the pellet was resuspended in 1 mL of KPBS. One-tenth of the cells were lysed in 80% methanol for whole-cell metabolomics analysis, and an aliquot was used for protein quantification by a BCA assay. The remaining cells were homogenized using a Teflon pestle and glass tube, followed by centrifugation at 600 × g for 5 min at 4°C. The supernatant was then centrifuged at 7,000 × g for 10 min at 4°C to isolate mitochondria. The resulting mitochondrial pellets were lysed in 80% methanol for metabolomics analysis, and an aliquot was used for protein quantification by a BCA assay. Metabolomics data were acquired using a UHPLC system (Vanquish Horizon, Thermo Scientific) coupled to an orbitrap mass spectrometer (Exploris 240, Thermo Scientific) as described previously ^85^. Waters ACQUITY UPLC BEH Amide column (particle size, 1.7 μm; 100mm (length) × 2.1mm (i.d.)) was used for LC separation. The column temperature was kept at 25°C. Mobile phases A = 25mM ammonium acetate and 25mM ammonium hydroxide in 100% water, and B = 100% acetonitrile, were used for negative mode. The linear gradient eluted from 95% B (0.0–1 min), 95% B to 65% B (1–7.0 min), 65% B to 40% B (7.0–8.0 min), 40% B (8.0–9.0 min), 40% B to 95% B (9.0–9.1 min), then stayed at 95% B for 5.9 min. The flow rate was 0.4 mL/min. The sample injection volume was 2 μL for cell and 5 μL for media. ESI source parameters were set as follows: spray voltage, 3500 V or −2800 V, in positive or negative modes, respectively; vaporizer temperature, 350°C; sheath gas, 50 arb; aux gas, 10 arb; ion transfer tube temperature, 325°C. The full scan was set as: orbitrap resolution, 60,000; maximum injection time, 100 ms; scan range, 70–1050 Da. The ddMS2 scan was set as: orbitrap resolution, 30,000; maximum injection time, 60 ms; top N setting, 6; isolation width, 1.0 m/z; HCD collision energy (%), 30; Dynamic exclusion mode was set as auto. Metabolite peak areas were quantified using TraceFinder software.

#### Metabolite Tracing

For whole-cell tracing, cells were incubated in high-glucose DMEM with 10 mM fully ^13^C-labeled pyruvate (Cambridge Isotope Laboratories, CLM-2440) at 37°C for one hour. Cells were washed in ice-cold PBS twice and were lysed in 80% MeOH. After one-hour incubation at −80°C, the supernatant was collected for metabolomics. For mitochondria tracing, cells in a 150 mm cell culture dish were washed in ice-cold PBS twice, and were scraped with KPBS (136 mM KCl, 10 mM KH_2_PO_4_, pH 7.25). Cells were pelleted by 1000 x g for 2 min at 4°C, and were resuspended in 1 mL of KPBS. Cells were homogenized with Teflon pestle and glass tube, and were centrifuged by 600 x g for 5 min at 4°C. The supernatant was centrifuged by 7,000 x g for 10 min at 4°C. The resultant pellets were incubated in KPBS with fully ^13^C-labeled pyruvate for the indicated time at 4°C and were washed two times with KPBS. The pellets were lysed in 80% MeOH and were stored at −80°C for the following metabolomics.

Metabolomics data were acquired using the same UHPLC settings as those used for differentiated adipocytes. The labeling metabolomics was quantified by a FreeStyle 1.8 SP2 or Compound Discoverer 3.3. The annotation of metabolites was performed by searching the retention time and MS2 against our in-house library which was generated using the Mass Spectrometry Metabolite Library (MSMLS™) (IROA Technologies LLC®, Bolton, MA, USA). The labeling rate was calculated by dividing the level of labeled metabolites by the total metabolite level. An additive log-ratio (ALR) transformation using M+0 as the denominator was applied to account for the compositional nature of isotopologue fractions (*i.e.*, [M+n] values are interdependent and constrained to sum to 100%). To avoid undefined values arising from zeros, a small pseudocount (0.1) was added to both M+n and M+0 prior to the ALR transformation. This log-ratio transformation maps the data from the simplex to an unconstrained scale, enabling the use of standard parametric analyses such as two-way ANOVA.

#### Mitochondrial PEP Export Assay

Cells in a 150 mm cell culture dish were washed in ice-cold PBS twice, and were scraped with KPBS (136 mM KCl, 10 mM KH_2_PO_4_, pH 7.25). Cells were pelleted by 1000 x g for 2 min at 4°C, and were resuspended in 1 mL of KPBS. Cells were homogenized with Teflon pestle and glass tube, and were centrifuged by 600 x g for 5 min at 4°C. The supernatant was centrifuged by 7,000 x g for 10 min at 4°C. The resultant pellets were incubated in KPBS with 10 mM fully ^13^C-labeled pyruvate, 10 mM glutamate, 2 mM malate, and 10 mM succinate for the indicated time at 4°C and were centrifuged by 21,000 x g for 10 min at 4°C. The supernatant was added with MeOH to make the final concentration 80% for the following LC-MS analysis to detect ^13^C_3_-labeled PEP. The protein concentration in the pellets was determined by a BCA assay for normalization.

#### Protein expression and purification

Sf9 cells (11496015, Thermo Fisher Scientific) were cultured in Sf-900 II SFM (10902088, Thermo Fisher Scientific) with 10% FBS, 1% penicillin–streptomycin, and 1% GlutaMAX™ at 27°C without CO_2_. Baculovirus packaging and amplification were performed based on the commercial protocol for a Bac-to-Bac C-His TOPO Expression System (A11100, Thermo Fisher Scientific). In brief, codon-optimized c-terminally Twin-Strep-tagged mouse *Slc25a35* cDNA was synthesized in Integrated DNA Technologies and was inserted into a pFastBac TOPO vector. The vector was transformed into DH10Bac *Escherichia coli* competent cells to form a bacmid. The bacmid was transfected into Sf9 cells using an ExpiFectamine Sf Transfection Reagent for the production of recombinant baculovirus particles (P0 virus). P0 virus was infected into Sf9 cells to generate P1 virus, which was further infected into Sf9 cells to generate high-titer P2 virus. For protein purification, Sf9 cells in 200 ml of medium were infected with P2 virus and 72-120 hours later collected by a centrifugation. Cells were homogenized with Teflon pestle and glass tube, and were centrifuged by 600 x g for 5 min at 4°C. The supernatant was centrifuged by 10,000 x g for 10 min at 4°C. The resultant pellets were stored at −80 °C. The pellets were lysed in W buffer (IBA, 2-1003-100) with 2% Triton X-100 and a protease inhibitor cocktail (Roche) and were rotated at 4°C for 2 hours. The lysate was centrifuged at 21,000 rpm for 5 min, and the supernatant was rotated with a Strep-TactinXT 4Flow high-capacity resin (IBA, 2-5030-010) at 4°C for 3 hours. The resin was washed five times with W buffer containing 2% Triton X-100 and a protease inhibitor cocktail, followed by four rounds of rotation in BXT buffer (IBA, 2-1042-025) supplemented with 2% Triton X-100 and a protease inhibitor cocktail at 4°C for a total of 2 hours to purify the protein. For SLC25A35 mutant analysis, the codons TAT, AGA, and CGT were mutated to GCC to generate the Y124A, R175A, and R276A mutants, respectively, using an In-Fusion Cloning kit (Takara Bio).

#### Proteoliposomes

To prepare liposomes, 100 mg of the lipids (Egg PC, E coli Polar lipids, 18:1 Cardiolipin at 4:4.2:9 ratio) in 10 ml of chloroform was incubated in a rotary evaporator at 100 rpm at 50°C overnight. The lipid film on the internal surface of the flask was rehydrated with 2 ml of PIPES buffer (10 mM PIPES, 50 mM NaCl, pH 7.0) containing 20 mM unlabeled PEP. The liposomes were extruded 15 times using a mini-extruder with a 200-nm pore membrane at 60°C. Extruded liposomes were rotated with a purified SLC25A35 protein or background proteins (purified from empty virus-infected cells) at 4°C for 1 hour. Proteo-liposomes were rotated with Bio-Beads SM-2 five times to completely remove triton X-100. The resultant proteo-liposomes were isolated on a PD-10 desalting column to remove the external substrates and residual Bio-Beads, and were centrifuged at 100,000 rpm for 10 min. Proteo-liposomes were incubated with 5 mM of ^13^C_2_-labeled PEP (Cambridge Isotope Laboratories, CLM-3398) in PIPES buffer at 37°C for 20 min, and centrifuged at 100,000 rpm for 5 min. The pellet was washed twice with PIPES buffer, each followed by centrifugation at 100,000 rpm for 5 min. The final pellet was lysed in 80% MeOH for the following LC/MS analysis to detect ^13^C_2_-labeled PEP. To mimic more physiological conditions, liposomes were loaded with 4 mM or 10 mM non-labeled PEP and then incubated with 1 mM or 2.5 mM ¹³C₂-labeled PEP to evaluate PEP transport. For proteinase K assay, the liposomes were treated with or without 50 μg/mL of proteinase K in PIPES buffer at 4°C for 10 min prior to the transport assay. For KIC/KIV transport, liposomes were loaded with 20 mM non-labeled KIC/KIV and incubated with 5 mM fully ¹³C-labeled KIC/KIV. To exclude non-specific signals, liposomes without reconstituted proteins were subjected to the same transport assay, and the average background signal was subtracted from the values obtained with control and SLC25A35 proteoliposomes, although the background signal itself was very low.

#### *In silico* Structural Studies

The AlphaFold 2.0 structure of human SLC25A35 was retrieved from AlphaFoldDB (https://alphafold.ebi.ac.uk/) under the code: AF-Q3KQZ1-F1-model_v4. All figures were generated using Pymol 3.0.2. The AlphaFold structure was submitted to the Protein Preparation Workflow on Maestro, by Schrodinger (Suite 2024-2), to re-assign bond orders using CCD (Chemical Component Dictionary) database, add/replace hydrogens, fill in missing side chains, generate disulfide bonds, optimize hydrogen bonds with PROPKA, and perform restrained minimization to 0.30 Å root-mean-square deviation (RMSD) using the OPLS4 force field.

Phosphoenolpyruvate (PEP) canonical SMILES string was retrieved from PubChem (https://pubchem.ncbi.nlm.nih.gov/) CID 1005. Ligprep was used in default settings to convert the SMILES string to 3D structure and minimize it to a most energetically favorable conformation, considering Epik pH = 7.0 ± 2.0, up to 32 tautomer/stereoisomers, and OPLS4 force field.

For docking studies, the Schrodinger’s module Receptor Grid Generation was used to specify the ligand binding site based on the volume of the prioritized Sitemap pocket 1 within the SLC25A35 central channel. This grid corresponds to 20 and 10 Å of outer and inner enclosed boxes, respectively, and the centroid coordinates of x = 5.09, y = 2.07, and z = −2.02. This binding site was used to run PEP docking by using default parameters and up to 10 poses generation with both GlideSP and GlideXP scoring functions. The poses generated were submitted to the Molecular Mechanics with Generalized Born and Surface Area (MMGBSA) solvation module from Schrodinger’s Prime to estimate binding free energy ΔG values in kcal/mol. The best pose (lowest ΔG) was then used as reference for conducting Induced-Fit Docking (IFD), by considering default settings and up to 20 poses generation per molecule. The resulting IFD poses were again rescored through Prime MMGBSA and clustered by visual inspection in groups with matching poses (orientation + conformation). The most populated group consisted of 12 matching poses. From this, the pose with the lowest ΔG free energy value, in kcal/mol, was selected as the most representative SLC25A35-PEP docking complex.

Molecular dynamics (MD) simulation studies: MD simulations were performed for the *apo* AlphaFold prepared structure of SLC25A35, as well as those in complex/docked to the ligand PEP. Desmond’s System Builder was used to prepare each system comprising the *apo* protein or the protein-ligand complex. These were solvated with water TIP3P solvent model and neutralized with addition of ions and buffer NaCl at 0.15 M. The OPM webserver (https://opm.phar.umich.edu/) was used to obtain a mitochondrial inner membrane orientation model (POPC at 300 K) for the SLC25A35 prepared structures. Also, we considered orthorhombic boxes with minimized volumes, keeping each side distanced by a minimum distance of 10 Å from protein atoms. The force field OPLS4 was used and all the other options were set to default values. MD runs were conducted using Desmond by producing trajectories of 300 ns (ca. 1000 frames), 1.2 ps for energy recording interval, isothermal-isobaric (NPT) ensemble at 300 K and 1.01325 bar, and allowing relaxation of system before simulation. Each run was repeated 5 times (5x replicates). Jobs were written and conducted on the O2 High Performance Compute Cluster at Harvard Medical School, through licensed software of the SBGrid Consortium. Produced trajectories were submitted to Schrodinger’s Simulation Interactions Diagram to build plots and related metrics for analysis, such as Root Mean Square Fluctuation (RMSF), RMSD, among others. Additionally, RMSD boxplots as well as merged RMSF graphs were generated using our in-house script at the https://github.com/guimsilvaa/desmotools repository.

The MMGBSA ΔG free energy values between protein-ligand complexes were calculated for each MD trajectory-frame by Schrodinger’s thermal_mmgbsa.py script. These were calculated using the OPLS4 force field and the default Prime tool. The protocol involves five fundamental entities for energy calculations: optimized free protein, optimized free ligand, optimized complex, protein from optimized complex, and ligand from optimized complex. Different components contribute to the calculation of total ΔG binding affinity energy in kcal/mol: Coulomb energy, covalent binding energy, van der Waals (vdW) energy, lipophilic energy, generalized Born electrostatic solvation energy, total Prime energy, hydrogen-bonding correction, π-π stacking correction, and self-contact correction. Statistical parameters such as average and corresponding standard deviation values for ΔG and associated components were also calculated using available scripts at our custom-built desmotools repository.

#### Glycerolipid Synthesis Assay

To assess the contribution of mitochondrial PEP in glycerolipid formation, we used ^14^C pyruvate (1-^14^C Sodium pyruvate, ARC) and quantified the incorporation of ^14^C pyruvate into the lipids. In cultured differentiated adipocytes, media were replaced in high-glucose DMEM containing 10% FBS, 1% penicillin–streptomycin, and 1% GlutaMAX™ on day 5. Cells on day 6 were incubated in high-glucose DMEM (Wako, 048-33575) that contained ^14^C pyruvate at 0.25 μCi ml^-1^ in the presence or absence of 5 μg ml^-1^ insulin at 37°C with 5% CO_2_ for 8 hours. Similarly, we used ^14^C glucose (U-^14^C_6_ D-GLUCOSE, ARC) at 0.5 μCi ml^-1^ as an alternative tracer.

Cells were washed with PBS twice and lysed in 200 μl of lysis buffer (25 mM Tris-HCl, pH 7.5, 1 mM EDTA, 1% Triton X-100). 150 μl of lysate was mixed with the same amount of Folch solution (2:1 (v/v) chloroform/methanol) and centrifuged at 21,000 x g for 5 min. The lower (lipid) phase was added with 5 ml of scintillation cocktail Ultima Gold to measure the ^14^C radioactivity (dpm) using a liquid scintillation counter. The remaining lysate was used to determine the protein concentration by BCA assays for normalization.

To quantify 1-^14^C pyruvate incorporation into the fatty acid fraction of glycerolipids, lipids were extracted as described above, evaporated, and mixed with 500 μl of 1 M KOH in methanol, and incubated at 60°C for 1 h. To neutralize KOH and protonate fatty acids, 50 μl of 37% HCl was added. Fatty acids were extracted three times with 800 μl of hexane:diethyl ether (1:1, v/v), with vortexing and centrifugation after each extraction, followed by an additional wash with 500 μl of hexane to ensure complete extraction of fatty acids. The fatty acid fraction was evaporated and resuspended in 20 μl of Folch solution for TLC separation. Samples (10 μl) were spotted 1 cm from the bottom of TLC plates and developed in 80:20:1 (v/v/v) hexane: diethyl ether: acetic acid until the solvent front reached 10 cm. A 10 mg/mL stearic acid solution in Folch solution was used as a standard. Based on the Rf value of fatty acids (∼0.30), as visualized by CuSO₄/H₃PO₄ staining, the corresponding regions were scraped into 1 ml of water in scintillation vials. Finally, 5 ml of scintillation cocktail (Ultima Gold) was added, and ¹⁴C radioactivity (dpm) was measured using a liquid scintillation counter.

To assess glycerolipid synthesis in adipose tissue, lipids were separated by TLC, and the glycerol backbone–and fatty acid–containing fractions were visualized by PAS staining and CuSO₄/H₃PO₄ staining, respectively ^53^. Approximately 50 mg of white adipose tissues that were incubated in high-glucose DMEM (Wako, 048-33575) containing 0.5 μCi ml^-1^ ^14^C glucose (U-^14^C_6_ D-GLUCOSE, ARC) with or without 5 μg ml^-1^ insulin at 37°C with 5% CO_2_ for 4 hours. Tissues were washed with PBS twice, homogenized in 1 ml of Folch solution (2:1 (v/v) chloroform:methanol), and incubated on an orbital shaker at room temperature overnight. The homogenates were mixed with 200 μL of 150 mM NaCl solution and were centrifuged at 21,000 x g for 5 min. The lower (lipid) phase was collected, evaporated, mixed with 500 μl of 1 M KOH in methanol, and incubated at 60°C for 1 h. To neutralize KOH and protonate fatty acids, 50 μl of 37% HCl was added. Fatty acids were extracted three times with 800 μl of hexane:diethyl ether (1:1, v/v), with vortexing and centrifugation after each extraction, followed by an additional wash with 500 μl of hexane to ensure complete extraction of fatty acids. The fatty acid fraction was evaporated and resuspended in 50 μl of Folch solution for TLC separation. The glycerol-containing aqueous phase was evaporated to approximately 100 μl, mixed with 1 ml of acetone, and vortexed to precipitate KCl. After centrifugation, glycerol-containing supernatants were evaporated to dryness and resuspended in 50 μl of methanol for TLC separation. Samples (10 μl) were spotted 1 cm from the bottom of TLC plates and developed in 80:19:1 (v/v/v) acetonitrile:water:acetic acid and 80:20:1 (v/v/v) hexane:diethyl ether:acetic acid until the solvent front reached 10 cm for glycerol fraction and fatty acid fraction, respectively. A 10% (v/v) glycerol solution in methanol and a 10 mg/mL stearic acid solution in Folch solution were used as standards. Based on the Rf values of glycerol (∼0.55) and fatty acids (∼0.30), as visualized by PAS staining and CuSO₄/H₃PO₄ staining, respectively, the corresponding regions were scraped into 1 ml of water in scintillation vials. For PAS staining, plates were sprayed with 0.5% periodate in 70% ethanol, dried, then sprayed with 50% Schiff reagent in ethanol to visualize glycerol spots. For CuSO₄/H₃PO₄ staining, plates were sprayed with 10% (w/v) CuSO₄·5H₂O and 8% (v/v) phosphoric acid (85% stock) in distilled water, dried, and then heated at 180°C to visualize fatty acid spots. Finally, 5 ml of scintillation cocktail (Ultima Gold) was added, and ¹⁴C radioactivity (dpm) was measured using a liquid scintillation counter. Radioactivity values were normalized to the initial tissue weight.

#### Lipidomics

Liver lipids were extracted in butanol/methanol (1:1) with 5 mM ammonium formate, and were analyzed using a Dionex Ultimate 3000 RSLC system (Thermo Scientific) coupled with a QExactive mass spectrometer (Thermo Scientific, Waltham, MA, USA). Chromatographic separation was achieved on an ACQUITY UPLC CSH C18 column (130Å, 1.7 μm, 2.1 mm × 100 mm) with an ACQUITY UPLC CSH C18 VanGuard pre-column (130Å, 1.7 μm, 2.1 mm × 5 mm) (Waters, Milford, MA) with column temperature at 50°C. For the gradient, mobile phase A consisted of an acetonitrile-water mixture (6:4), and mobile phase B was a 2-propanol-acetonitrile mixture (9:1), both phases containing 10 mM ammonium formate and 0.1% formic acid. The linear elution gradient was: 0-3 min, 20% B; 3-7 min, 20-55% B; 7-15 min, 55-65% B; 15-21 min, 65-70% B; 21-24 min, 70-100% B; and 24-26 min, 100% B, 26-28 min, 100-20% B, 28-30 min, 20% B, with a flow rate of 0.35 mL/ min. The autosampler was at 4°C. The injection volume was 5 μL. Needle wash was applied between samples using a mixture of dichloromethane-isopropanol-acetonitrile (1:1:1).

ESI-MS analysis was performed in positive and negative ionization polarities using a combined full mass scan and data-dependent MS/MS (Top 10) (Full MS/dd-MS2) approach. The experimental conditions for full scanning were as follows: resolving power, 70,000; automatic gain control (AGC) target, 1 × 106; and maximum injection time (IT), 100 ms. The scan range of the instrument was set to m/z 100-1200 in both positive and negative ion modes. The experimental conditions for the data-dependent product ion scanning were as follows: resolving power, 17,500; AGC target, 5 × 104; and maximum IT, 50 ms. The isolation width and stepped normalized collision energy (NCE) were set to 1.0 m/z, and 10, 20, and 40 eV. The intensity threshold of precursor ions for dd-MS2 analysis and the dynamic exclusion were set to 1.6 × 105 and 10 s. The ionization conditions in the positive mode were as follows: sheath gas flow rate, 50 arb; auxiliary (AUX) gas flow rate, 15 arb; sweep gas flow rate, 1 arb; ion spray voltage, 3.5 kV; AUX gas heater temperature, 325°C; capillary temperature, 350°C; and S-lens RF level, 55. The ionization conditions in the negative mode were as follows: sheath gas flow rate, 45 arb; auxiliary (AUX) gas flow rate, 10 arb; sweep gas flow rate, 1 arb; ion spray voltage, 2.5 kV; AUX gas heater temperature, 320°C; capillary temperature, 320°C; and S-lens RF level, 55.

Thermo Scientific LipidSearch software version 5.0 was used for lipid identification and quantitation. First, the product search mode was used during which lipids are identified based on the exact mass of the precursor ions and the mass spectra resulting from product ion scanning. The precursor and product tolerances were set to 10 and 10 ppm mass windows. The absolute intensity threshold of precursor ions and the relative intensity threshold of product ions were set to 30000 and 1%. Next, the search results from the individual positive or negative ion files from each sample were aligned within a retention time window (±0.25 min) and then all the data were merged for each annotated lipid with a retention time correction tolerance of 0.5 min. The annotated lipids were then filtered to reduce false positives by only including the lipids with a total grade of A or B. To improve data quality and interpretability, lipid species with non-physiological features were excluded from downstream analysis. These included lipids with an unusually low (14 or fewer) or high (26 or greater) number of total carbon atoms, and those with odd-chain fatty acid moieties, as these are rarely found in mammalian tissues and may arise from artifacts or misannotation. Nonetheless, all of the statistically significant lipids are listed in **Table S4 and S5** for *Slc25a35*^Liver^ KO and AAV-KO^Liver^ mice, respectively. The intensities were normalized to the sum of total lipid intensities. The data were analyzed using MetaboAnalyst. Non-informative variables were filtered based on the interquartile range, with a 20% cutoff.

#### Animal Physiology

Body weight was measured every week. For an insulin-tolerance test, mice were fasted for 6 hours and intraperitoneally injected with insulin (0.75 U kg^−1^ body weight). For a pyruvate-tolerance test, mice were fasted for 16 hours and intraperitoneally injected with pyruvate (1.5 g kg^−1^ body weight). Blood glucose levels were measured at the indicated time points using blood glucose test strips (Freestyle Lite). *Slc25a35*^Adipo^ KO mice and littermate control mice were monitored using a Promethion Metabolic Cage System (Sable Systems) to measure whole-body energy expenditure (VO_2_, VCO_2_), food intake, and locomotor activity (beam break counts). Mice were intraperitoneally injected with CL-316,243 (Sigma-Aldrich; 0.5 mg kg^-1^ body weight).

#### Histology

Adipose tissues and the liver were fixed in 4% paraformaldehyde at 4°C overnight, followed by incubation in 70% ethanol. After the dehydration procedure, tissues were embedded in paraffin and cut into sections at 5 μm thickness. The sections were stained with haematoxylin and eosin according to the standard protocol at the BIDMC pathology core. Images were acquired using a Zeiss AxioImager M1 microscope with a 20× objective at the BIDMC imaging core.

#### Primary hepatocyte Assays

Mouse liver was perfused with liberase (05401127001, Sigma-Aldrich), and the dissociated cells were washed with William’s E medium (12551032, Thermo Fisher Scientific). After purification by Percoll (17089101, Cytiva) density gradient centrifugation, the cells were cultured in William’s E medium supplemented with 10% FBS, 1% penicillin–streptomycin, and 1% GlutaMAX™ at 37°C. After a 6-hour attachment period, the cells were subjected to gluconeogenesis assays and mitochondrial metabolomics. For the gluconeogenesis assay, cells were incubated in glucose-free DMEM containing 10 mM fully ^13^C-labeled pyruvate with or without 100 nM glucagon at 37°C. The culture medium was collected at the indicated time points for measurement of ^13^C-labeled glucose by LC-MS. The cells were lysed, and protein content was quantified using a BCA assay for normalization. For mitochondrial metabolomics, cells cultured in a 150-mm dish were washed twice with ice-cold PBS and scraped into KPBS buffer (136 mM KCl, 10 mM KH₂PO₄, pH 7.25). The cell suspension was centrifuged at 1,000 × g for 2 min at 4°C, and the pellet was resuspended in 1 mL of KPBS. One-tenth of the cells were lysed in 80% methanol for whole-cell metabolomics analysis, and an aliquot was used for protein quantification by a BCA assay. The remaining cells were homogenized using a Teflon pestle and glass tube, followed by centrifugation at 600 × g for 5 min at 4°C. The supernatant was then centrifuged at 7,000 × g for 10 min at 4°C to isolate mitochondria. The resulting mitochondrial pellets were lysed in 80% methanol for metabolomics analysis, and an aliquot was used for protein quantification by a BCA assay. The mitochondrial metabolomics analysis was performed using the same method as that used for differentiated adipocytes.

#### Lipolysis and Glucose Oxidation Assays

Approximately 50 mg of adipose tissues were incubated in 600 μl of high-glucose DMEM containing 2% BSA with or without 10 μM isoproterenol (Sigma-Aldrich) at 37°C for 3 hours. The cultured medium was collected to measure the glycerol levels using a free glycerol reagent (Sigma-Aldrich). The glycerol content was normalized to the initial tissue weight. Glucose oxidation assays were performed according to the protocol described in our previous work ^84,86^. In brief, adipose tissues were minced into small pieces, placed into a polypropylene round-bottom tube, and incubated in 1 ml of KRB-HEPES buffer with 0.5 μCi ml^−1^ ^14^C glucose (U-^14^C_6_ D-GLUCOSE, ARC) at 37°C for 1 hour. The reaction mixture was added with 350 μl of 30% hydrogen peroxide, and ^14^CO_2_ was trapped in the center well with 300 μl of 1 M benzethonium hydroxide solution for 20 min at room temperature. The ^14^C radioactivity was measured using a liquid scintillation counter and normalized to the initial tissue weight.

#### Lipid profiling

For the measurement of liver triglyceride contents, approximately 50 mg of liver tissues were homogenized in 1 ml of Folch solution (2:1 (v/v) chloroform:methanol), and were incubated on an orbital shaker at room temperature overnight. The homogenates were mixed with 200 μL of 150 mM NaCl solution and were centrifuged at 21,000 x g for 5 min. The lower (lipid) phase was evaporated to measure the triglyceride amount using an Infinity Triglycerides kit (Thermo Fisher Scientific, TR22421). The liver triglyceride levels were normalized to the initial tissue weight. Serum AST and ALT levels were measured using a Stanbio AST testing (Stanbio, SBL-2930-430) and Stanbio ALT testing (Stanbio, SBL-2920-430), respectively. Serum glycerol levels were measured using a free glycerol reagent (F6428, Sigma-Aldrich) and glycerol standard solution (G7793, Sigma-Aldrich). For liver glycerol measurement, approximately 30-40 mg of liver tissues were homogenized in 1 ml of PBS. The glycerol levels in the homogenates were measured using a free glycerol reagent and glycerol standard solution. The liver glycerol levels were normalized to the initial tissue weight. Cellular glycerol-3P was determined using a Glycerol 3-Phosphate (G3P) Assay Kit (A3854, Sigma-Aldrich). The glycerol-3P levels were normalized to protein levels measured by a BCA assay. For cellular triglyceride measurement, cells were lysed in 200 μl of lysis buffer (25 mM Tris-HCl, pH 7.5, 1 mM EDTA, 1% Triton X-100). An aliquot of the lysate was mixed with the same amount of Folch solution (2:1 (v/v) chloroform/methanol). The lower phase was collected, and TG content determined, using an Infinity Triglycerides kit (Thermo Fisher Scientific, TR22421). Another aliquot of the lysate was used for protein normalization by a BCA assay.

#### Immunoblotting

Protein lysates were mixed with 5 x SDS sample buffer, were incubated at 37°C for 10 min, and were separated by SDS–PAGE to be transferred to a PVDF membrane. Membranes were blocked in 1% skim milk TBS-T and were incubated with a specific primary antibody. Immunoreactive bands were detected using an HRP-conjugated secondary antibody, were visualized with a Clarity Western ECL Substrate (Bio-Rad) or an ImmunoStar LD (Wako), and were imaged using a ChemiDoc Touch (Bio-Rad). β-actin was used as a loading control. The band intensity was normalized against the loading control for quantification. Band intensity was quantified using an ImageJ software (NIH). Molecular mass (kDa) is shown at the top right.

#### Immunostaining

Cells were placed in glass-bottom dishes (VWR 10810-054) and cultured for 24 hours. Differentiation was induced and maintained on the same glass-bottom dish until day 3. The cells were washed twice with PBS and were fixed with 4% PFA at 37°C for 30 min. The samples were rinsed three times with PBS and were permeabilized with 0.3% NP-40, 0.05% Triton X-100, and 0.1% BSA in PBS for 3 min. After three rinses with wash buffer (0.05% NP-40, 0.05% Triton X-100, and 0.2% BSA in PBS), the samples were blocked with SuperBlock Blocking Buffer (Thermo Fisher, 37515) at room temperature for 1 hour. The samples were incubated with a primary antibody in wash buffer at 4°C overnight, were washed three times with wash buffer. The samples were incubated with a secondary antibody at room temperature for 3 hours and washed three times with wash buffer. Right panels show line-scan fluorescence intensity profiles measured along the dashed white lines in the magnified images. Images were acquired with a Zeiss LSM900 confocal microscope. Images were processed using Zeiss Zen software.

For immunostaining in the liver, liver tissues from *Slc25a35*^liver^ KO and control mice were fixed in 4% paraformaldehyde (PFA) overnight and processed them for paraffin embedding. Heat-mediated antigen retrieval was performed using citrate buffer (pH 6.0). After blocking with 10% donkey serum (D9663, Sigma-Aldrich) for 30 minutes, the sections were incubated with a rabbit monoclonal anti-F4/80 antibody (70076S, Cell Signaling Technology; 1:200) overnight at 4°C. The sections were then washed with PBS and incubated with a secondary antibody (ab150062, Abcam; 1:500) for 1 hour, followed by nuclear staining with Hoechst (ab228551, Abcam; 1:2000). Fluorescent images were acquired using a Zeiss LSM900 confocal microscope and analyzed in four randomly selected fields per liver to quantify the number of F4/80⁺ Hoechst^+^ cells (macrophages) in the liver tissue.

#### Proteinase K assay

Cells were scraped and pelleted at 1100 x g for 2 min. Cell pellets were washed with phosphate buffer solution (PBS) and re-pelleted at 1100 x g for 2 min. Cell pellets were then homogenized in buffer (22 mM mannitol, 75 mM sucrose, 1 mM EGTA, 30 mM Tris-HCl, pH 7.4). A part of the homogenate was served as a whole cell fraction. The remaining homogenate was centrifuged at 600 x g for 5 min and the supernatant was pelleted at 7000 x g for 10 min. The supernatant was served as a cytosol fraction. The pellet was added with 50 μg/mL of proteinase K in buffer (150 mM KCl, 10 mM HEPES, 200 μM CaCl2, pH 7.2), incubated on ice for 10 min, and centrifuged at 7000 x g for 10 min. The resultant pellet was added with sample buffer for immunoblotting. Immunoblotting was performed to detect SLC25A35-FLAG, TOMM20 (mitochondrial outer membrane), ATP5A (mitochondrial inner membrane), and CS (mitochondrial matrix).

#### qRT-PCR

Total RNA was isolated from cells or tissues using a TRIzol reagent (Invitrogen) and a Zymo Direct-zol RNA preparation kit (R2052, Zymo). The RNA samples were reverse-transcribed into cDNAs using an iScript cDNA Synthesis Kit (Bio-Rad Laboratories). The cDNAs were quantified using a QuantStudio 6 system (Applied Biosystems). *36b4* was served as an internal control. A list of the qRT-PCR primers used in this study is provided in **Table S6.**

#### Single-nuclei RNA-seq

Data from GSE176171 ^33^ were analyzed and visualized using R and Seurat (v4.0.1), following standard workflows. Dot size indicates the percentage of cells in each cluster with detectable transcripts, and color represents average scaled transcript abundance.

### QUANTIFICATION AND STATISTICAL ANALYSIS

Statistical analyses were performed using a GraphPad Prism v.10 (GraphPad). All data are represented as mean ± s.e.m. unless otherwise specified. Data were obtained from biologically independent samples. Unpaired Student’s *t*-tests were performed for two-group comparisons (Figure 2A, 3D, 5B, 5I, 5J, 6D-G, 6I, 6J (AUC), 6L-N, Figure S7D-F, S7H-J, S8G, S9A-D, S9E (AUC), S10A, S10E). One-way ANOVA followed by appropriate post-hoc tests was performed for multiple-group comparisons (Tukey’s multiple comparisons test: Figure 5F, Figure S2D, S7A, S8H, S8I; Holm-Šídák’s multiple comparisons test: Figure 1E, 4G, 5C-E, 5K, 5L, Figure S2B, S2E, S6F, S6G, S7B, S7C, S8E, S8F). Two-way ANOVA followed by appropriate post-hoc tests was performed to analyze the effects of two independent factors (uncorrected Fisher’s LSD test: Figure 2I, 3C, 3H, Figure S4A; Bonferroni’s multiple comparisons test: Figure 3F, 3G; Tukey’s multiple comparisons test: Figure 3J; Šídák’s multiple comparisons test: Figure S3E). Two-way repeated-measures ANOVA followed by uncorrected Fisher’s LSD test was performed to determine the statistical difference in body weight, whole-body energy expenditure, food intake, locomotor activity, insulin tolerance, and pyruvate tolerance (Figure 5H, 6C, 6J, Figure S7G, S7K, S7L, S8A-D, S9E, S10B). Two-way repeated-measures ANOVA with Holm-Šídák’s multiple comparisons test was performed for gluconeogenesis assay (Figure S9F). For isotopic tracing studies, ANOVA with appropriate post-hoc tests was performed on ALR-transformed data, using M+0 as the denominator (one-way ANOVA with Tukey’s multiple comparisons test: Figure S1B-D; two-way ANOVA with Tukey’s multiple comparisons test: Figure 1B, 1C, Figure S1A, S1E, S1F; two-way ANOVA with Šídák’s multiple comparisons test: Figure 2G, Figure S3F-K). For metabolomics and lipidomics data, unpaired t-test was performed on log_10_-transformed values (Figure 2C-E, 6H, Figure S3B-D, S10C). For RNA-seq data, unpaired t-test was performed on log₂(TPM+0.01) (Figure 1D, 6A, 6B). For microarray data, one-way ANOVA with Holm-Šídák’s multiple comparisons test was performed on log₂-transformed values (Figure S2F). The sample numbers are provided in the figure legends. P < 0.05 was considered to be significant throughout the study.

---

**Figure S1.**
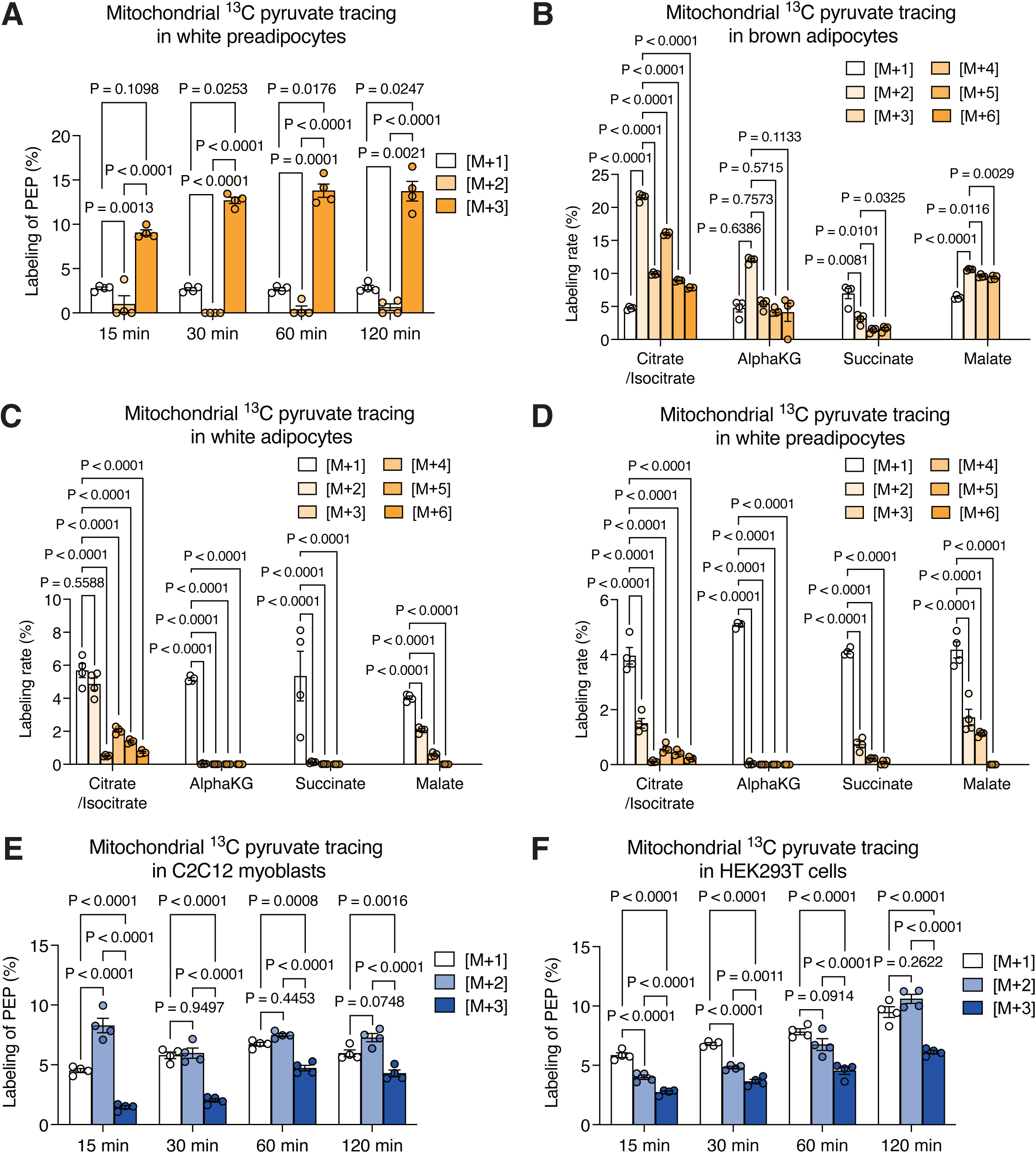
Mitochondrial pyruvate tracing studies in adipocytes, related to Figure 1. (A) Mitochondria from white preadipocytes were incubated with 13C3-pyruvate (10 mM) for the indicated time at 4°C to quantify labeled PEP by LC-MS. n = 4. (B) Mitochondria from brown adipocytes were incubated with 13C3-pyruvate (10 mM) for 15 min to quantify the labeled TCA metabolites (see Figure 1C). n = 4. (C) Mitochondria from white adipocytes were incubated with 13C3-pyruvate (10 mM) for 15 min to quantify the labeled TCA metabolites (see Figure 1B). n = 4. (D) Mitochondria from white preadipocytes were incubated with 13C3-pyruvate (10 mM) for 15 min to quantify the labeled TCA metabolites (see Figure S1A). n = 4. (E) Mitochondria from C2C12 myoblasts were incubated with 13C3-pyruvate (10 mM) for the indicated time at 4°C to quantify labeled PEP by LC-MS. n = 4. (F) Mitochondria from HEK293T cells were incubated with 13C3-pyruvate (10 mM) for the indicated time at 4°C to quantify labeled PEP by LC-MS. n = 4

**Figure S2.**
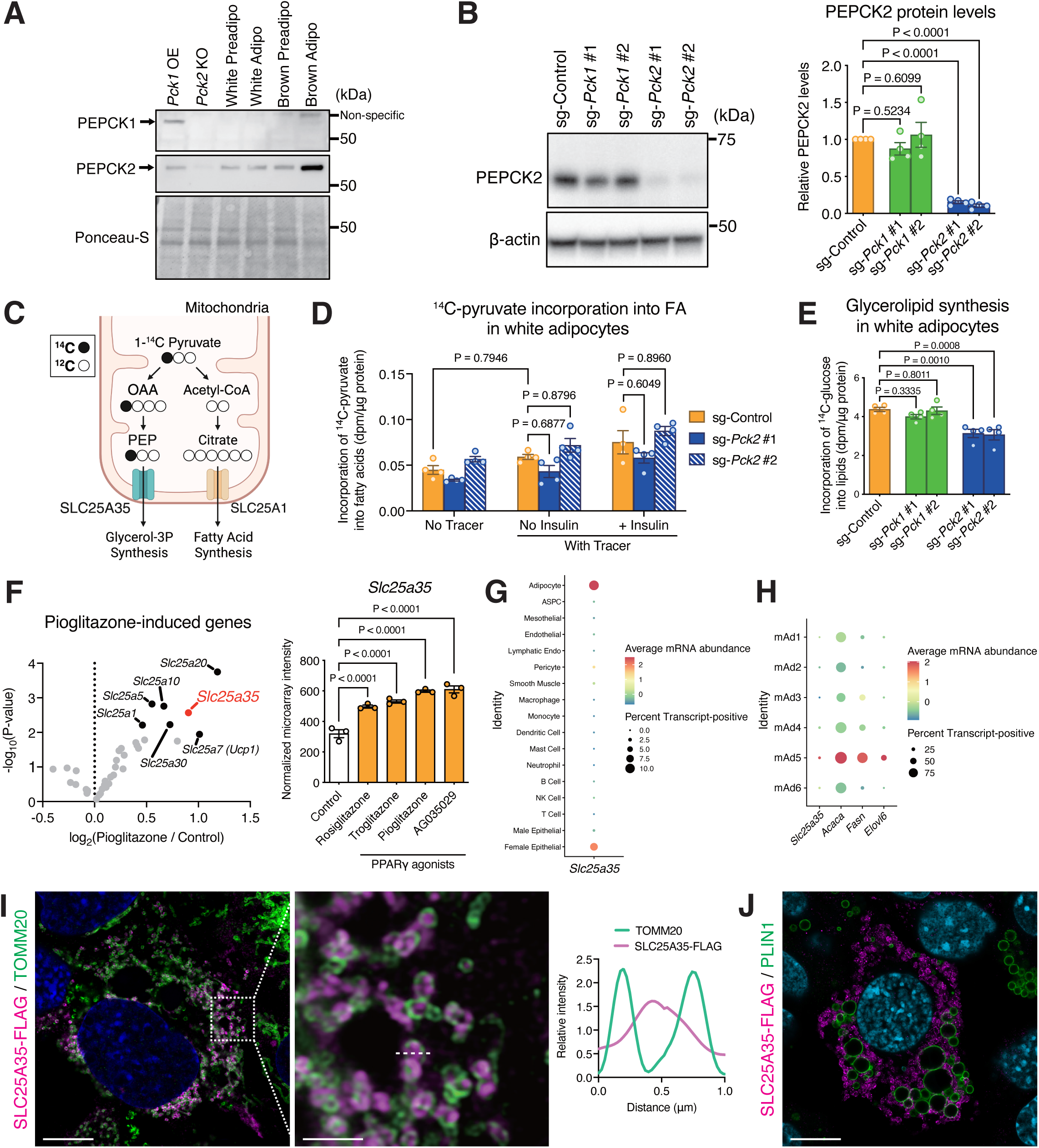
Mitochondrial carrier proteins that are linked to mitochondrial PEP synthesis, related to Figure 1. (A) Immunoblotting in differentiated white and brown adipocytes at day 6 and 4, respectively. *Pck1* overexpression and *Pck2* #1 sgRNA served as controls. (B) Immunoblotting of white preadipocytes transduced with *Pck1* or *Pck2* sgRNAs. Relative PEPCK2 protein levels to β-actin are quantified on the right. n = 4. (C) Schematic of the mitochondrial metabolic fate of 1-^14^C pyruvate. Labeled PEP is generated from 1-^14^C pyruvate, whereas citrate is not directly labeled. (D) Incorporation of 1-^14^C pyruvate into fatty acyl moiety of glycerolipids in white adipocytes transduced with *Pck2* sgRNAs in the presence or absence of insulin for 8 hours. Lipids were extracted and saponified, and ^14^C radioactivity (dpm) in the fatty acid fraction was quantified by TLC. n = 4. (E) Incorporation of ^14^C_6_ glucose into glycerolipids in white adipocytes transduced with either *Pck1* or *Pck2* sgRNAs in the presence of insulin for 8 hours. n = 4. (F) Volcano plot of SLC25A-family transcript changes in epididymal WAT from HFD-fed rats with or without pioglitazone. Normalized microarray intensity of *Slc25a35* across the indicated PPARγ agonists is shown on the right. Data are from microarray dataset GSE21329. n = 3. (G) Dot plot showing *Slc25a35* transcript abundance across cell types in mouse white adipose tissue from snRNA-seq dataset GSE176171. (H) Dot plot showing *Slc25a35*, *Acaca*, *Fasn*, and *Elovl6* mRNA abundance across six mouse white adipocyte subpopulations in snRNA-seq dataset GSE176171. (I) Representative immunofluorescence image of white adipocytes at day 3 with SLC25A35-FLAG, TOMM20, and DAPI. Scale bars, 10 μm (Left) and 2 μm (Right). (J) Representative immunofluorescence image of white adipocytes at day 3 with SLC25A35-FLAG, PLIN1, and DAPI. Scale bars, 10 μm.

**Figure S3.**
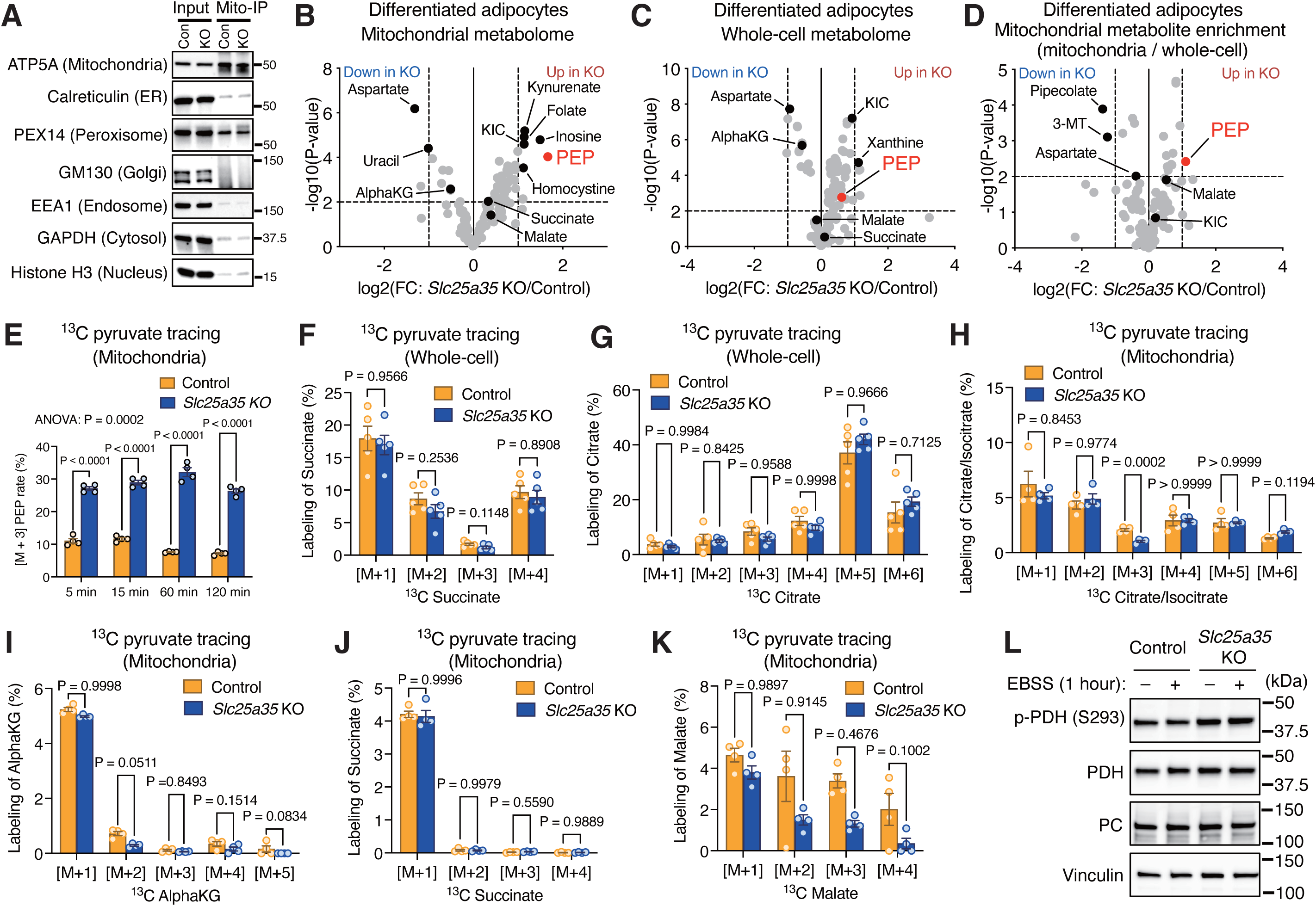
SLC25A35 deletion results in mitochondrial accumulation of PEP, related to Figure 2. (A) Immunoblotting of MITO-Tag–isolated mitochondria from white preadipocytes for the indicated organelle marker proteins. (B) Mitochondrial metabolomics of differentiated white adipocytes at day 6. The intensity was normalized to protein levels in each sample. n = 5. (C) Whole-cell metabolomics of differentiated white adipocytes at day 6 in (B). The intensity was normalized to protein levels in each sample. n = 5. (D) Mitochondrial metabolite enrichment was calculated as the ratio of mitochondrial to whole-cell intensity for each metabolite in (B) and (C). n = 5. (E) Time-course of mitochondrial ^13^C_3_ -pyruvate tracing in white preadipocytes to quantify M+3 PEP. n = 4. (F) Whole-cell ^13^C_3_ -pyruvate tracing in white preadipocytes to quantify labeled succinate. n = 5. (G) Whole-cell ^13^C_3_ -pyruvate tracing in white preadipocytes to quantify labeled citrate in (F). n = 5. (H) Mitochondrial ^13^C_3_ -pyruvate tracing to quantify labeled citrate/isocitrate at 15 min in (E). n = 4. (I) Mitochondrial ^13^C_3_ -pyruvate tracing to quantify labeled alpha-KG at 15 min in (E). n = 4. (J) Mitochondrial ^13^C_3_ -pyruvate tracing to quantify labeled succinate at 15 min in (E). n = 4. (K) Mitochondrial ^13^C_3_ -pyruvate tracing to quantify labeled malate at 15 min in (E). n = 4. (L) Immunoblotting of indicated proteins in white preadipocytes with or without one-hour starvation with EBSS.

**Figure S4.**
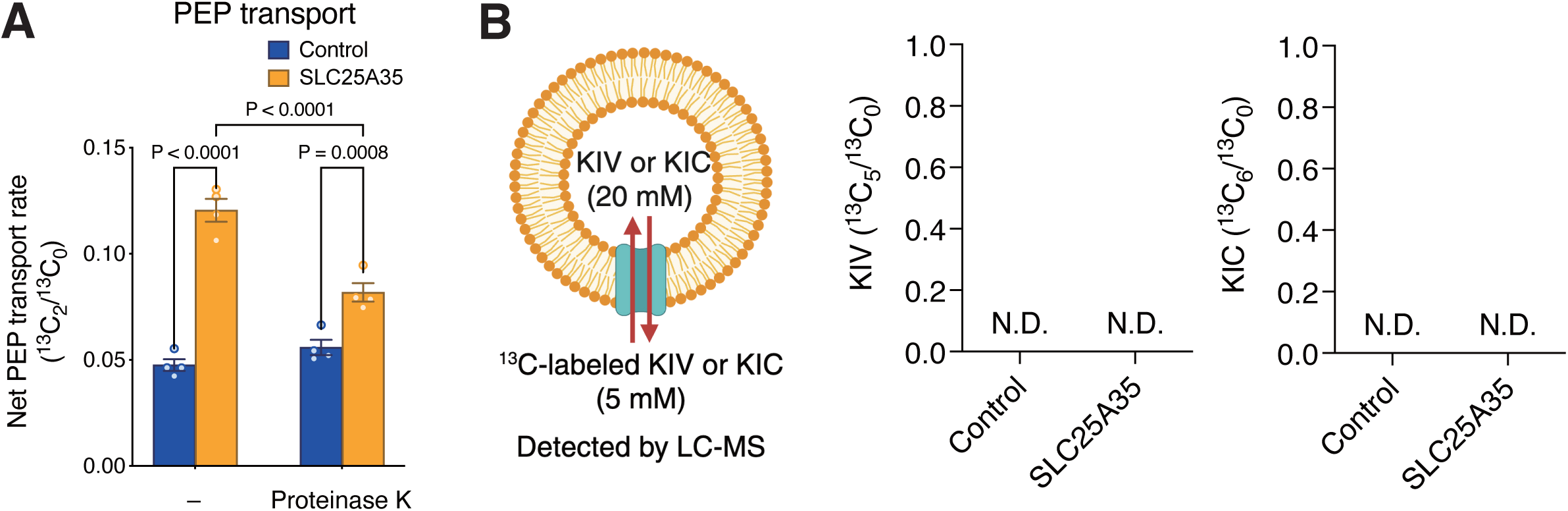
The substrate specificity of SLC25A35, related to Figure 3. (A) PEP transport assay using proteo-liposomes pretreated with or without proteinase K (50 μg/mL) for 10 min at 4°C prior to the transport assay. n = 4. (B) Schematic of KIV/KIC transport assays. Purified SLC25A35 protein or an empty-vector control eluate was reconstituted into liposomes containing unlabeled KIV/KIC (20 mM). SLC25A35 or control liposomes were incubated with fully 13C-labeled KIV/KIC (5 mM) for 20 min. KIV/KIC was not detected by LC-MS under the same analytical conditions used for the PEP assay, as shown on the right.

**Figure S5.**
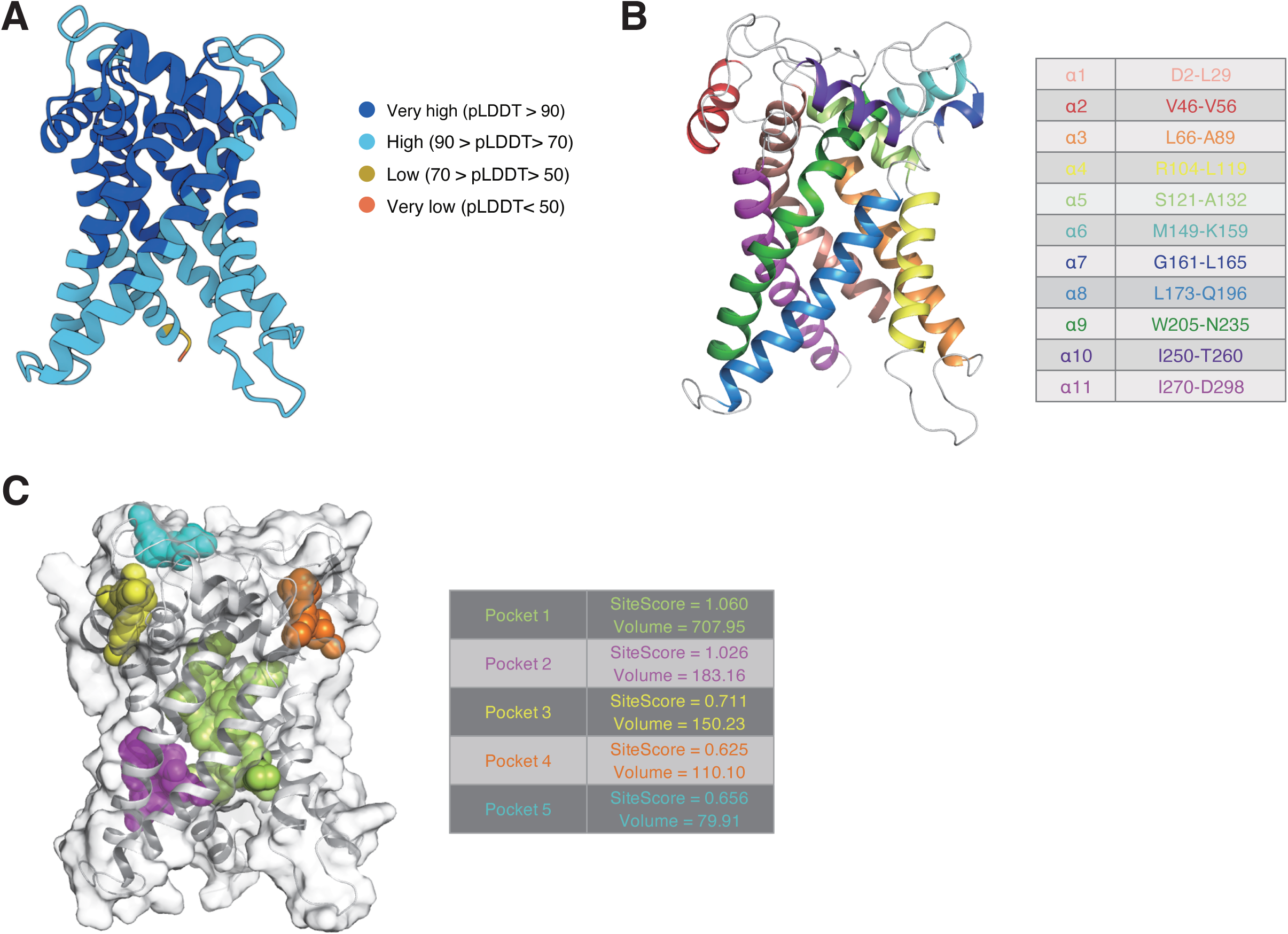
Structure of SLC25A35 predicted by AlphaFold, related to Figure 4. (A) AlphaFold SLC25A35 model (AF-Q3KQZ1-F1-model_v4) shows predominantly very high or confident per-residue pLDDT scores (pLDDT > 90 or 90–70), except for residues D298, T299, and K300 with pLDDT of 66.64, 62.10, and 46.45, respectively. The pLDDT for putative substrate-binding site residues A10, Y72, Q73, M76, N77, R80, A117, Y124, K127, R175, V176, S180, M223, R276, L277, and H280 are 87.52, 93.85, 90.99, 87.90, 85.36, 86.20, 90.86, 94.61, 95.99, 90.24, 88.90, 86.03, 93.42, 89.62, 84.53, and 88.56, respectively. Mean pLDDT is 89.25 for the full model and 89.66 for the binding site residues. (B) Cartoon/ribbon representation of the predicted SLC25A35 structure highlighting the 11 transmembrane α-helices (six vertical), colored by residue range. (C) Surface and cartoon (in white) representation of the predicted SLC25A35 structure with the cavities (pockets) identified by SiteMap.

**Figure S6.**
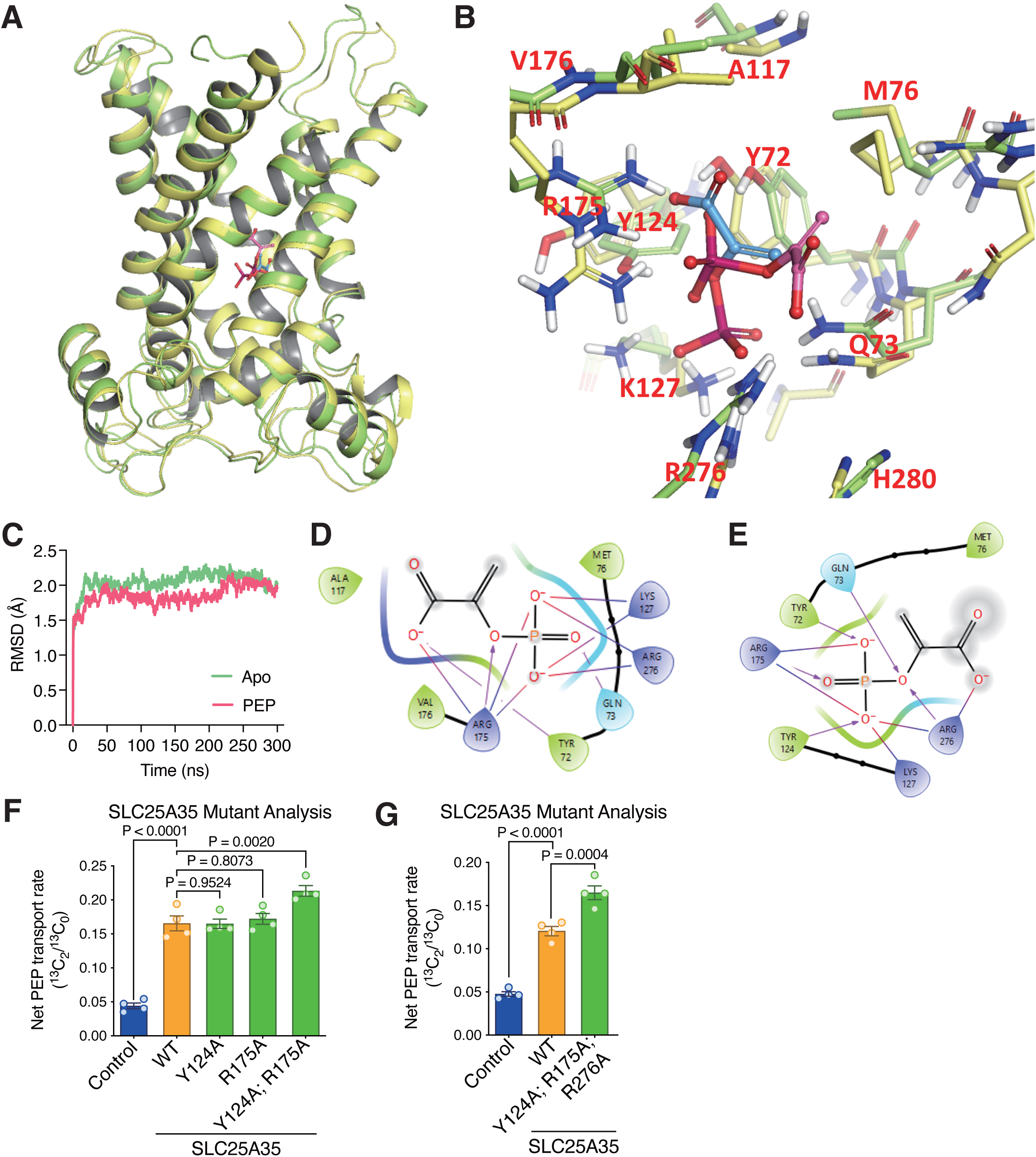
Molecular interactions between SLC25A35 and PEP by MD simulation, related to Figure 4. (A) Side view of the SLC25A35-PEP complex. The “before” structure corresponds to the best-scoring induced fit docking pose and is shown in protein (yellow) and PEP (blue), while the “after” structure is shown in protein (green) and PEP (magenta). (B) Close-up view of the SLC25A35–PEP binding site showing minor shifts in PEP position and overlapping residues before and after MD simulation. Key interacting residue pairs are labeled in red. (C) Backbone Root-Mean-Square Deviation (RMSD) from MD simulations of SLC25A35 in the apo state (blue) and in complex with PEP (orange). (D) 2D interaction diagram of PEP in the docking pose (before MD simulation). Magenta arrows indicate hydrogen bonds; blue-to-red lines indicate salt bridges. (E) 2D interaction diagram for PEP after MD simulation. Magenta arrows indicate hydrogen bonds; blue-to-red lines indicate salt bridges. (F) PEP transport assay with wild-type and indicated SLC25A35 mutants. n = 4. (G) PEP transport assay with wild-type and indicated SLC25A35 mutants. The control and wild-type SLC25A35 samples are identical to those in Figure S4A. n = 4.

**Figure S7.**
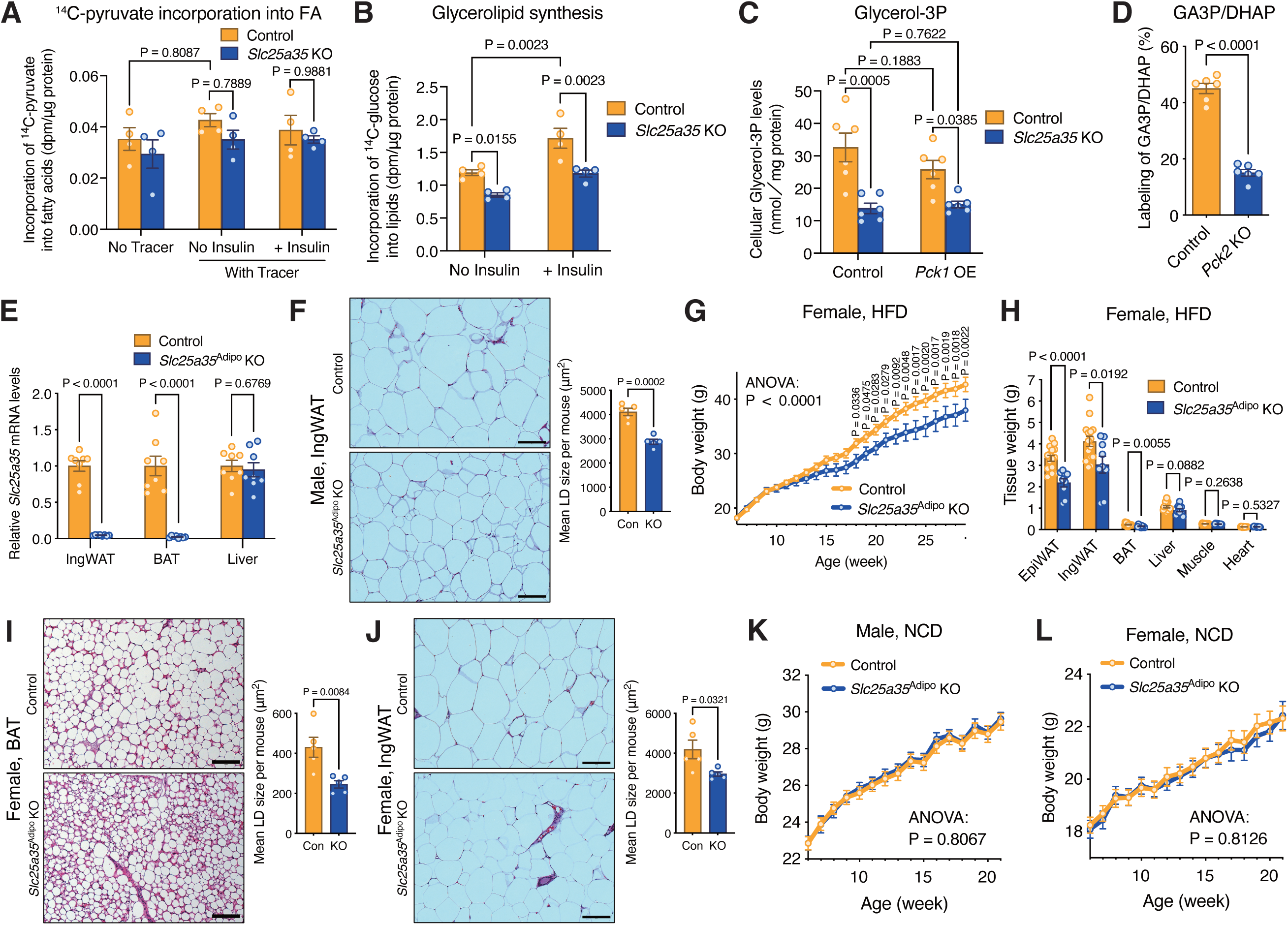
SLC25A35 is required for glyceroneogenesis in adipose tissue, related to Figure 5. (A) Incorporation of 1-^14^C pyruvate into fatty acyl moiety of glycerolipids in control or *Slc25a35* KO white adipocytes in the presence or absence of insulin for 8 hours. Lipids were extracted and saponified, and ^14^C radioactivity (dpm) in the fatty acid fraction was quantified by TLC. n = 4. (B) Incorporation of ^14^C_6_ glucose into glycerolipids in control and *Slc25a35* KO white adipocytes in the presence or absence of insulin for 8 hours. n = 4. (C) Cellular glycerol-3P levels in control or *Slc25a35* KO white adipocytes with control or mouse *Pck1* overexpression. n = 6. (D) Whole-cell ^13^C_3_ -pyruvate tracing in white preadipocytes transduced with control or *Pck2* #1 sgRNAs to quantify labeled GA3P/DHAP. n = 6. (E) Relative *Slc25a35* mRNA levels in the indicated tissues of male control and *Slc25a35*^Adipo^ KO mice on an HFD for 23 weeks at 30°C, measured by qPCR. n = 8. (F) Representative H&E-stained IngWAT sections from male mice in Figure 5I. Scale bars, 100 μm. Mean LD size per mouse is quantified on the right. n = 5. (G) Body weight changes in female control and *Slc25a35*^Adipo^ KO mice on an HFD at 30°C. n = 15 (control), 9 (KO). (H) Tissue weight of female mice in (G) after 23 weeks of HFD. n = 15 (control), 9 (KO). (I) Representative H&E-stained interscapular BAT sections from female mice in (H). Scale bars, 100 μm. Mean LD size per mouse is quantified on the right. n = 5. (J) Representative H&E-stained IngWAT sections from female mice in (H). Scale bars, 100 μm. Mean LD size per mouse is quantified on the right. n = 5. (K) Body weight changes in male control and *Slc25a35*^Adipo^ KO mice on a normal chow diet (NCD) at room temperature (RT). n = 16 (control), 21 (KO). (L) Body weight changes in female control and *Slc25a35*^Adipo^ KO mice on a NCD at RT. n = 21 (control), 15 (KO).

**Figure S8.**
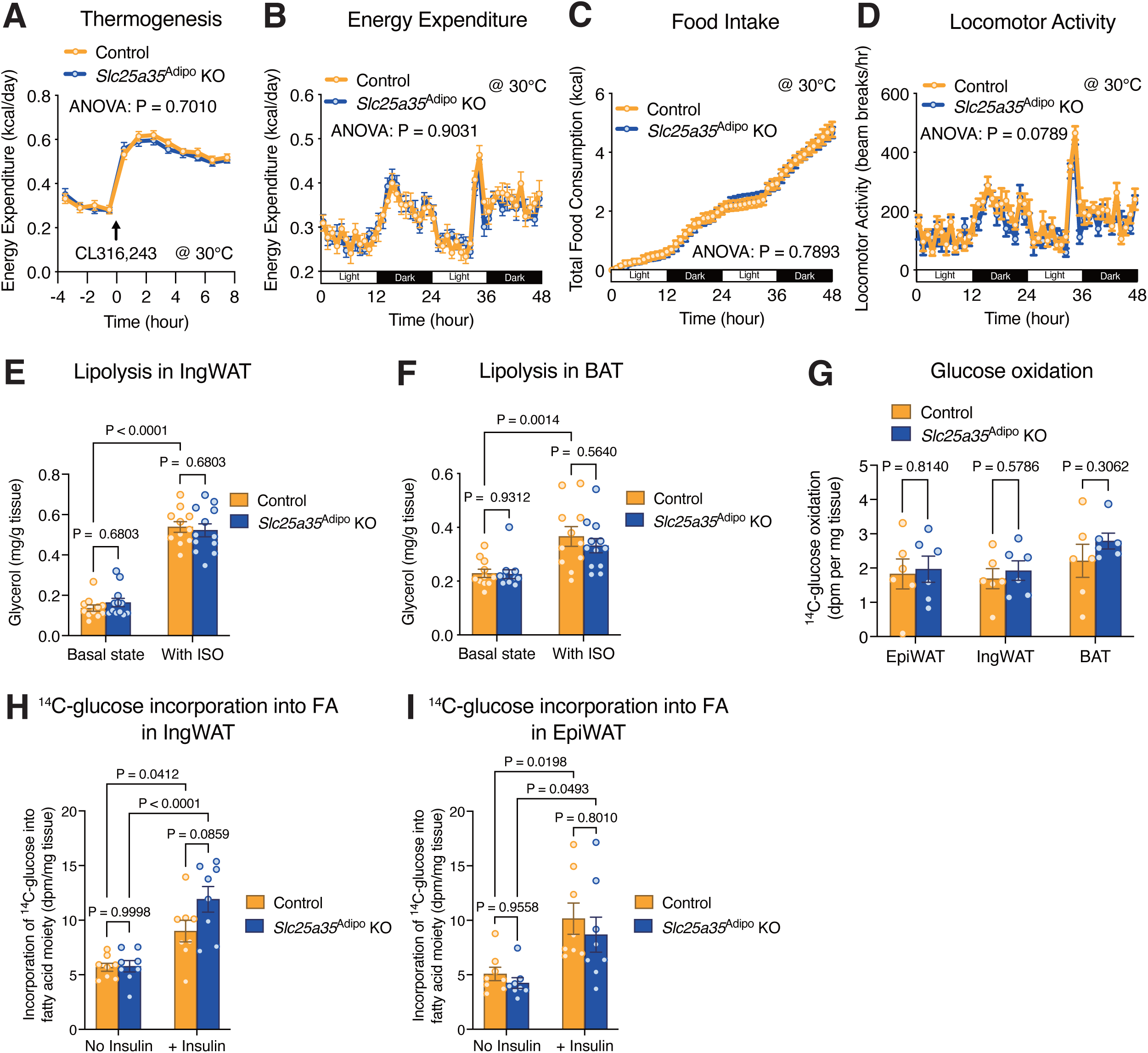
Metabolic characterization of Slc25a35Adipo KO mice, related to Figure 5. (A) Whole-body energy expenditure of male control and *Slc25a35*^Adipo^ KO mice on an HFD for 10 weeks, measured using the Comprehensive Lab Animal Monitoring System (CLAMS). Mice received an intraperitoneal injection of CL316,243 (0.5 mg kg-1) at 0 hour (arrow) at 30°C. n = 8. (B) Whole-body energy expenditure of male control and *Slc25a35*^Adipo^ KO mice, measured for 48 hours at 30°C using CLAMS. n = 8. (C) Total food consumption of male control and *Slc25a35*^Adipo^ KO mice in (B). n = 8. (D) Locomotor activity of male control and *Slc25a35*^Adipo^ KO mice in (B). n = 8. (E) Lipolysis assay in IngWAT of HFD-fed male control and *Slc25a35*^Adipo^ KO mice with or without isoproterenol (ISO). n = 11 (control), 12 (KO). (F) Lipolysis assay in interscapular BAT of HFD-fed male control and *Slc25a35*^Adipo^ KO mice with or without isoproterenol (ISO). n = 11 (control), 12 (KO). (G) Glucose oxidation assay in the indicated tissues from male control and *Slc25a35*^Adipo^ KO mice on an HFD. n = 6. (H) Incorporation of ^14^C_6_ -glucose into the fatty acid fraction of glycerolipids in IngWAT explants from HFD-fed male control and *Slc25a35*^Adipo^ KO mice for 4 hours with or without insulin. Lipids were extracted and saponified, and ^14^C radioactivity (dpm) in the fatty acid fraction was quantified by TLC. n = 8. (I) Incorporation of ^14^C_6_-glucose into the fatty acid fraction of glycerolipids in EpiWAT explants from HFD-fed male mice, performed as in (H). n = 8.

**Figure S9.**
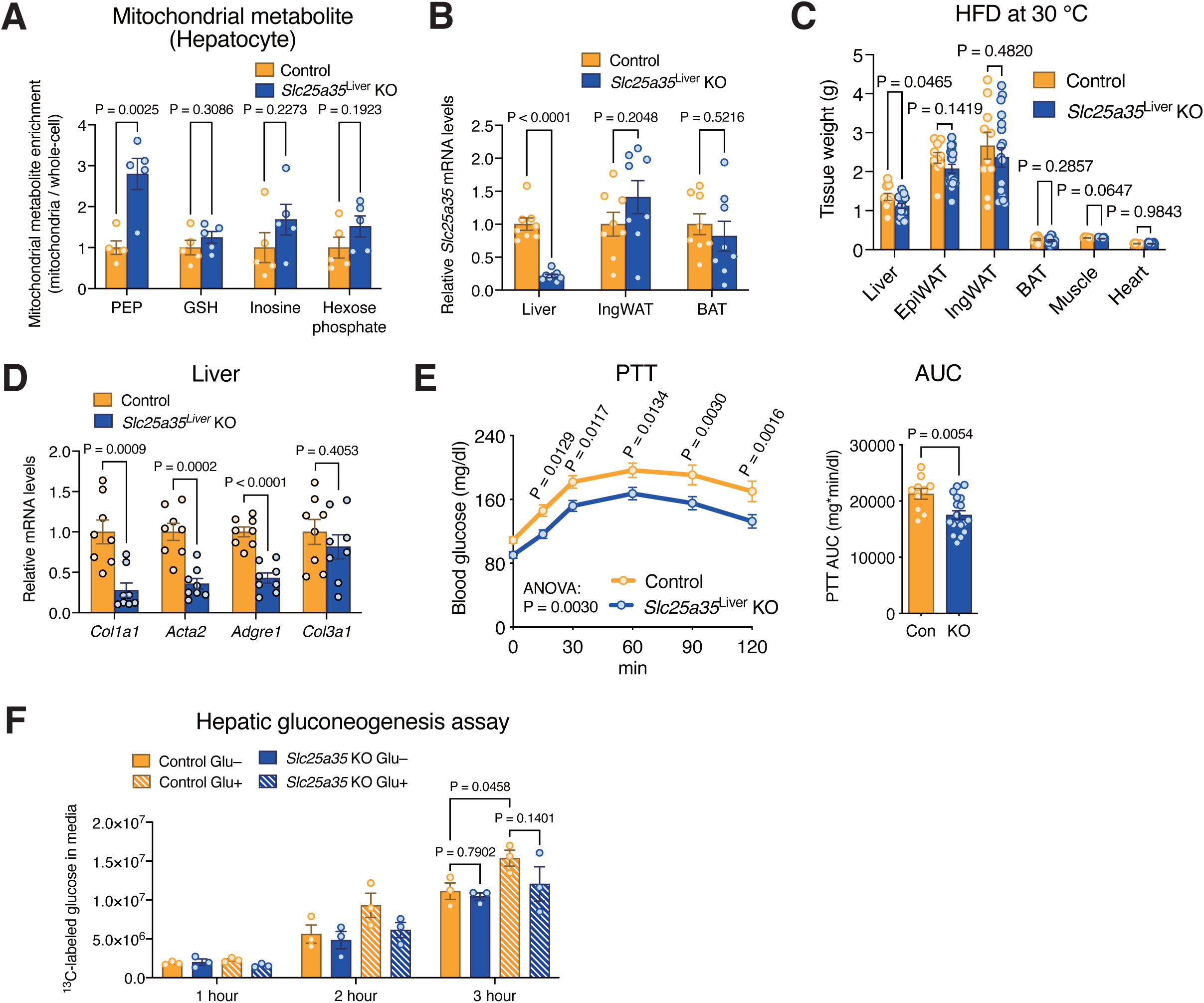
Blockade of mitochondrial PEP transport in the liver ameliorates hepatic steatosis, related to Figure 6. (A) Mitochondrial metabolite enrichment in primary hepatocytes from control and *Slc25a35*^Liver^ KO mice. Mitochondrial metabolite enrichment was calculated as the ratio of mitochondrial to whole-cell intensity for each metabolite. n = 5. (B) Relative *Slc25a35* mRNA levels in the indicated tissues from male control and *Slc25a35*^Liver^ KO mice on an HFD, measured by qPCR. n = 8. (C) Tissue weight of male control and *Slc25a35*^Liver^ KO mice on an HFD for 19 weeks at 30°C. n = 10 (control), 18 (KO). (D) Relative mRNA levels of pro-inflammatory and fibrosis genes in the liver of male control and *Slc25a35*^Liver^ KO mice on an HFD. n = 8. (E) Pyruvate tolerance test (PTT) of male control and *Slc25a35*^Liver^ KO mice on an HFD for 17 weeks. AUC is shown on the right. n = 10 (control), 18 (KO). (F) Gluconeogenesis assay in primary hepatocytes from control and *Slc25a35*^Liver^ KO mice with or without glucagon (100 nM). n = 3.

**Figure S10.**
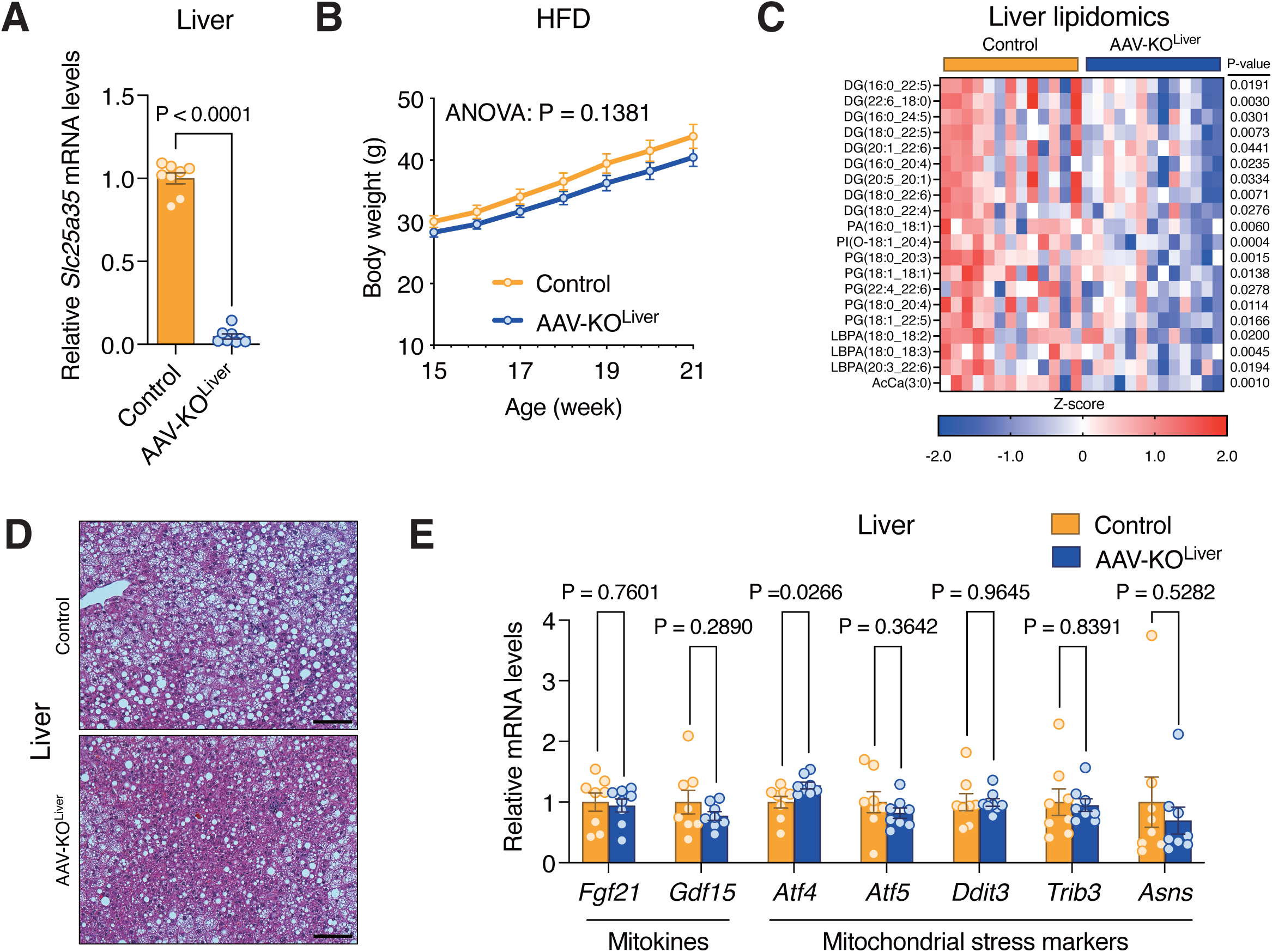
AAV-based depletion of mitochondrial PEP transport in the liver ameliorates hepatic steatosis, related to Figure 6. (A) Relative *Slc25a35* mRNA levels in the livers from male control and AAV-KO^Liver^ mice on an HFD. n = 8. (B) Body weight changes in male control and AAV-KO^Liver^ mice on an HFD. n = 13. (C) Liver lipidomics in male control and AAV-KO^Liver^ mice on an HFD. The top 20 lipid species significantly reduced in KO mice are shown. n = 13. (D) Representative H&E-staining of the liver from male control and AAV-KO^Liver^ mice on an HFD. Scale bars, 100 μm. LD area (%) is quantified in Figure 6M. (E) Relative mRNA levels of indicated genes in the liver of male control and AAV-KO^Liver^ mice on an HFD. n = 8.

**Table S1.** Differential mitochondrial metabolite profiles between control and *Slc25a35* KO white preadipocytes (P-value < 0.05), related to Figure 2.

Metabolites in negative ion mode
| Metabolite | FC | log<sub>2</sub>(FC) | P-value | -log<sub>10</sub>(P-value) |
| --- | --- | --- | --- | --- |
| PEP | 11.313 | 3.5 | 5.05E-05 | 4.2963 |
| 1,3-BPG | 6.5276 | 2.7065 | 0.004467 | 2.35 |
| 2,3-BPG | 4.3703 | 2.1277 | 0.0031327 | 2.5041 |
| NAD<sup>+</sup> | 3.6488 | 1.8674 | 4.19E-09 | 8.3778 |
| IDP | 3.6437 | 1.8654 | 0.014107 | 1.8506 |
| Glutathione disulfide | 3.6231 | 1.8572 | 2.65E-05 | 4.5771 |
| NADP<sup>+</sup> | 2.4277 | 1.2796 | 0.002372 | 2.6249 |
| N-carbamoyl-L-aspartate | 2.2635 | 1.1786 | 0.026998 | 1.5687 |
| dGDP | 2.1671 | 1.1157 | 0.00034007 | 3.4684 |
| Myo-inositol | 1.989 | 0.99206 | 0.00015592 | 3.8071 |
| UDP-D-glucose | 1.9231 | 0.94344 | 0.00054668 | 3.2623 |
| Pantothenate | 1.6462 | 0.7191 | 0.031365 | 1.5036 |
| ADP | 1.4905 | 0.57584 | 0.037204 | 1.4294 |
| p-Hydroxybenzoate | 0.80804 | -0.3075 | 0.044113 | 1.3554 |
| Pyrophosphate | 0.78851 | -0.3428 | 0.0054717 | 2.2619 |
| 2-Isopropylmalic acid | 0.76574 | -0.38507 | 0.030711 | 1.5127 |
| dTTP | 0.75155 | -0.41206 | 0.048522 | 1.3141 |
| Glucose-1-phosphate | 0.75067 | -0.41374 | 0.043039 | 1.3661 |
| Lactate | 0.72554 | -0.46288 | 0.009162 | 2.038 |
| Oxaloacetate | 0.71312 | -0.48779 | 0.00047587 | 3.3225 |
| Ribose-phosphate | 0.71063 | -0.49283 | 0.0035489 | 2.4499 |
| Orotidine-5-phosphate | 0.69861 | -0.51743 | 0.04981 | 1.3027 |
| Hydroxyisocaproic acid | 0.68268 | -0.55071 | 0.0080463 | 2.0944 |
| Nicotinate | 0.6729 | -0.57154 | 0.0024572 | 2.6096 |
| Succinate | 0.67124 | -0.57511 | 3.60E-09 | 8.4434 |
| Methylmalonic acid | 0.67004 | -0.57767 | 0.00019697 | 3.7056 |
| Glycolate | 0.66775 | -0.58261 | 0.049519 | 1.3052 |
| UTP | 0.64346 | -0.63607 | 0.046267 | 1.3347 |
| O8P-O1P | 0.63406 | -0.6573 | 0.0068197 | 2.1662 |
| KIV | 0.63164 | -0.66284 | 0.00616 | 2.2104 |
| Indole-3-carboxylic acid | 0.622 | -0.68502 | 0.0029617 | 2.5285 |
| Citramalate | 0.59976 | -0.73753 | 1.13E-06 | 5.9456 |
| Hydroxyphenylpyruvate | 0.59918 | -0.73893 | 0.0082194 | 2.0852 |
| Atrolactic acid | 0.57759 | -0.79189 | 0.0078693 | 2.1041 |
| Shikimate | 0.57395 | -0.80101 | 0.00067513 | 3.1706 |
| SBP | 0.55034 | -0.86161 | 0.02785 | 1.5552 |
| Trehalose-6-phosphate | 0.54718 | -0.86991 | 0.011921 | 1.9237 |
| Cholic acid | 0.5348 | -0.90293 | 0.011627 | 1.9345 |
| 2-Dehydro-D-gluconate | 0.53194 | -0.91066 | 0.00015064 | 3.8221 |
| Kynurenic acid | 0.52527 | -0.92887 | 0.0005645 | 3.2483 |
| Xanthine | 0.52441 | -0.93122 | 0.0061272 | 2.2127 |
| Deoxyuridine | 0.50749 | -0.97856 | 0.0063001 | 2.2007 |
| Orotate | 0.50172 | -0.99503 | 0.0052227 | 2.2821 |
| Citrate | 0.49844 | -1.0045 | 0.033355 | 1.4768 |
| Isocitrate | 0.41563 | -1.2666 | 0.0025722 | 2.5897 |
| Hypoxanthine | 0.41501 | -1.2688 | 0.0032769 | 2.4845 |
| GlcNAc-1P | 0.35461 | -1.4957 | 0.00030243 | 3.5194 |

Metabolites in positive ion mode
| Metabolite | FC | log<sub>2</sub>(FC) | P-value | -log<sub>10</sub>(P-value) |
| --- | --- | --- | --- | --- |
| NAD+ | 3.8811 | 1.9565 | 6.89E-06 | 5.1617 |
| Phosphorylcholine | 3.4105 | 1.77 | 3.40E-05 | 4.468 |
| NADP+ | 3.2937 | 1.7197 | 0.0015624 | 2.8062 |
| Glutathione disulfide | 3.2288 | 1.691 | 0.007612 | 2.1185 |
| Adenosine | 2.4111 | 1.2697 | 7.67E-05 | 4.1152 |
| Nicotinamide | 2.1539 | 1.1069 | 0.00070315 | 3.153 |
| Creatine | 2.0111 | 1.008 | 0.004455 | 2.3511 |
| Glycerophosphocholine | 1.8389 | 0.87883 | 0.0061687 | 2.2098 |
| CMP | 1.8008 | 0.84866 | 0.026267 | 1.5806 |
| N-acetyl-L-alanine | 1.7339 | 0.79405 | 0.0029267 | 2.5336 |
| AMP | 1.6065 | 0.68396 | 0.0011597 | 2.9357 |
| Phenylalanine | 1.4827 | 0.56821 | 0.014325 | 1.8439 |
| FAD | 1.4631 | 0.54907 | 0.00010183 | 3.9921 |
| Cholesterol | 0.71542 | -0.48313 | 0.0012747 | 2.8946 |
| Putrescine | 0.71078 | -0.49252 | 0.0018243 | 2.7389 |
| Urea | 0.69688 | -0.52101 | 0.049519 | 1.3052 |
| Cysteine | 0.68876 | -0.53792 | 0.0076617 | 2.1157 |
| Betaine | 0.67314 | -0.57102 | 0.01466 | 1.8339 |
| Serine | 0.65871 | -0.60229 | 0.040778 | 1.3896 |
| Betaine aldehyde | 0.65224 | -0.61652 | 0.013633 | 1.8654 |
| Carnitine | 0.64586 | -0.63072 | 0.0052957 | 2.2761 |
| Tryptophan | 0.64472 | -0.63325 | 0.031959 | 1.4954 |
| Ethanolamine | 0.61506 | -0.70119 | 0.041701 | 1.3798 |
| N-acetyl-L-ornithine | 0.61126 | -0.71015 | 0.020839 | 1.6811 |
| Indole | 0.58228 | -0.78022 | 0.016577 | 1.7805 |
| N-acetyl spermine | 0.56878 | -0.81407 | 0.00041088 | 3.3863 |
| S-methyl-5-thioadenosine | 0.5678 | -0.81655 | 0.0050257 | 2.2988 |
| Spermidine | 0.5358 | -0.90025 | 3.05E-06 | 5.5155 |
| N-acetylputrescine | 0.52076 | -0.94131 | 0.016373 | 1.7859 |
| Methylnicotinamide | 0.42086 | -1.2486 | 0.00020289 | 3.6927 |

**Table S2.**
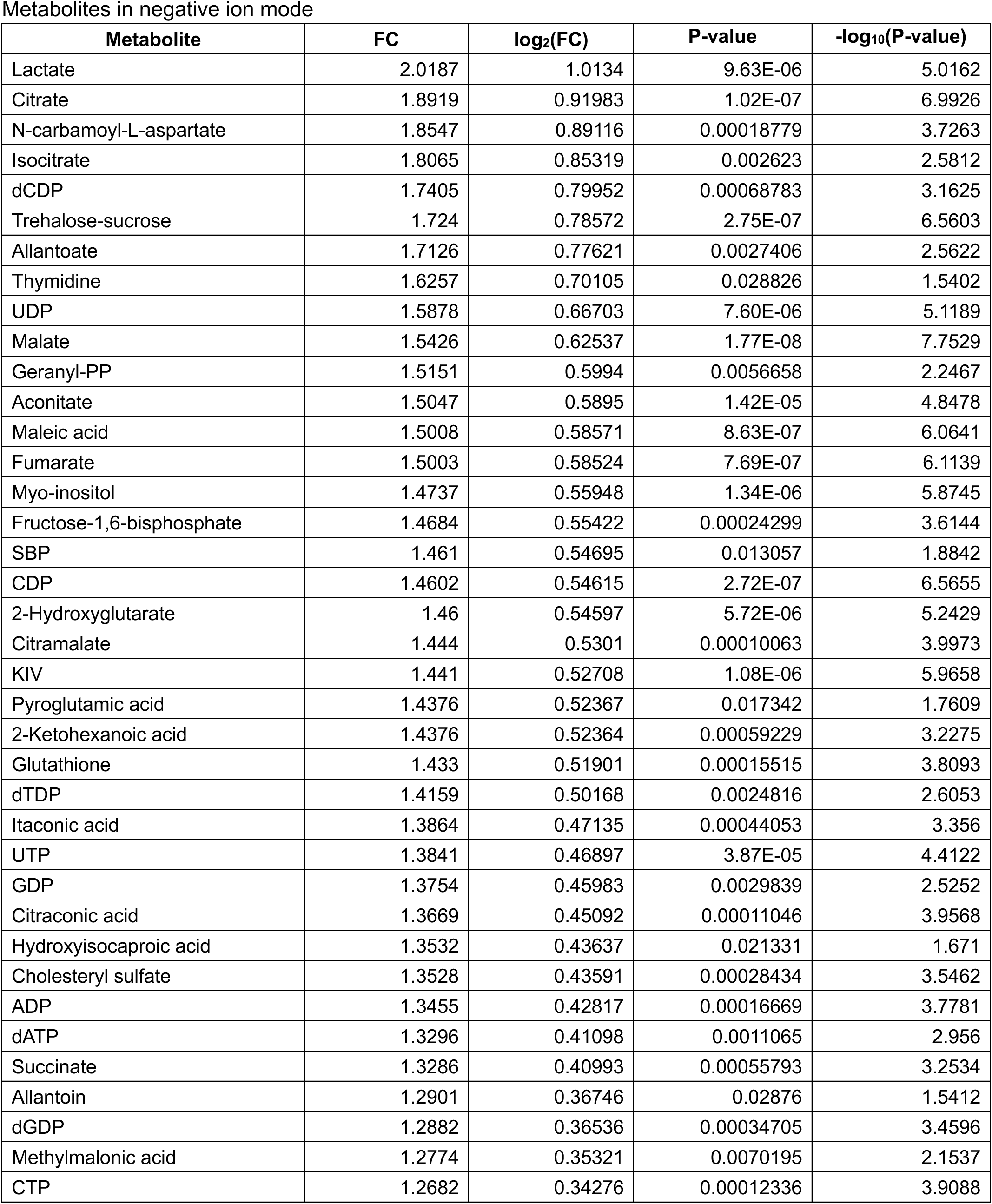
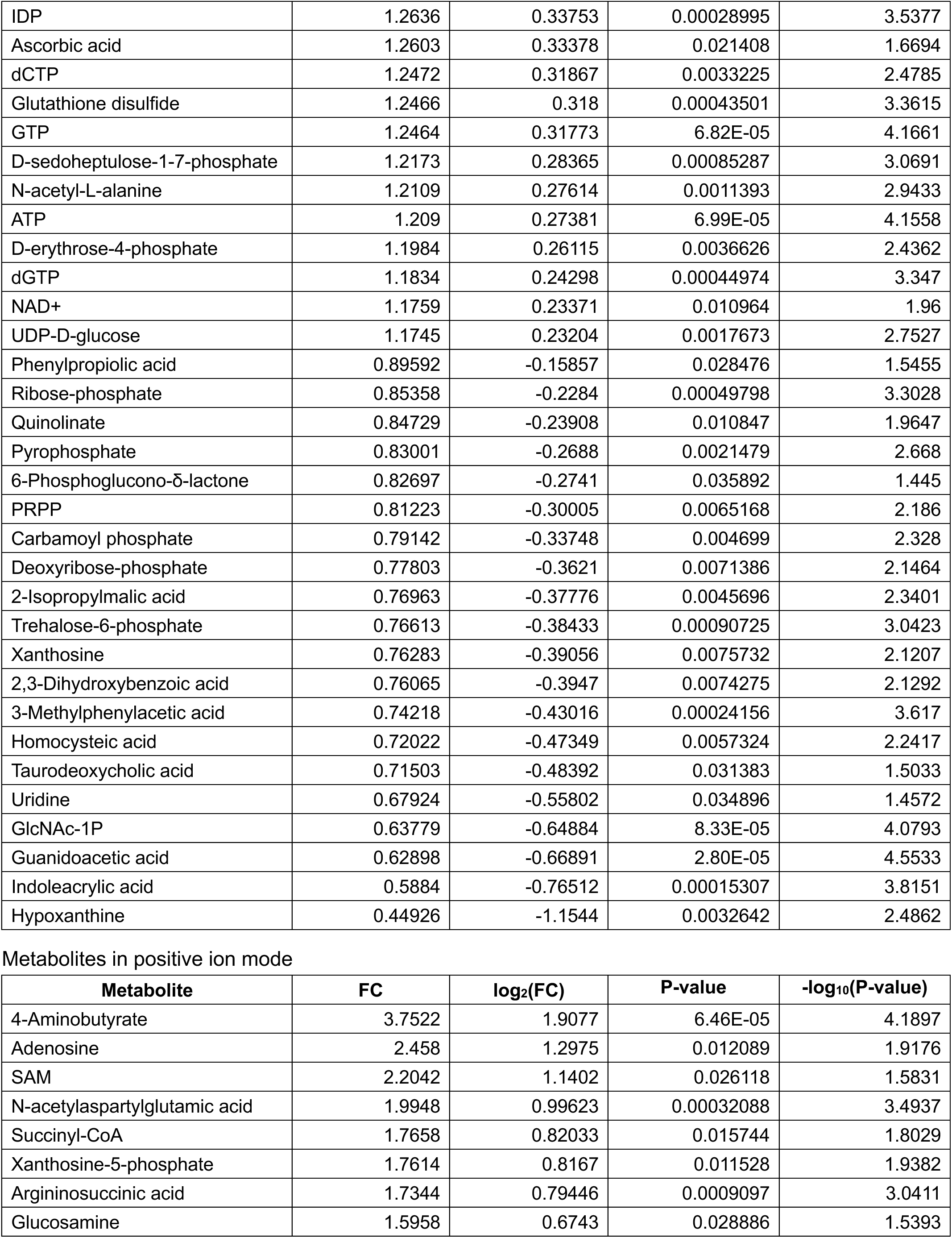
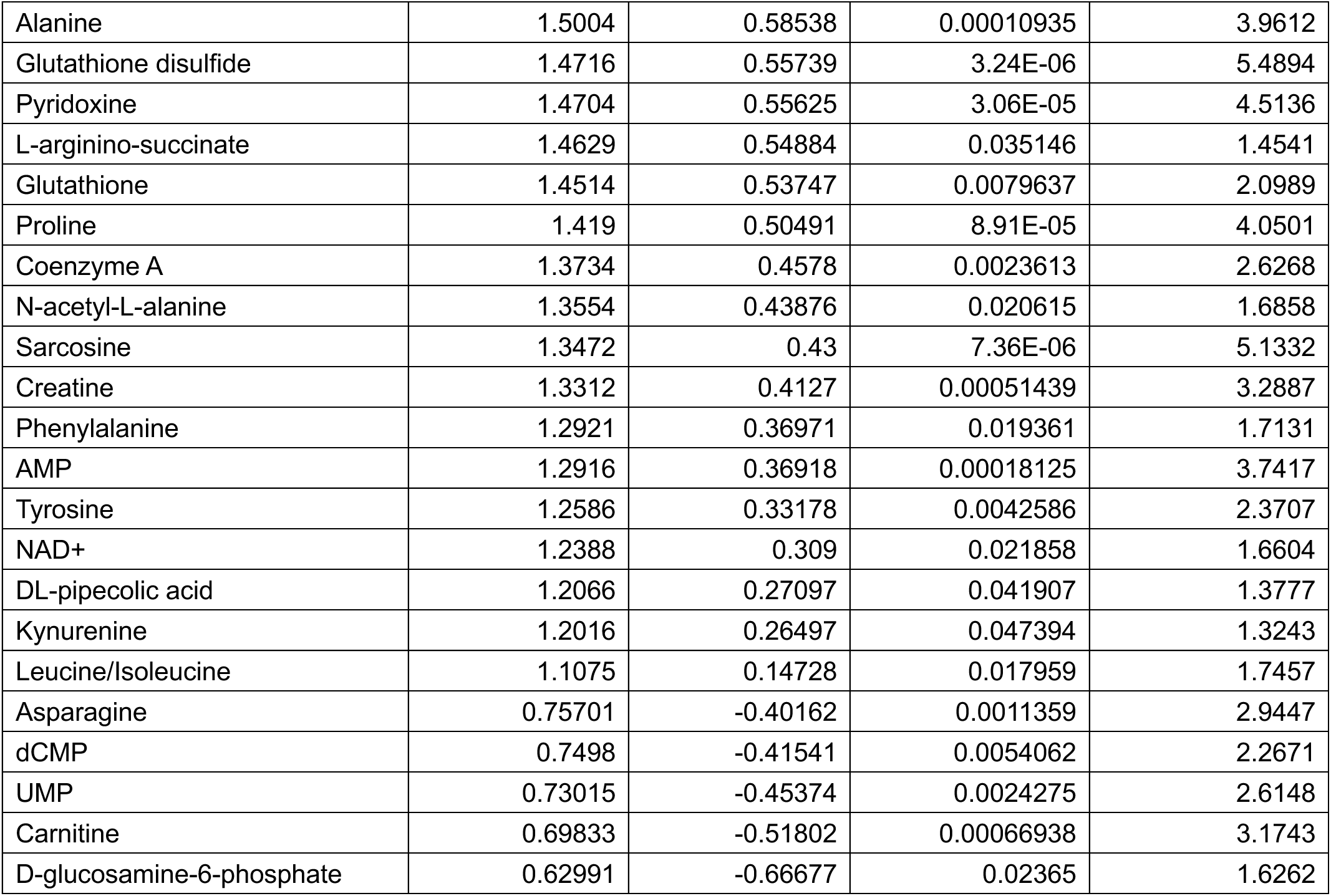
Differential whole-cell metabolite profiles between control and *Slc25a35* KO white preadipocytes (P-value < 0.05), related to Figure 2.

**Table S3.** Differential mitochondrial metabolite enrichment (mitochondria/whole-cell ratio) between control and *Slc25a35* KO white preadipocytes (P-value < 0.05), related to Figure 2.

Metabolites in negative ion mode
| Metabolite | FC | log<sub>2</sub>(FC) | P-value | -log<sub>10</sub>(P-value) |
| --- | --- | --- | --- | --- |
| PEP | 9.4068 | 3.2337 | 0.00019936 | 3.7004 |
| 1,3-BPG | 5.3866 | 2.4294 | 0.0039683 | 2.4014 |
| 2,3-BPG | 4.0233 | 2.0084 | 0.044286 | 1.3537 |
| NAD<sup>+</sup> | 3.1142 | 1.6388 | 7.98E-09 | 8.098 |
| Glutathione disulfide | 2.9733 | 1.5721 | 0.00031403 | 3.503 |
| Xanthosine | 2.806 | 1.4885 | 0.0043053 | 2.366 |
| NADP<sup>+</sup> | 2.3164 | 1.2119 | 0.0018933 | 2.7228 |
| dGDP | 1.7212 | 0.78343 | 0.013124 | 1.8819 |
| UDP-D-glucose | 1.6238 | 0.69937 | 0.0015652 | 2.8054 |
| Myo-inositol | 1.3535 | 0.43666 | 0.042123 | 1.3755 |
| 6-Phospho-D-gluconate | 1.3239 | 0.40478 | 0.0066092 | 2.1799 |
| Oxaloacetate | 0.73162 | -0.45084 | 0.022373 | 1.6503 |
| Nicotinate | 0.71749 | -0.47896 | 0.0045445 | 2.3425 |
| O8P-O1P | 0.70455 | -0.50523 | 0.023993 | 1.6199 |
| Itaconic acid | 0.70017 | -0.51421 | 0.00028494 | 3.5452 |
| UDP | 0.69962 | -0.51536 | 0.04024 | 1.3953 |
| dTTP | 0.68979 | -0.53577 | 0.025298 | 1.5969 |
| Cholesteryl sulfate | 0.68886 | -0.53773 | 0.0018983 | 2.7216 |
| dTDP | 0.68203 | -0.55208 | 0.030723 | 1.5125 |
| Glycerate | 0.67966 | -0.55711 | 0.034088 | 1.4674 |
| Trehalose-6-Phosphate | 0.66771 | -0.5827 | 0.039425 | 1.4042 |
| ATP | 0.66492 | -0.58875 | 0.039163 | 1.4071 |
| dGTP | 0.65891 | -0.60184 | 0.029265 | 1.5337 |
| GTP | 0.65798 | -0.60389 | 0.027675 | 1.5579 |
| Citraconic acid | 0.65674 | -0.6066 | 0.024393 | 1.6127 |
| Glutathione | 0.64267 | -0.63784 | 0.00017197 | 3.7646 |
| Phenylpyruvate | 0.63606 | -0.65276 | 0.012649 | 1.8979 |
| Aconitate | 0.63411 | -0.6572 | 0.002161 | 2.6653 |
| Phenyllactic acid | 0.624 | -0.68038 | 0.014099 | 1.8508 |
| 2-Hydroxyglutarate | 0.60667 | -0.72103 | 0.0015947 | 2.7973 |
| Xanthine | 0.59467 | -0.74985 | 0.008507 | 2.0702 |
| Shikimate | 0.59399 | -0.75148 | 0.003905 | 2.4084 |
| GlcNAc-1P | 0.59292 | -0.7541 | 0.01475 | 1.8312 |
| Indole-3-carboxylic acid | 0.58826 | -0.76547 | 0.002551 | 2.5933 |
| Hydroxyisocaproic acid | 0.56885 | -0.81389 | 0.0056825 | 2.2455 |
| Fructose-1,6-bisphosphate | 0.55688 | -0.84455 | 0.0045901 | 2.3382 |
| Maleic acid | 0.55063 | -0.86085 | 0.00045047 | 3.3463 |
| Atrolactic acid | 0.55044 | -0.86134 | 0.017777 | 1.7501 |
| Methylmalonic acid | 0.54837 | -0.86677 | 0.00024337 | 3.6137 |
| Fumarate | 0.54694 | -0.87054 | 0.0058505 | 2.2328 |
| Malate | 0.54081 | -0.88679 | 6.45E-05 | 4.1905 |
| Succinate | 0.51615 | -0.95414 | 3.22E-07 | 6.4926 |
| UTP | 0.47325 | -1.0793 | 0.0088739 | 2.0519 |
| Kynurenic acid | 0.45173 | -1.1465 | 0.0024091 | 2.6181 |
| KIV | 0.43613 | -1.1972 | 0.00012904 | 3.8893 |
| Citramalate | 0.42518 | -1.2339 | 3.95E-07 | 6.4037 |
| Lactate | 0.37 | -1.4344 | 1.26E-06 | 5.9009 |
| SBP | 0.3543 | -1.497 | 0.0022776 | 2.6425 |
| Citrate | 0.28809 | -1.7954 | 0.0047038 | 2.3276 |
| Isocitrate | 0.27756 | -1.8491 | 0.0013914 | 2.8566 |

Metabolites in positive ion mode
| Metabolite | FC | log<sub>2</sub>(FC) | P-value | -log<sub>10</sub>(P-value) |
| --- | --- | --- | --- | --- |
| NADP+ | 3.2416 | 1.6967 | 0.0016731 | 2.7765 |
| NAD+ | 3.111 | 1.6374 | 7.93E-06 | 5.101 |
| Nicotinamide | 2.8821 | 1.5271 | 0.0095716 | 2.019 |
| Phosphorylcholine | 2.7953 | 1.483 | 0.00062614 | 3.2033 |
| Glycerophosphocholine | 1.8672 | 0.90086 | 0.01373 | 1.8623 |
| FAD | 1.6007 | 0.67874 | 0.013781 | 1.8607 |
| Creatine | 1.508 | 0.59262 | 0.048826 | 1.3114 |
| Cholesterol | 0.74128 | -0.43191 | 0.0046586 | 2.3317 |
| Serine | 0.64465 | -0.63341 | 0.03916 | 1.4072 |
| Glutathione | 0.60286 | -0.7301 | 0.0071256 | 2.1472 |
| Tryptophan | 0.58244 | -0.77983 | 0.018957 | 1.7222 |
| N-acetyl-glutamine | 0.58025 | -0.78526 | 0.004202 | 2.3765 |
| N-acetyl spermidine | 0.57242 | -0.80487 | 0.013972 | 1.8547 |
| Spermidine | 0.53371 | -0.90588 | 0.0002391 | 3.6214 |
| Putrescine | 0.53232 | -0.90964 | 6.11E-05 | 4.2141 |
| Indole | 0.48249 | -1.0514 | 0.014877 | 1.8275 |
| S-methyl-5-thioadenosine | 0.44918 | -1.1546 | 0.0024362 | 2.6133 |
| Pyridoxine | 0.44319 | -1.174 | 0.0076904 | 2.1141 |

**Table S4.** Differential liver lipid profiles between control and *Slc25a35* KO^Liver^ mice (P-value < 0.05), related to Figure 6.

| Lipid | FC | log<sub>2</sub>(FC) | P-value | -log<sub>10</sub>(P-value) |
| --- | --- | --- | --- | --- |
| TG(40:7_16:0)+O:(s) | 0.57122 | -0.80789 | 0.026388 | 1.5786 |
| TG(14:0_18:2_22:6) | 0.60338 | -0.72886 | 0.033985 | 1.4687 |
| PG(30:0_16:0) | 0.60934 | -0.71467 | 0.03043 | 1.5167 |
| TG(20:4_18:2_18:2)+O:(s) | 0.6229 | -0.68292 | 0.0063108 | 2.1999 |
| TG(12:0_18:2_18:3) | 0.63048 | -0.66547 | 0.029964 | 1.5234 |
| d5-TG(32:5_22:6) | 0.63359 | -0.65838 | 0.045333 | 1.3436 |
| TG(28:3_16:0) | 0.63664 | -0.65146 | 0.039697 | 1.4012 |
| TG(10:0_16:0_20:4) | 0.63722 | -0.65014 | 0.041954 | 1.3772 |
| DG(16:1_20:4) | 0.64174 | -0.63994 | 0.0065615 | 2.183 |
| TG(16:0_16:3_22:6) | 0.64384 | -0.63524 | 0.046288 | 1.3345 |
| PG(O-24:0_18:2) | 0.6702 | -0.57734 | 0.021409 | 1.6694 |
| TG(18:1_18:2_18:2)+O:(s) | 0.67077 | -0.57612 | 0.015569 | 1.8078 |
| TG(15:0_18:2_20:5) | 0.68018 | -0.55601 | 0.04185 | 1.3783 |
| PG(O-28:0_22:6) | 0.68772 | -0.54011 | 0.023304 | 1.6326 |
| PEt(O-12:0_20:3) | 0.68937 | -0.53664 | 0.047624 | 1.3222 |
| TG(16:0_14:1_18:1)+O:(s) | 0.69006 | -0.53521 | 0.048572 | 1.3136 |
| TG(16:0_18:1_16:2)+O:(s) | 0.69458 | -0.52578 | 0.028204 | 1.5497 |
| TG(16:1_18:1_18:2)+O:(s) | 0.69633 | -0.52215 | 0.048166 | 1.3173 |
| TG(16:0_15:2_22:6) | 0.71458 | -0.48484 | 0.03246 | 1.4887 |
| PEt(23:6_18:0) | 0.72036 | -0.47321 | 0.046256 | 1.3348 |
| TG(18:1_18:1_16:3)+O:(s) | 0.72918 | -0.45566 | 0.047214 | 1.3259 |
| TG(16:0_22:6_22:6) | 0.73641 | -0.44142 | 0.041115 | 1.386 |
| Cer(d18:1_22:1) | 0.74454 | -0.42557 | 0.039551 | 1.4028 |
| TG(16:0_13:1_18:2) | 0.74636 | -0.42205 | 0.029561 | 1.5293 |
| PI(18:0_18:0) | 0.75487 | -0.40569 | 0.013035 | 1.8849 |
| PC(18:0_22:5) | 0.75559 | -0.40432 | 0.042893 | 1.3676 |
| PI(22:4_18:0) | 0.75739 | -0.4009 | 0.021613 | 1.6653 |
| TG(18:1_18:1_20:3) | 0.76028 | -0.3954 | 0.041778 | 1.3791 |
| Cer(d18:0_24:1) | 0.77365 | -0.37025 | 0.018319 | 1.7371 |
| LPS(18:0) | 0.77933 | -0.35968 | 0.036381 | 1.4391 |
| TG(18:1_18:1_20:1) | 0.79095 | -0.33834 | 0.038308 | 1.4167 |
| TG(18:1_15:2_18:2) | 0.7998 | -0.32229 | 0.035004 | 1.4559 |
| d5-TG(18:3_16:0_20:4) | 0.80708 | -0.30921 | 0.04071 | 1.3903 |
| Cer(d18:0_16:0) | 0.83643 | -0.25768 | 0.045802 | 1.3391 |
| TG(16:0_18:1_22:6) | 0.83997 | -0.25158 | 0.030713 | 1.5127 |
| TG(16:0_18:1_22:5) | 0.85085 | -0.23303 | 0.032551 | 1.4874 |
| d5-TG(36:5_20:2) | 0.85398 | -0.22772 | 0.023344 | 1.6318 |
| TG(36:7_25:7)+O:(s) | 1.0745 | 0.1036 | 0.0056993 | 2.2442 |
| TG(26:1_9:0)+OO:(s) | 1.1277 | 0.17338 | 0.041832 | 1.3785 |
| d5-TG(36:6_20:4) | 1.1279 | 0.1737 | 0.045671 | 1.3404 |
| SM(t34:1) | 1.1352 | 0.18301 | 0.041257 | 1.3845 |
| TG(14:0_16:0_18:1) | 1.1677 | 0.22372 | 0.04058 | 1.3917 |
| SM(d15:0_21:5) | 1.1745 | 0.23204 | 0.0051183 | 2.2909 |
| TG(16:0_16:0_18:1) | 1.1968 | 0.25923 | 0.04496 | 1.3472 |
| PEt(O-23:5_16:1) | 1.2206 | 0.28755 | 0.022309 | 1.6515 |
| SM(d44:6) | 1.2206 | 0.28756 | 0.048151 | 1.3174 |
| TG(13:1_21:0_21:0) | 1.228 | 0.2963 | 0.042382 | 1.3728 |
| DG(O-18:4_18:2) | 1.249 | 0.3208 | 0.0294 | 1.5316 |
| TG(18:2_18:2_22:5) | 1.2664 | 0.34077 | 0.038751 | 1.4117 |
| TG(30:1_9:0)+OO:(s) | 1.2701 | 0.3449 | 0.029242 | 1.534 |
| SM(d42:5) | 1.2955 | 0.37348 | 0.029677 | 1.5276 |
| PG(O-30:1_18:2) | 1.3074 | 0.38675 | 0.0097043 | 2.013 |
| SPH(d19:0) | 1.3115 | 0.39117 | 0.02762 | 1.5588 |
| PC(18:1_22:6) | 1.3118 | 0.39157 | 0.021946 | 1.6587 |
| TG(37:7_22:6) | 1.3224 | 0.40318 | 0.035872 | 1.4452 |
| PE(18:2_17:0) | 1.3506 | 0.43361 | 0.042375 | 1.3729 |
| DG(O-42:6) | 1.3612 | 0.44489 | 0.031779 | 1.4979 |
| d5-TG(36:5_22:6) | 1.3781 | 0.46266 | 0.0075338 | 2.123 |
| d5-DG(14:0_16:0) | 1.4375 | 0.52354 | 0.029879 | 1.5246 |
| PE(16:0_18:2) | 1.4598 | 0.54578 | 0.0090021 | 2.0457 |
| TG(34:2_6:0)+OO:(s) | 1.4681 | 0.55397 | 0.023077 | 1.6368 |
| ChE(22:6) | 1.4827 | 0.56825 | 0.04796 | 1.3191 |
| PE(22:6_15:0) | 1.5111 | 0.59558 | 0.0092944 | 2.0318 |
| AcCa(16:2) | 1.6365 | 0.71059 | 0.047258 | 1.3255 |
| SM(d41:6) | 1.6422 | 0.71561 | 0.0052426 | 2.2805 |
| ChE(18:2) | 1.9298 | 0.94848 | 0.0083663 | 2.0775 |
| TG(O-16:0_18:1_22:6) | 2.4262 | 1.2787 | 0.018623 | 1.73 |
| TG(23:0_18:1_18:1) | 2.7218 | 1.4446 | 0.022076 | 1.6561 |

**Table S5.**
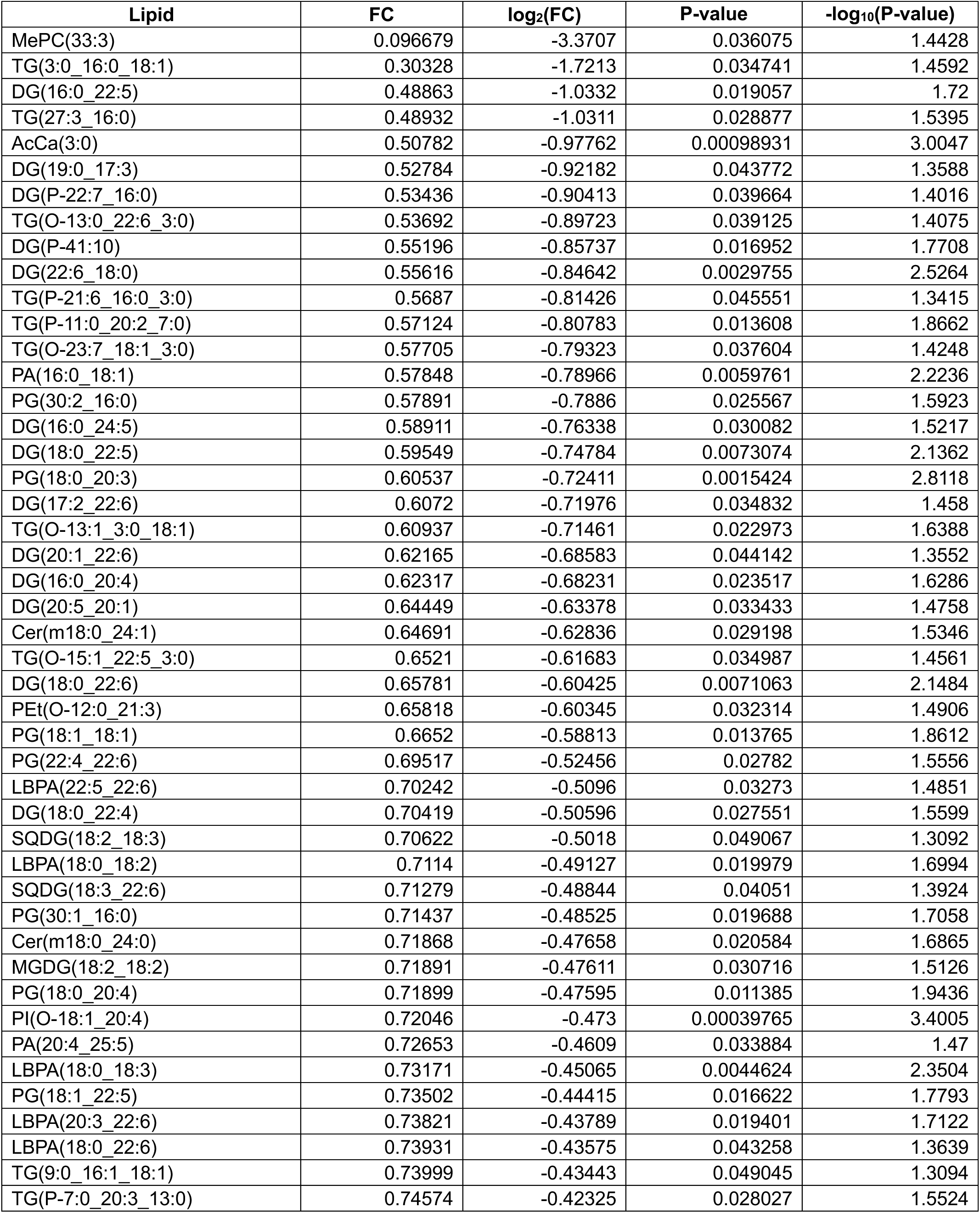
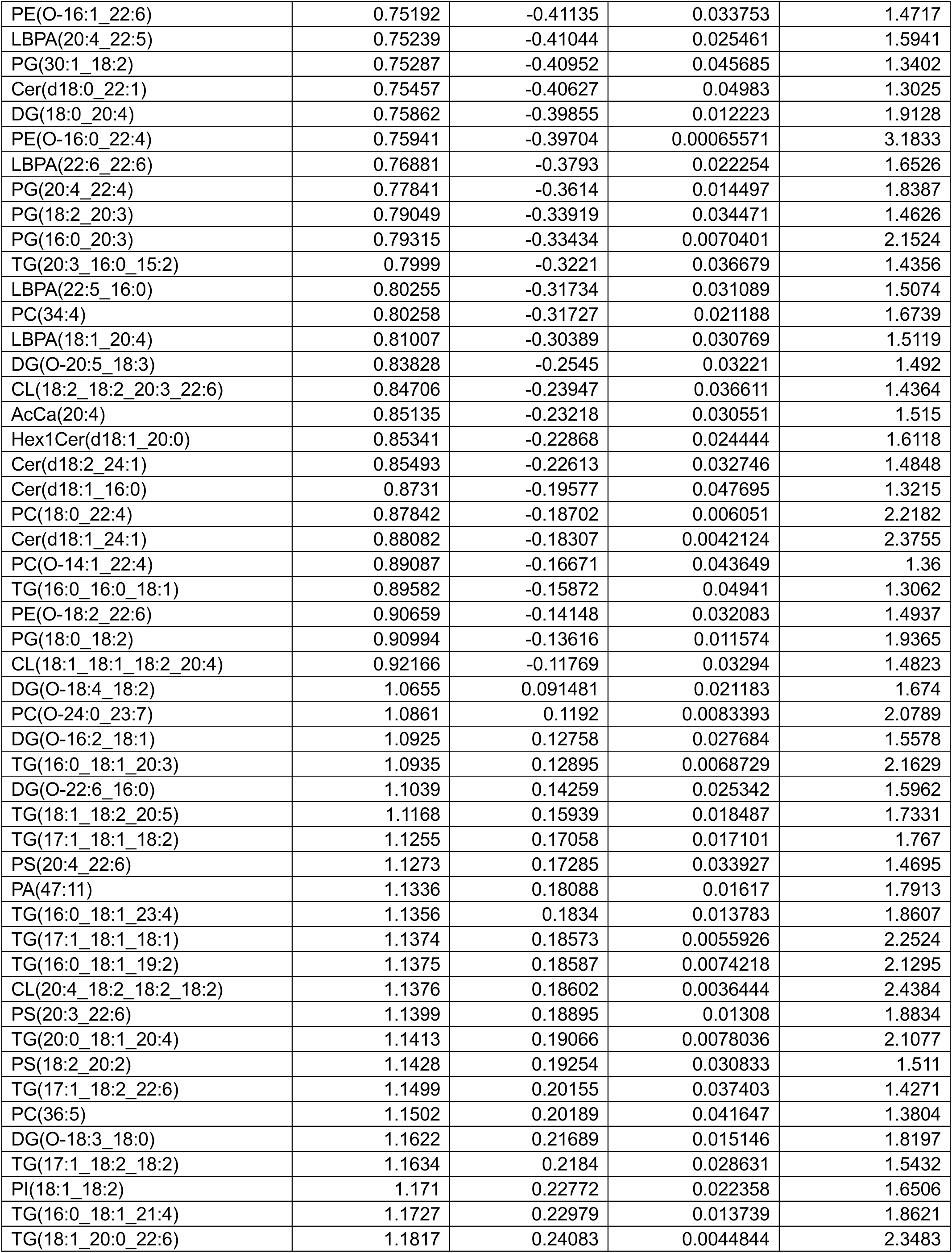
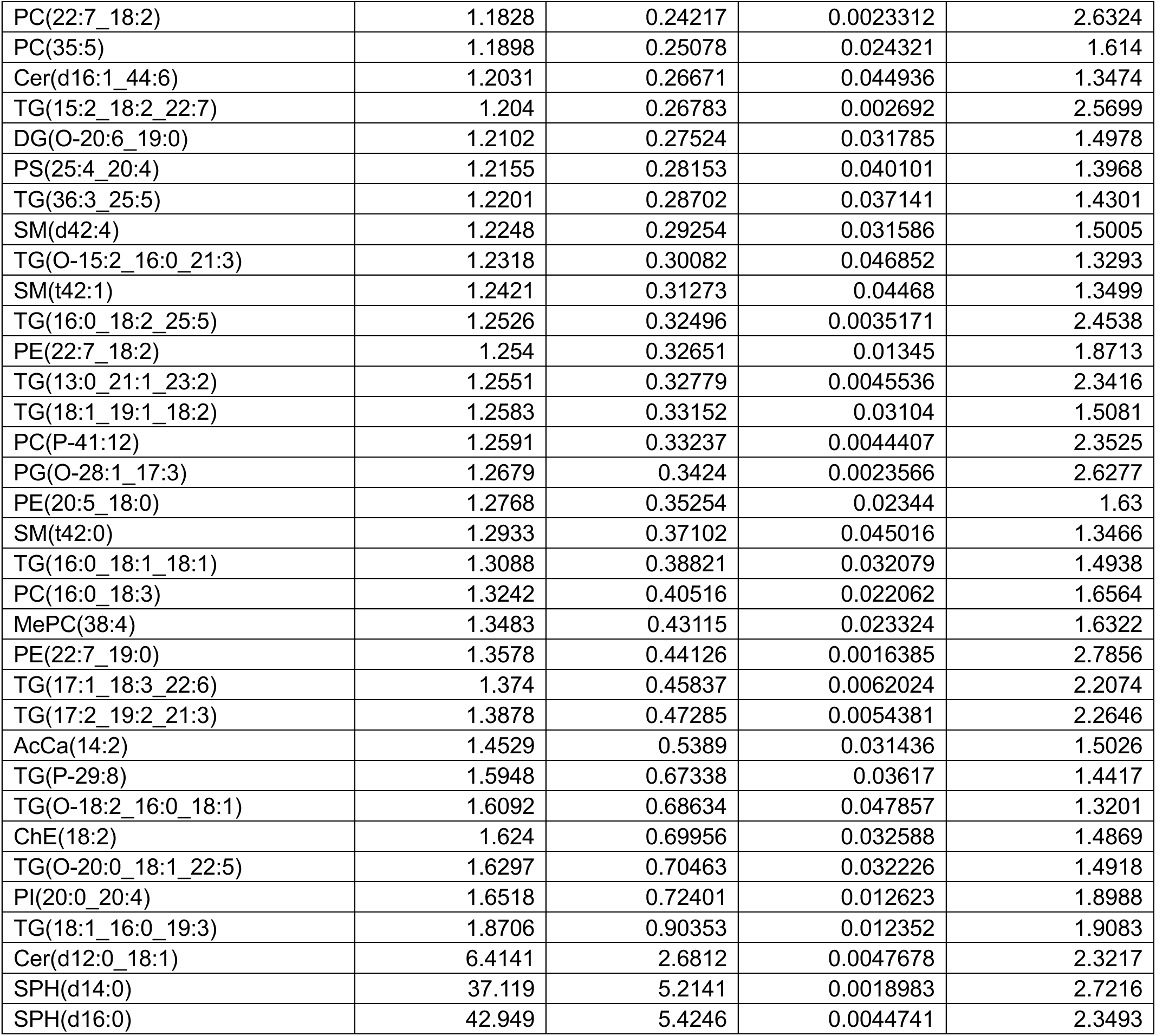
Differential liver lipid profiles between control and *Slc25a35* AAV-KO^Liver^ mice (P-value < 0.05), related to Figure 6.

**Table S6.**
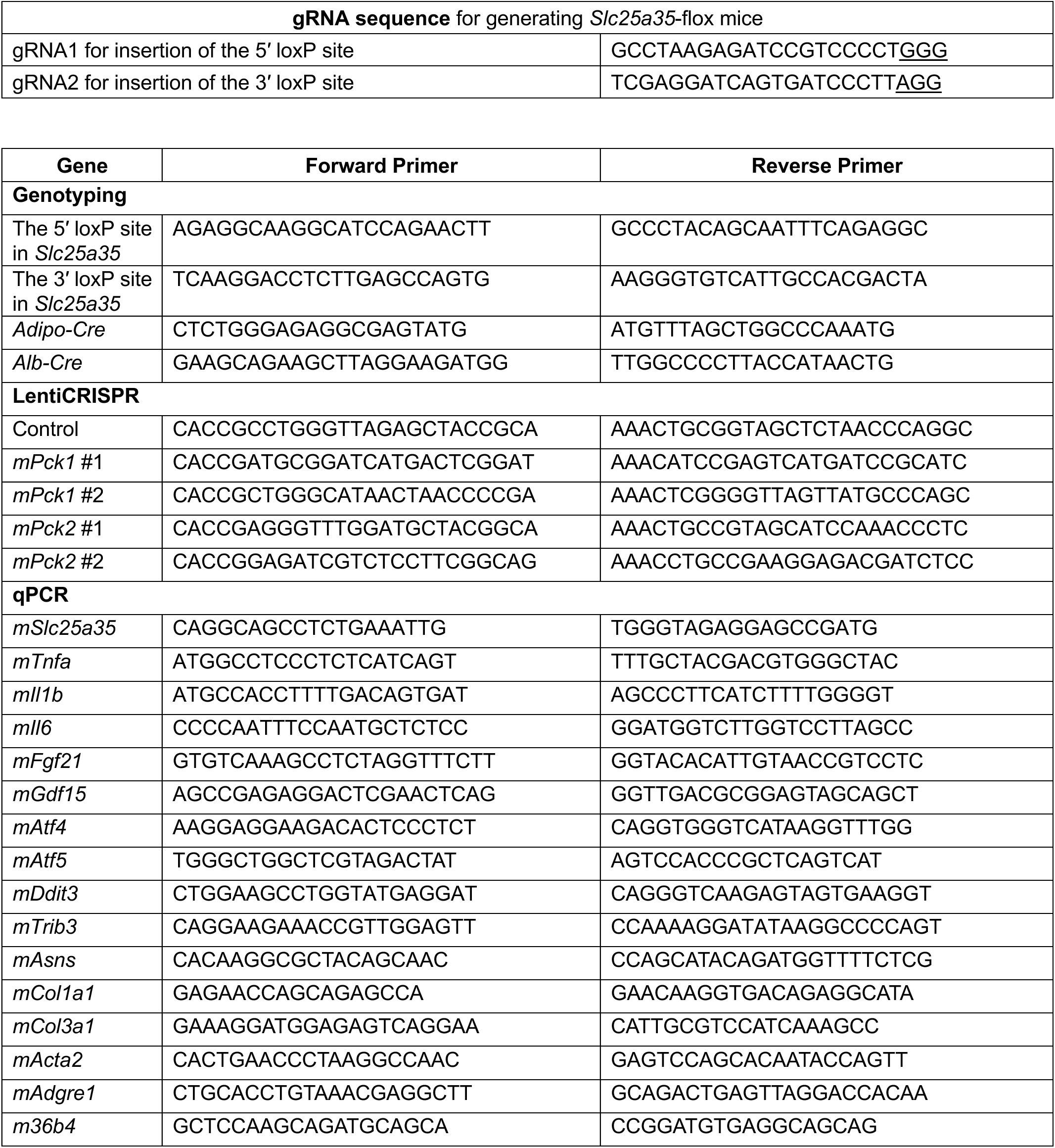
Primer sequences used for gRNA, genotyping, LentiCRISPR, and qPCR, related to STAR Methods.

